# No genetic evidence yet for hinnies at Mazongshan (400-160 BCE), northwestern China

**DOI:** 10.64898/2026.03.27.714239

**Authors:** Gaétan Tressières, Hojjat Asadollahpour Nanaei, Xuexue Liu, Yanli Zhang, Ludovic Orlando

## Abstract

In their recent study entitled “Ancient DNA reveals the co-existence of domestic horses, donkeys and their hybrids in the prehistorical northwestern China”, Li and colleagues (2026) report the genetic identification of three horses, three donkeys and four first-generation hinny hybrids dating to 400–160 BCE from the Mazongshan jade mining site in northwestern China. While a re-analysis of their ancient DNA sequence data confirms the horse and donkey identifications, it indicates that the four putative hinny specimens were, in fact, donkeys. This revision removes the primary evidence originally shown for the presence of hinnies at this site. Therefore, new data from the Mazongshan bone assemblage are required to support the proposed role of hinny hybrids as integral components of trans-regional trade networks during the Late Warring States and Early Han periods.

---

Li and colleagues (2026) recently reported ancient DNA analyses of ten ancient equine specimens (Table S1) excavated from the Mazongshan jade mining site, Northwestern China, radiocarbon dated to 400-160 BCE. The authors identified three horses, three donkeys, and most notably four hinny specimens, purportedly representing first-generation offspring sired by horse stallions and donkey jennets. This claim is remarkable for two key reasons.

Previous ancient DNA studies suggested that hinnies are exceedingly rare in ancient bone assemblages. For instance, in present-day France, genetic analyses of 874 equine specimens spanning 700 BCE–1950 CE identified only a single hinny (Lepetz et al. 2021). Another specimen could be added when extending the analysis to Byzantine Türkyie (Clavel et al. 2021). By contrast, mules–the reciprocal hybrid, produced by mating donkey jacks with horse mares–are more frequent in the same assemblages (8.90% versus 0.21%). This pattern aligns with evidence from experimentally manipulated crossings, showing stark asymmetry in pregnancy rates following reciprocal matings between donkeys and horses, with 54.0% success at day 30 for mule compared to just 8.5% for hinny (Jónsson et al. 2014a).

The 40% hinny prevalence reported at Mazongshan would, thus, imply highly unusual breeding practices within this Late Warring States–Early Han archaeological context, intentionally prioritizing the most difficult and least economical production of hinny hybrids. Such a scenario challenges established understanding of equine husbandry and warrants scrutiny of the genetic evidence underpinning these identifications.

The mapDamage analyses (Jónsson et al. 2013) presented by Li and colleagues (2026) in their Figure 2 raises potential methodological concerns. The size distribution of equine DNA templates exhibits multiple peaks, suggestive of either atypical DNA fragmentation after death or mapping artifacts. Notably, these peaks align with the default edit distance thresholds (38, 64 and 93 bp) tolerated by the BWA aligner (Li & Durbin 2010).

Furthermore, the absence of an excess of purines at the reference position preceding read starts indicates that DNA fragmentation was not driven by DNA depurination, the primary mechanism of post-mortem DNA fragmentation (Dabney et al. 2013). The expected cytosine deamination signature–a hallmark of ancient DNA–is also apparently missing. Instead, the first and last four positions of the endogenous DNA templates show an excess of soft-clipped bases and non-canonical mutation types, deviating from the standard C→T and G→A transitions (Jonsson et al. 2013). Combined, these profiles contrast with the typical ancient DNA degradation patterns yet recovered by the authors when testing previously published ancient DNA data as positive controls (Gaunitz et al. 2018, Fages et al. 2019, Librado et al. 2021, Todd et al. 2022).

The unexpected identification of four hinnies, coupled with the atypical post-mortem DNA damage profiles, prompted us to re-examine the original sequence data. We retrieved raw FASTQ files from the National Genomics Data Centre repository (CRA027067), comprising 426.01–453.03 million sequence pairs per specimen. After demultiplexing based on the two most common dual indexes, 256.98– 315.14 million sequence pairs remained for further processing with Paleomix v1.2.13 (Schubert et al. 2014). Data processing included adapter removal and collapsing of overlapping sequence pairs with Adapter Removal v2.3.0 (Schubert et al. 2016), and alignment to the EquCab2 horse reference genome (Wade et al. 2009) using Bowtie2 v2.3.5.1 (Langmead et al. 2012), with parameters optimized for ancient DNA (Poullet & Orlando 2020). Read alignment revealed a modest fraction of endogenous DNA across all ten samples, with high-quality unique alignments accounting for only 2.27×10^-6^–2.11×10^-4^ of the data, yielding an average depth-of-coverage of 5.03×10^-5^–7.96×10^-3^-fold (Fig. 1).

**Fig. 1.**
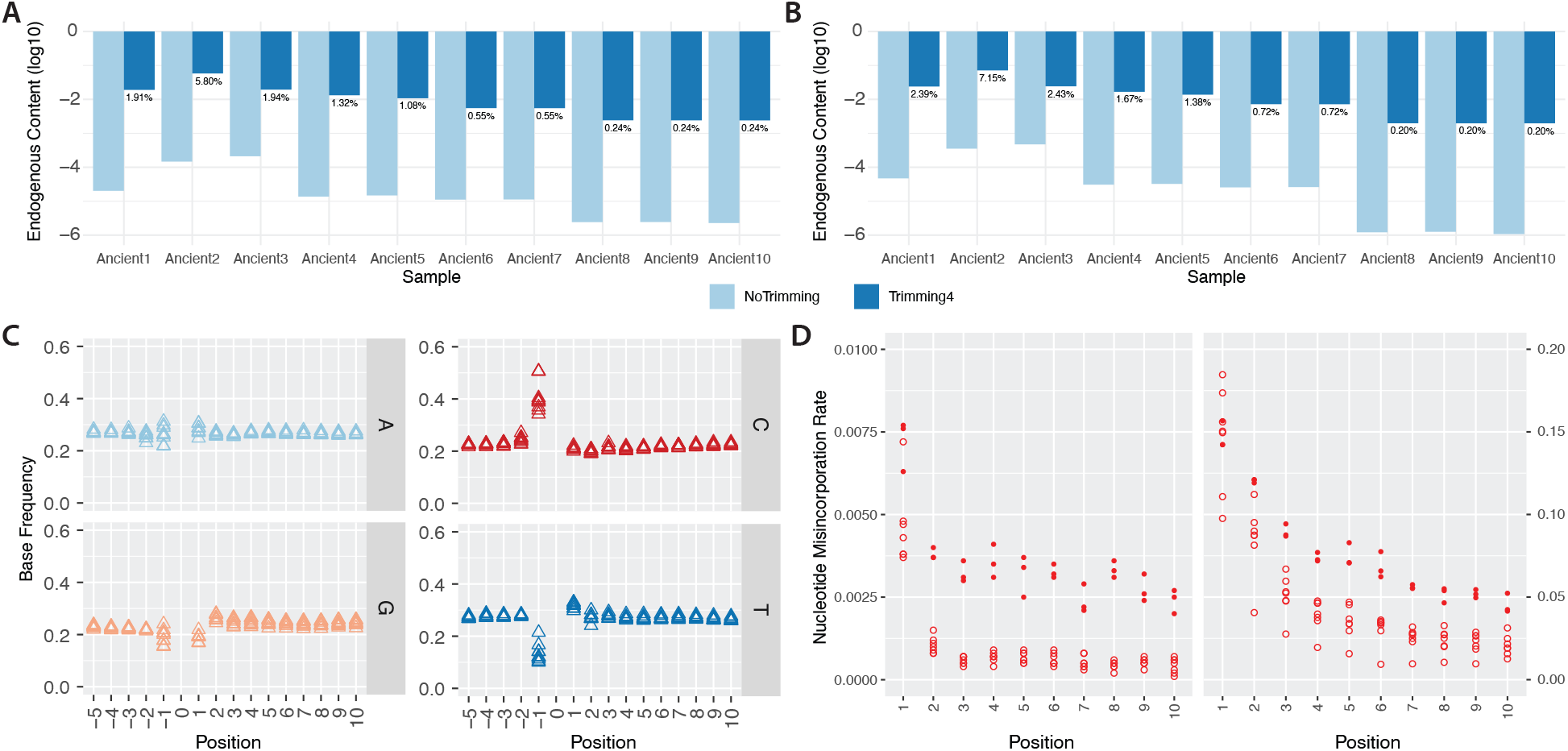
Endogenous DNA content and DNA decay signatures. (A) Endogenous DNA content estimates. Read alignment was carried out against the EquCab2 nuclear genome and the NC_001640 horse mitogenomes. BAM files were filtered for PCR duplicates, low-quality (MQ<25) or too short (<25 bp) alignments. In the ‘NoTrimming’ condition, demultiplexed collapsed and uncollapsed reads were used for mapping. In the ‘Trimming4’ condition, demultiplexed collapsed and uncollapsed reads were trimmed for the four bases of each DNA template. Percentages indicate the endogenous content estimate post-trimming. (B) Same as (A), excepting that the donkey ASM1607732v2 nuclear assembly and the NC_001788 mitogenome were used for mapping. (C) mapDamage (Jónsson et al. 2013) base composition profile around read starts, with -1 representing the position in the nuclear reference genome immediately preceding read starts and +1–+10 the first read positions. The ASM1607732v2 assembly was used as the reference genome. (D) PMDtools (Skoglund et al. 2014) C → T mis-incorporation rates along the first 10 read positions within CpG contexts (right), or across all contexts (left). The three horse specimens (filled circles) show inflated misincorporation rates due to the more distant reference genome selected for mapping.

**Fig. 2.**
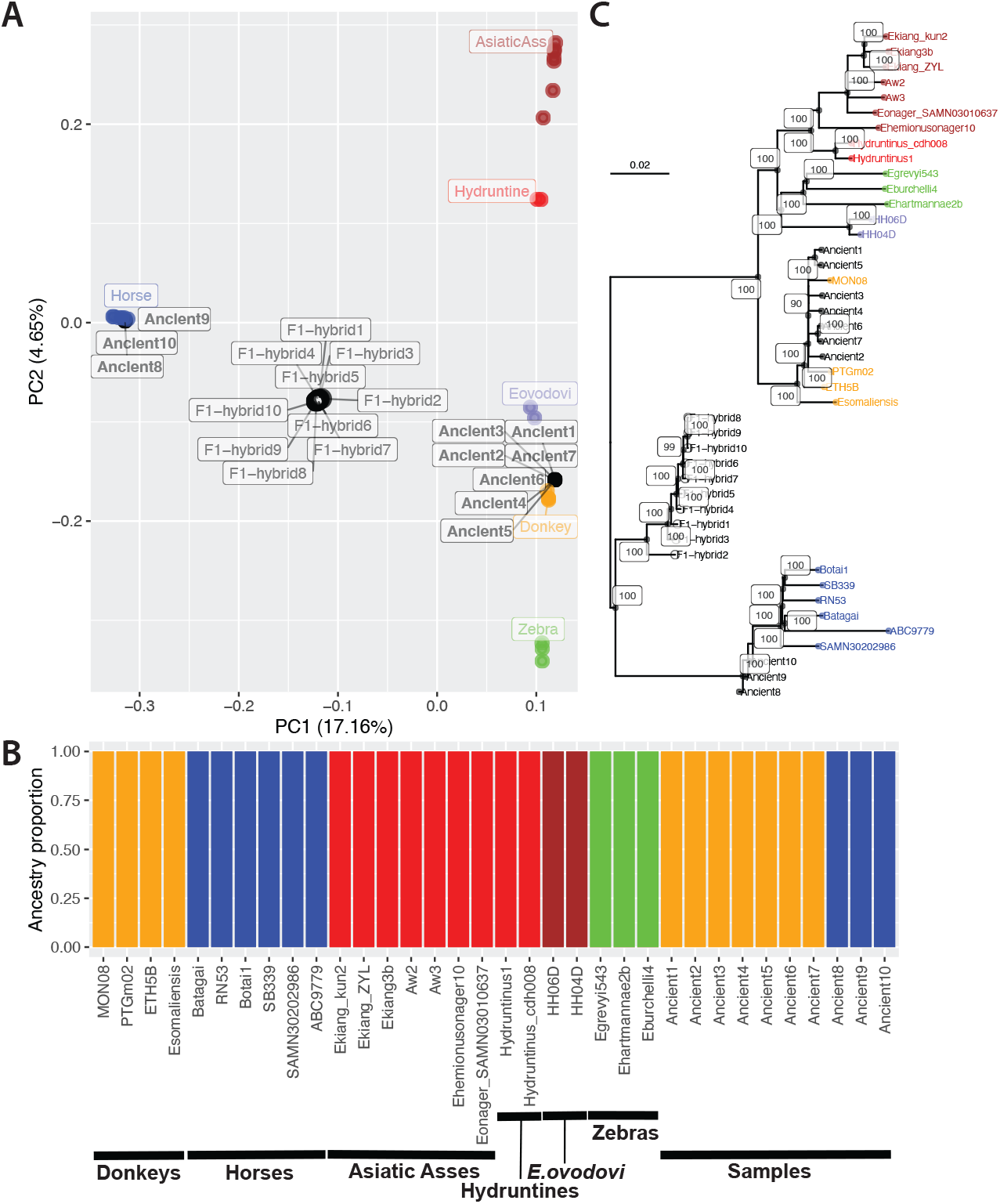
smartPCA (Patterson et al. 2006), ADMIXTURE (Alexander et al. 2009), and Neighbor-Joining (Lefort et al. 2015) phylogenetic reconstructions. (A) Principal Components 1 and 2. The ten ancient specimens (Ancient1–Ancient10, filled circles) and the ten simulated first-generation hybrids (F1-hybrid1–F1-hybrid10, open circles) were projected against the PCA space formed by the 24 high-quality genomes present in our extended panel (filled colored circles). The fraction of the genetic variance explained by each Principal Component is shown between parentheses. (B) Supervised ADMIXTURE (Alexander et al. 2009) profiles for K=3 predefined genetic ancestries (Horses, Zebras, and Asses). (C) Neighbor Joining tree. Node supports are based on 100 bootstrap pseudo-replicates.

However, nearly all high-quality alignments started with the GTCT motif, and ended with its reverse complement (AGAC). This pattern was consistent with the base compositional profiles shown by the authors across all ten libraries. It suggests an artifact introduced by the wet-lab methodology used, probably the addition of a unique four-base tag identifier. Since these exogenous bases likely interfered with mapping, we trimmed the collapsed demultiplexed reads by four bases at both termini. We also trimmed the uncollapsed demutiplexed pairs for the first four bases of each read forming a pair. Remapping substantially improved endogenous DNA content (0.24–5.80%) (Fig. 1A) and average depth-of-coverage (0.03-0.76-fold) in all ten ancient specimens. Mitochondrial coverage also increased dramatically when aligning to the horse (NC_001640, 3.11–45.92-fold versus 0.00–0.08-fold), and donkey (NC_001788, 1.22–214.84-fold versus 0.00–1.02-fold) reference mitogenomes.

The revised mapDamage (Jónsson et al. 2013) base composition profiles showed an excess of cytosine residues at the reference position immediately preceding read starts (Fig. 1C), consistent USER-treatment prior to library construction (Orlando et al. 2021). This was further supported by PMDtools (Skoglund et al. 2014) substitution profiles based on 100,000 random alignments against the longest chromosome (NC_052178.1), which showed declining C→T misincorporation rates from read starts in CpG contexts (Fig. 1D). Beyond CpG dinucleotides, these rates remained relatively flat along all alignment positions outside CpG contexts, except on the first position, where the efficacy of USER treatment is reduced (Rohland et al. 2015) (Fig. 1D). Combined, these analyses confirmed the ancient DNA data from Li and colleagues (2026) as authentic.

Processing read alignments with Zonkey (Schubert et al. 2017) confirmed the reported molecular sex for six males (Ancient1, Ancient3, Ancient5–Ancient8) and two females (Ancient2 and Ancient4), and identified two undetermined specimens as males (Ancient9 and Ancient10). These analyses also validated initial taxonomic classification of three horses (Ancient8–Ancient10), and three donkeys (Ancient1–Ancient3) (Supplementary files S1–S10). However, the four specimens initially identified as hinnies (Ancient4–Ancient7) were reclassified as donkeys based on Principal Component Analysis (PCA) clustering patterns within the comparative panel (Fig. S1A and Fig. S1B), and ADMIXTURE v1.3.0 (Alexander et al. 2009) ancestry profiles showing no zebra or caballine contribution (Fig. S1C).

To further evaluate the taxonomic status of all specimens, and to assess potential contributions from equine species beyond those considered in Zonkey (Schubert et al. 2017), we leveraged a subset (N=24) of a larger comparative panel published by Pan and colleagues (2024) (Table S2). This subset comprises representatives of multiple equine species, including *Equus ovodovi*, known in China until at least ∼1,300 BCE (Cai et al. 2022). Following alignment against the donkey reference genome from Wang and colleagues (2020) (GCA_016077325.2), we identified a total of N=15,889,005 pseudo-haploid transversion sites in at least half of the specimens.

The ten ancient specimens were then projected onto the PCA space derived from the genetic variation present in the comparative panel (Fig. 2A). PCA LSQ projections were carried out using smartPCA v16000 (Patterson et al. 2006), turning the auto-shrink and inbred options on, to compensate for coverage variation (0.02–0.96-fold; Fig. 1B) in ancient specimens. While six specimens clustered as expected with their respective species, the four putative hinny specimens grouped again with donkeys.

To validate this placement, we simulated hinny alignments by equally balancing sequence data from one modern horse (SAMN30202986) and one modern donkey (SAMEA110140444), targeting the average depth-of-coverage of each ancient genome. As expected, these simulated hinnies (F1-hybrid1 to F1-hybrid10) projected to intermediate projections between the horse and donkey clusters (Fig. 2A, Fig. S2). Supervised ADMIXTURE (Alexander et al. 2009) on 1,881,157 unlinked transversion SNPs further revealed that the four alleged hinny specimens exhibited >99% donkey genetic ancestry (Fig. 2B). Finally, Neighbor-Joining phylogenetic reconstruction showed that the three ancient horses clustered within a monophyletic group including the other horses from the panel, while all remaining specimens grouped together with donkeys (Fig. 2C). These placements were supported by maximal (100%) bootstrap values. Combined with the intermediate placement of the simulated hinnies between caballine and non-caballine clusters, our analyses demonstrate that the four specimens originally identified as hinnies (Ancient4–Ancient7) were, in fact, donkeys.

Notwithstanding the misidentification of the four donkey specimens as hinnies, the sequence data from Li and colleagues (2026) remain invaluable for elucidating the history of equine husbandry in eastern Asia. Their study provides the earliest genetic evidence for the presence of donkeys in China, a significant milestone in our understanding of ancient animal management in the region. The co-existence of donkeys and horses on site underscores integrated management practices of at least two equine species providing complementary traits to cover a range of activities useful for humans. This apparently contrasts with equine management in Europe primarily focused on horses at the time (Lepetz et al. 2021). While none of the ten Mazongshan specimens comprised hybrids, this apparent absence does not preclude the possibility that hybrids were bred locally at the time. Statistically, with the current sample size, mules or hinnies could still have represented up to 25.89% of the bone assemblage (one-sided Binomial test, *p*-value=9.77×10^-4^). To confidently assert that hybrids accounted for no more than 5% (or 1%) of the equine population at the site, analyses of 59 (or 299) specimens would be required.

## Supporting information

Supplementary File 10. Zonkey (Schubert et al. 2017) report for donkey specimen Ancient10.

Supplementary File 9. Zonkey (Schubert et al. 2017) report for donkey specimen Ancient9.

Supplementary File 8. Zonkey (Schubert et al. 2017) report for donkey specimen Ancient8.

Supplementary File 7. Zonkey (Schubert et al. 2017) report for donkey specimen Ancient7.

Supplementary File 6. Zonkey (Schubert et al. 2017) report for donkey specimen Ancient6.

Supplementary File 5. Zonkey (Schubert et al. 2017) report for donkey specimen Ancient5.

Supplementary File 4. Zonkey (Schubert et al. 2017) report for donkey specimen Ancient4.

Supplementary File 3. Zonkey (Schubert et al. 2017) report for donkey specimen Ancient3.

Supplementary File 2. Zonkey (Schubert et al. 2017) report for donkey specimen Ancient2.

Supplementary File 1. Zonkey (Schubert et al. 2017) report for donkey specimen Ancient1.

Table S2. Originally from Table S1 in Pan et al. (2024) with the 24 specimens used in our study.

Table S1. List of Ancient to Ancient10 published by Li and colleagues in Supplementary Table 1.

---

**Fig. S1.**
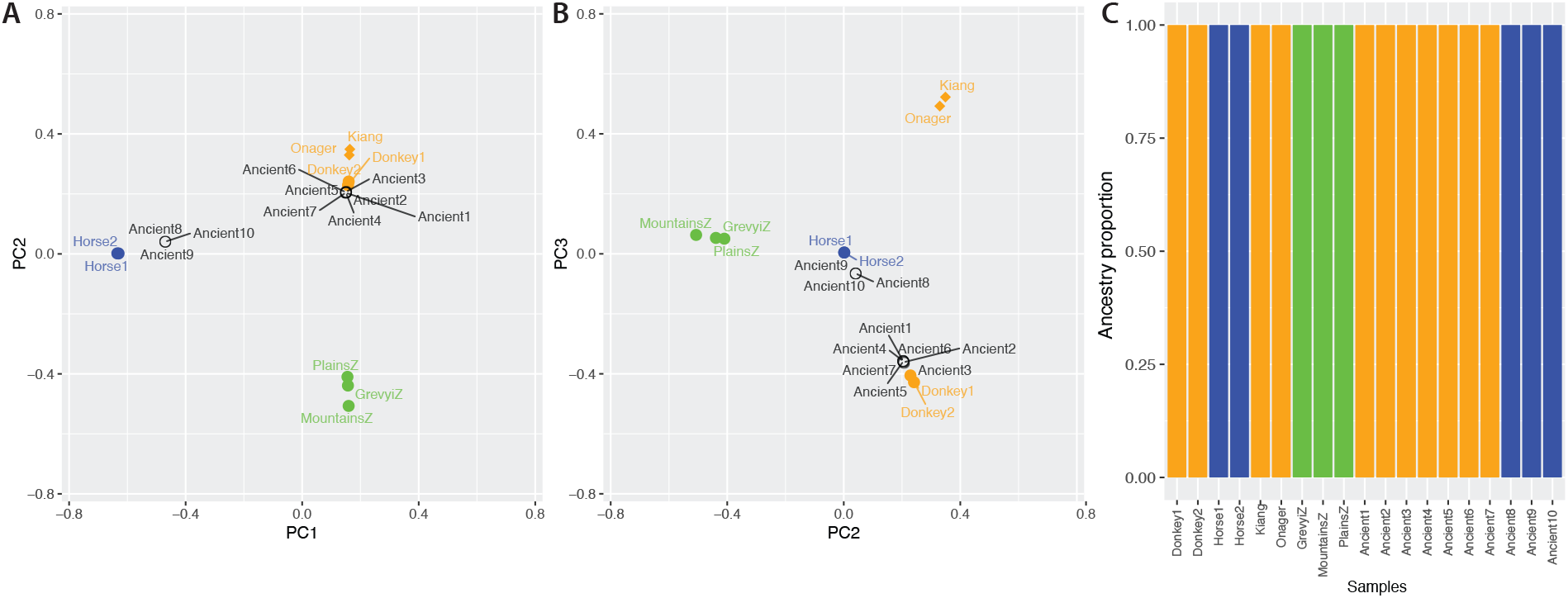
Zonkey (Schubert et al. 2017) PCA and ADMIXTURE analyses. (A) PCA for first and second Principal Components (PCs). A total of 10 analyses were carried out, each including the Zonkey comparative panel plus one ancient specimen. We used Procrustes factor rotation from the EFA.dimensions (O’Connor 2026) to co-visualize the 10 individual analyses within a single graph. GrevyiZ, MountainsZ and PlainsZ stand for Grevyi’s, Montains and Plains Zebras. The reference samples from the Zonkey comparative panel are shown in colors. (B) Same as (A) for second and third PCs. (C) Supervised ADMIXTURE (Alexander et al. 2009) profiles for K=3 predefined genetic ancestries (Horses, Zebras, and Asses).

**Fig. S2.**
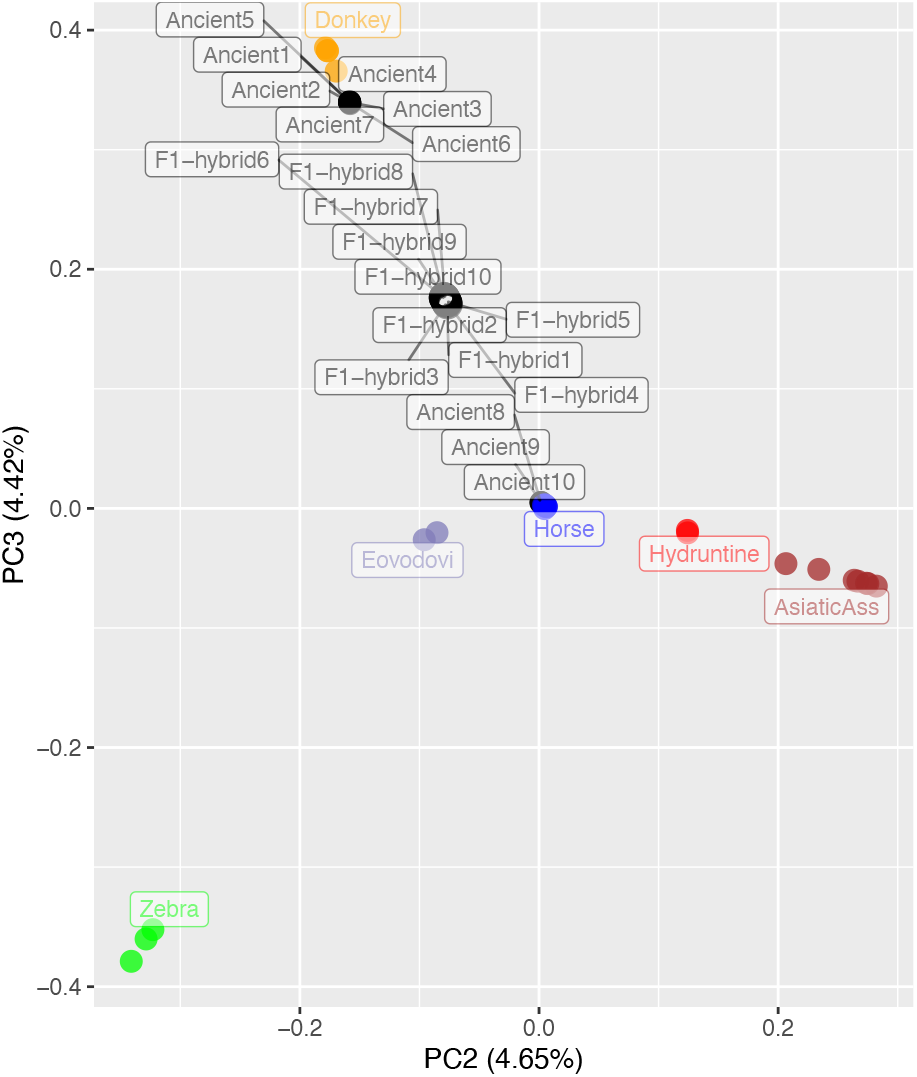
smartPCA (Patterson et al. 2006) projection on Principal Components 2 and 3. The ten ancient specimens (Ancient1–Ancient10, filled circles) and the ten simulated first-generation hybrids (F1-hybrid1–F1-hybrid10, open circles) were projected against the PCA space formed by the 24 high-quality genomes present in our extended panel (filled colored circles). The fraction of the genetic variance explained by each Principal Component is shown between parentheses.

**Table S1. List of Ancient to Ancient10 published by Li and colleagues in Supplementary Table 1**.

**Table S2. Originally from Table S1 in Pan et al. (2024) with the 24 specimens used in our study**.

**Supplementary File 1. Zonkey (Schubert et al. 2017) report for donkey specimen Ancient1**.

The analyses were carried out on the high-quality unique alignments against the horse EquCab2 nuclear reference genome (Wade et al. 2009), supplemented with those against the NC_001640 horse mitogenome.

**Supplementary File 2. Zonkey (Schubert et al. 2017) report for donkey specimen Ancient2**.

**Supplementary File 3. Zonkey (Schubert et al. 2017) report for donkey specimen Ancient3**.

**Supplementary File 4. Zonkey (Schubert et al. 2017) report for donkey specimen Ancient4**.

**Supplementary File 5. Zonkey (Schubert et al. 2017) report for donkey specimen Ancient5**.

**Supplementary File 6. Zonkey (Schubert et al. 2017) report for donkey specimen Ancient6**.

**Supplementary File 7. Zonkey (Schubert et al. 2017) report for donkey specimen Ancient7**.

**Supplementary File 8. Zonkey (Schubert et al. 2017) report for donkey specimen Ancient8**.

**Supplementary File 9. Zonkey (Schubert et al. 2017) report for donkey specimen Ancient9**.

**Supplementary File 10. Zonkey (Schubert et al. 2017) report for donkey specimen Ancient10**.

## CRediT authorship contribution statement

**Gaétan Tressières:** Writing – original draft. **Hojjat Asadollahpour Nanaei:** Writing – original draft - Resources. **Xuexue Liu:** Revision – original draft. **Yanli Zhang:** Writing – original draft. **Ludovic Orlando:** Writing – Methodology – Formal analysis – Supervision – Resources.

