## Supplementary File 10. Zonkey (Schubert et al. 2017) report for donkey specimen Ancient10. for "No genetic evidence yet for hinnies at Mazongshan (400-160 BCE), northwestern China": report.html

PALEOMIX Zonkey v1.2.13.2 - db rev. 20161101

### PALEOMIX Zonkey v1.2.13.2 - db rev. 20161101

#### A pipeline for detection of F1 hybrids in equids.

Schubert M, Ermini L, Sarkissian CD, Jónsson H, Ginolhac A,
Schaefer R, Martin MD, Fernández R, Kircher M, McCue M,
Willerslev E, and Orlando L. "**Characterization of ancient and
modern genomes by SNP detection and phylogenomic and metagenomic analysis
using PALEOMIX**". Nat Protoc. 2014 May;9(5):1056-82. doi:10.1038/nprot.2014.063.
Epub 2014 Apr 10. PubMed PMID: 24722405.

### Contents

Top
Introduction
Analysis overview
Reference Panel
Admixture Estimates
PCA Plots
Treemix Analyses
MT Phylogeny
References

### Introduction

The Zonkey Pipeline is a easy-to-use pipeline designed for the
analyses of low-coverage, ancient DNA derived from historical
equid samples, with the purpose of determining the species of
the sample, as well as determining possible hybridization between
horses, zebras, and asses. This is accomplished by comparing one
or more samples aligned against the *Equus caballus* 2.0
reference sequence with a reference panel of modern equids,
including wild and domesticated equids.

  

For more information, please refer to the
the documentation for the Zonkey pipeline
or
the documentation for the PALEOMIX pipeline,
on which the Zonkey pipeline is based.

  

### Analysis overview

Zonkey run using database rev. 20161101. Data processed using
pysam v0.13,
SAMTools v1.17.0
[*Li *et al.* 2009*] and
PLINK v1.7
[*Purcell *et al.* 2007*]; plotting was carried out using
R v3.6.0. Additional
tools listed below.

  

**Nuclear report from '*/disk/rene/data2/SeqGuanghui/Ancient10\_CRR1938516\_trim4.Horse\_nuc\_wY.realigned.q25.bam*'**

|  |  |
| --- | --- |
| Number of reads processed: | 941470 |
| Number of reads overlapping SNPs: | 475827 |
| Number of SNPs used (incl. transitions): | 472418 |
| Number of SNPs used (excl. transitions): | 142364 |

  

###### Mitochondrial report from '*/disk/rene/data2/SeqGuanghui/Ancient10\_CRR1938516\_trim4.Horse\_mt.realigned.q25.bam*'

|  |  |
| --- | --- |
| Reference sequence used: | 5835107Eq\_mito3 |
| Reference sequence length: | 16659 |
| Number of sites covered: | 16256 |
| Percentage of sites covered: | 97.6 |
| Mean coverage per site: | 5.6 |

**Autosomes vs. sex-chromosomes:**

### Reference Panel

| Group(2) | Group(3) | ID | Species | Sex | Sample Name | Publication |
| --- | --- | --- | --- | --- | --- | --- |
| Caballine Horse HCab | *E. caballus* | Male FM1798 | doi:10.1016/j.cub.2015.08.032 | | | |
| HPrz | *E. przewalskii* | Male SB281 | doi:10.1016/j.cub.2015.08.032 | | | |
| NonCaballine Ass AAsi | *E. a. asinus* | Male Willy | doi:10.1038/nature12323 | | | |
| AKia | *E. kiang* | Female KIA | doi:10.1073/pnas.1412627111 | | | |
| AOna | *E. h. onager* | Male ONA | doi:10.1073/pnas.1412627111 | | | |
| ASom | *E. a. somaliensis* | Female SOM | doi:10.1073/pnas.1412627111 | | | |
| Zebra ZBoe | *E. q. boehmi* | Female BOE | doi:10.1073/pnas.1412627111 | | | |
| ZGre | *E. grevyi* | Female GRE | doi:10.1073/pnas.1412627111 | | | |
| ZHar | *E. z. hartmannae* | Female HAR | doi:10.1073/pnas.1412627111 | | | |

### Admixture Estimates

Admixture proportions estimated using
ADMIXTURE
v1.3 *[Alexander *et al.* 2009]*, using default
parameters.

| 2 ancestral groups | 2 ancestral groups, excluding transitions |
| --- | --- |
|  | **No admixture detected.** |

| 3 ancestral groups | 3 ancestral groups, excluding transitions |
| --- | --- |
| **No admixture detected.** | **No admixture detected.** |

### PCA Plots

Principal Component Analysis carried out using SmartPCA v16000,
from the EIGENSOFT
toolkit.

| Including transitions | Excluding transitions |
| --- | --- |

### Treemix Plots

Detection of population mixture using
TreeMix
v1.13 *[Pickrell and Pritchard 2012]*; parameters were -k
0; -global; and supervised estimation using ancestral groups
listed in the Reference Panel.

#### Including transitions

| Edges = 0 | Residuals |
| --- | --- |

Variance explained by model = 0.999990.

| Edges = 1 | Residuals |
| --- | --- |

Variance explained by model = 0.999993.

#### Excluding transitions

| Edges = 0 | Residuals |
| --- | --- |

Variance explained by model = 0.999997.

| Edges = 1 | Residuals |
| --- | --- |

Variance explained by model = 0.999999.

### Mitochondrial Phylogeny

Phylogenetic inference performed using RAxML
v8.2.12 [*Stamatakis 2006*].

This report is based on the PLAIN 1.0 design by
6ix Shooter Media,
Creative Commons license.
