## Supplementary File 8. Zonkey (Schubert et al. 2017) report for donkey specimen Ancient8. for "No genetic evidence yet for hinnies at Mazongshan (400-160 BCE), northwestern China": report.html

PALEOMIX Zonkey v1.2.13.2 - db rev. 20161101

### PALEOMIX Zonkey v1.2.13.2 - db rev. 20161101

#### A pipeline for detection of F1 hybrids in equids.

Schubert M, Ermini L, Sarkissian CD, Jónsson H, Ginolhac A,
Schaefer R, Martin MD, Fernández R, Kircher M, McCue M,
Willerslev E, and Orlando L. "**Characterization of ancient and
modern genomes by SNP detection and phylogenomic and metagenomic analysis
using PALEOMIX**". Nat Protoc. 2014 May;9(5):1056-82. doi:10.1038/nprot.2014.063.
Epub 2014 Apr 10. PubMed PMID: 24722405.

### Contents

Top
Introduction
Analysis overview
Reference Panel
Admixture Estimates
PCA Plots
Treemix Analyses
MT Phylogeny
References

### Introduction

The Zonkey Pipeline is a easy-to-use pipeline designed for the
analyses of low-coverage, ancient DNA derived from historical
equid samples, with the purpose of determining the species of
the sample, as well as determining possible hybridization between
horses, zebras, and asses. This is accomplished by comparing one
or more samples aligned against the *Equus caballus* 2.0
reference sequence with a reference panel of modern equids,
including wild and domesticated equids.

  

For more information, please refer to the
the documentation for the Zonkey pipeline
or
the documentation for the PALEOMIX pipeline,
on which the Zonkey pipeline is based.

  

### Analysis overview

Zonkey run using database rev. 20161101. Data processed using
pysam v0.13,
SAMTools v1.17.0
[*Li *et al.* 2009*] and
PLINK v1.7
[*Purcell *et al.* 2007*]; plotting was carried out using
R v3.6.0. Additional
tools listed below.

  

**Nuclear report from '*/disk/rene/data2/SeqGuanghui/Ancient8\_CRR1938514\_trim4.Horse\_nuc\_wY.realigned.q25.bam*'**

|  |  |
| --- | --- |
| Number of reads processed: | 941609 |
| Number of reads overlapping SNPs: | 476143 |
| Number of SNPs used (incl. transitions): | 472716 |
| Number of SNPs used (excl. transitions): | 142751 |

  

###### Mitochondrial report from '*/disk/rene/data2/SeqGuanghui/Ancient8\_CRR1938514\_trim4.Horse\_mt.realigned.q25.bam*'

|  |  |
| --- | --- |
| Reference sequence used: | 5835107Eq\_mito3 |
| Reference sequence length: | 16659 |
| Number of sites covered: | 16338 |
| Percentage of sites covered: | 98.1 |
| Mean coverage per site: | 6.0 |
