## Supplementary figures and images for "No genetic evidence yet for hinnies at Mazongshan (400-160 BCE), northwestern China"

### coverage.pdf

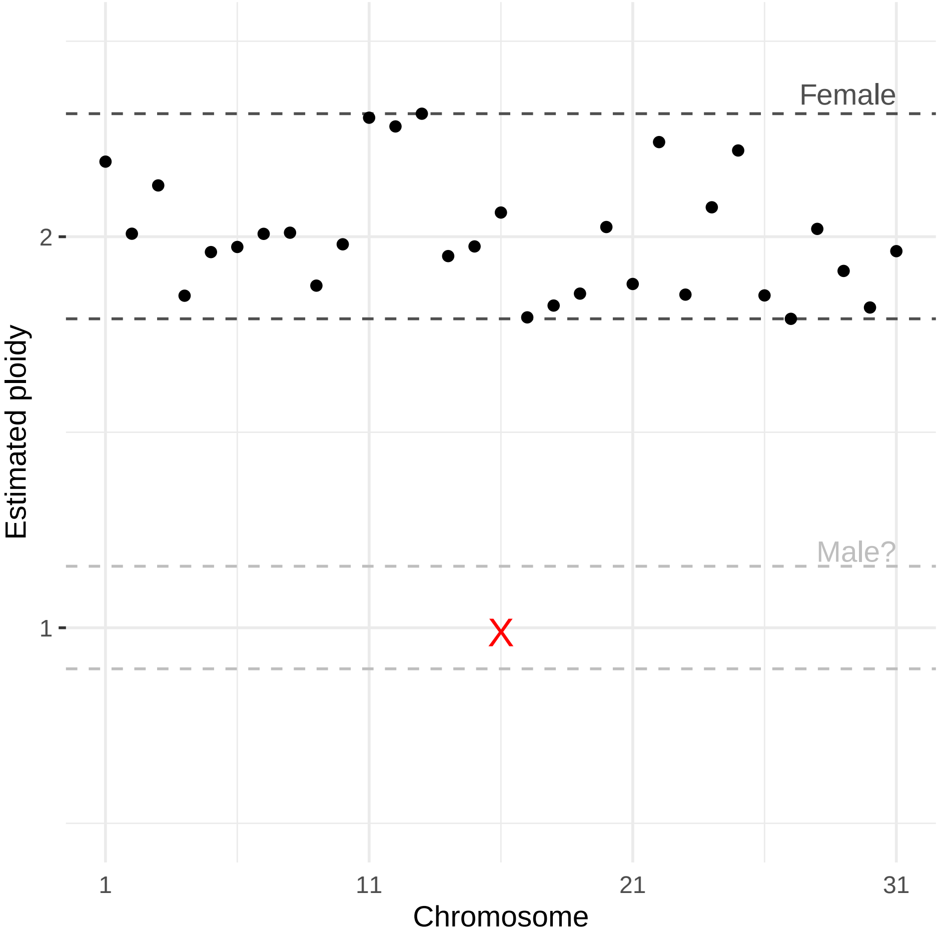

### coverage.pdf

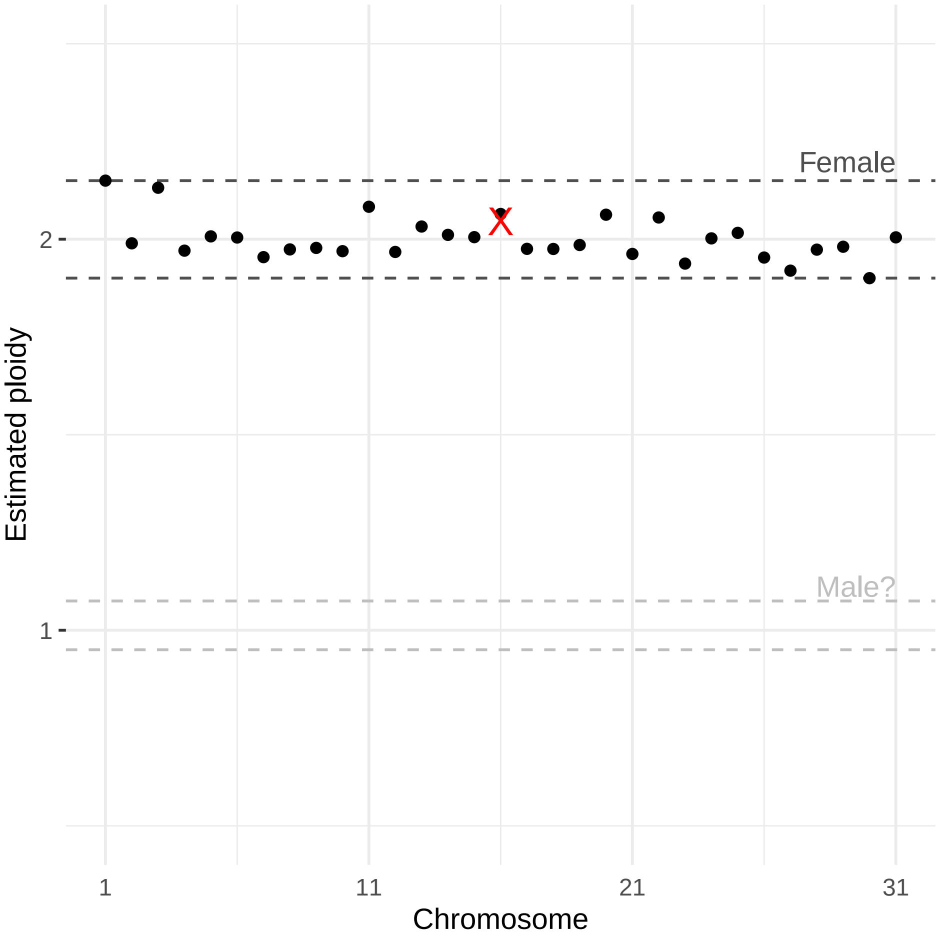

### coverage.pdf

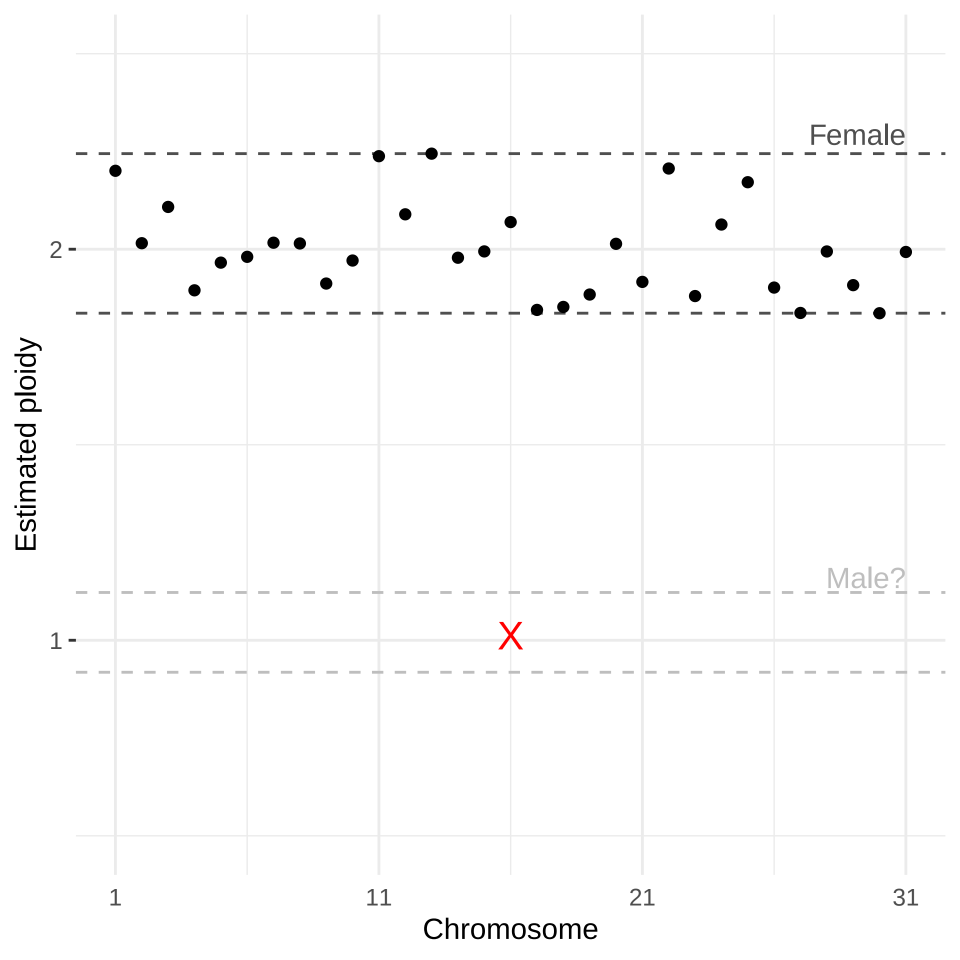

### coverage.png

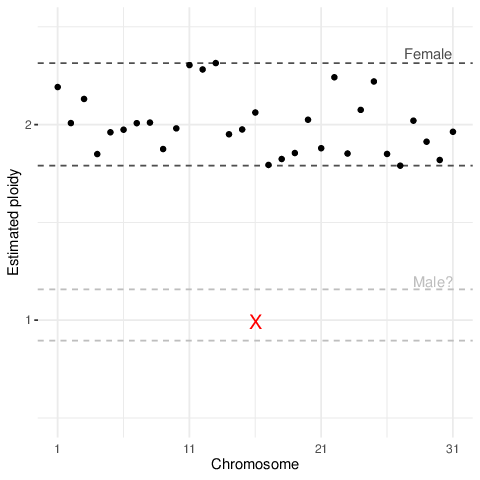

### coverage.png

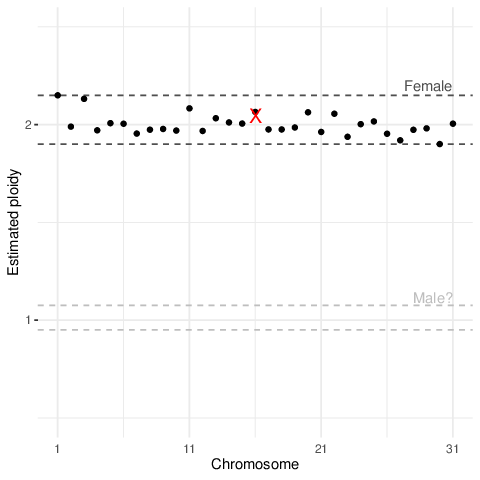

### coverage.png

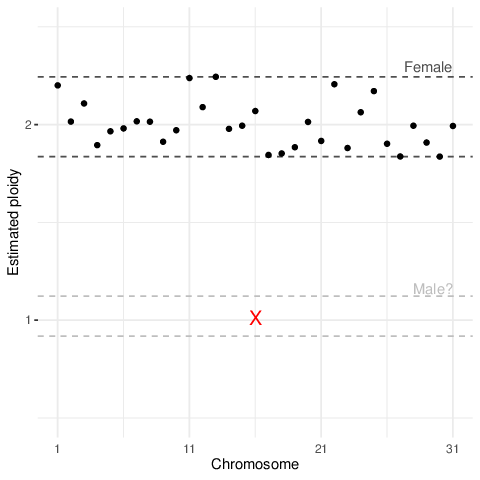

### excl_ts.pdf

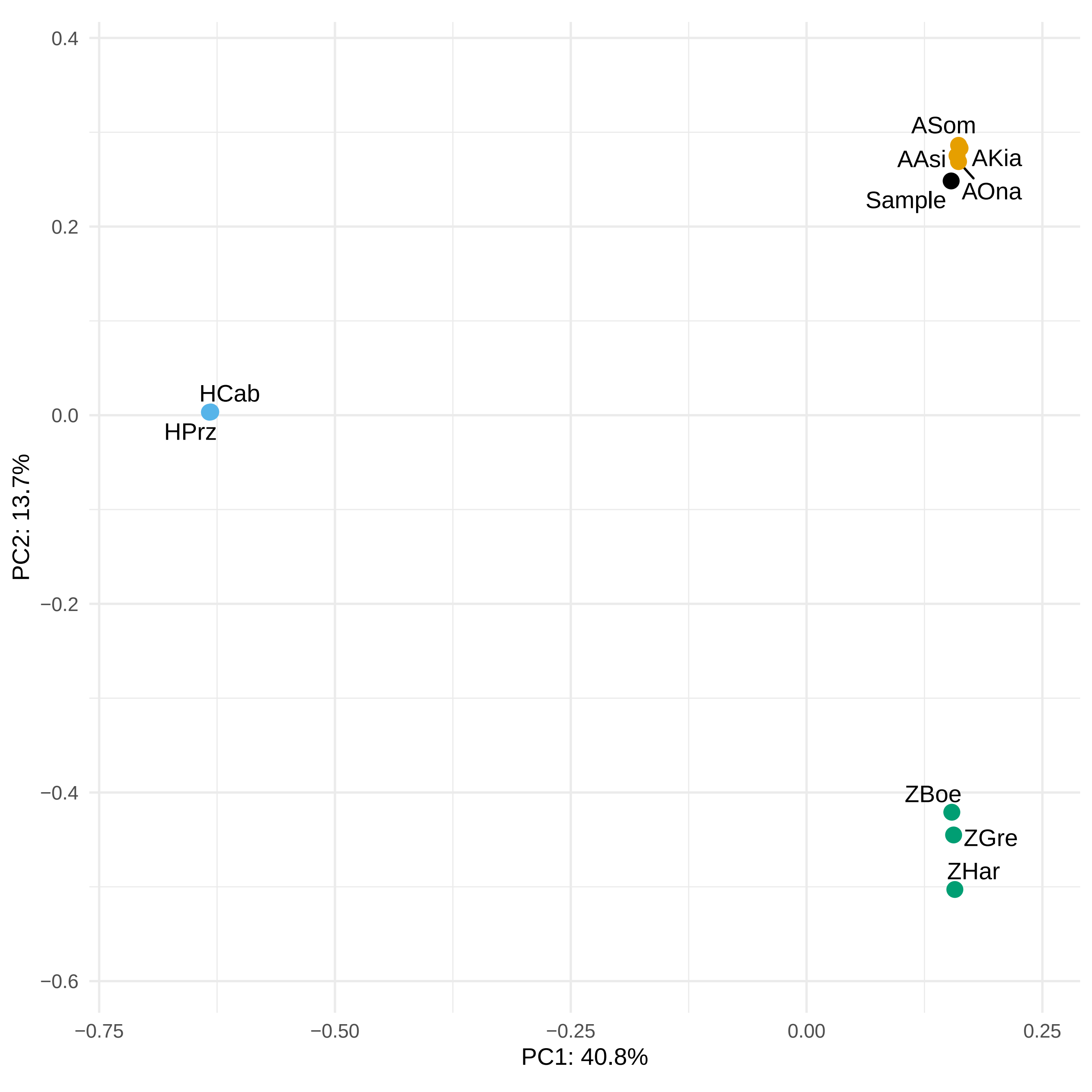

### excl_ts.pdf

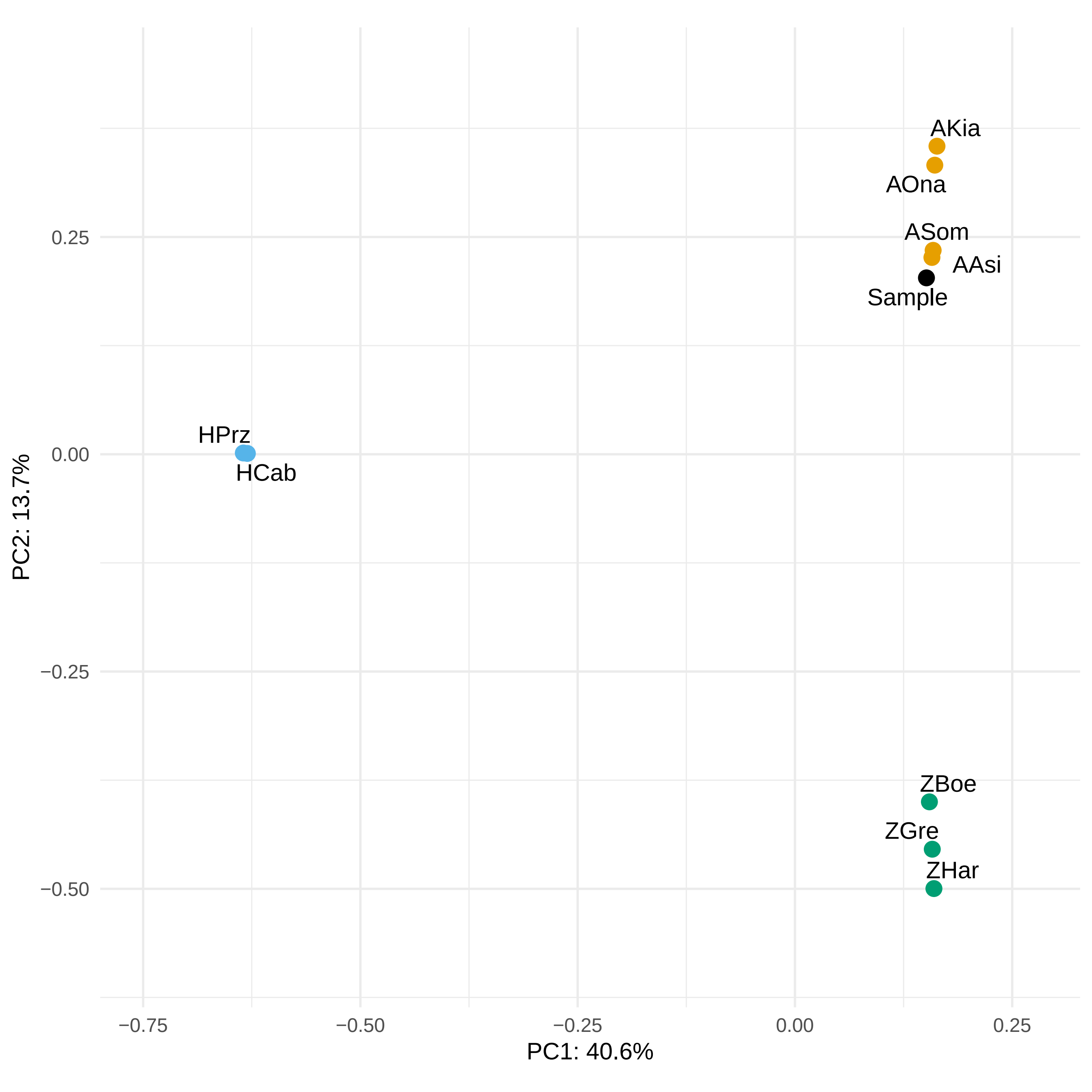

### excl_ts.pdf

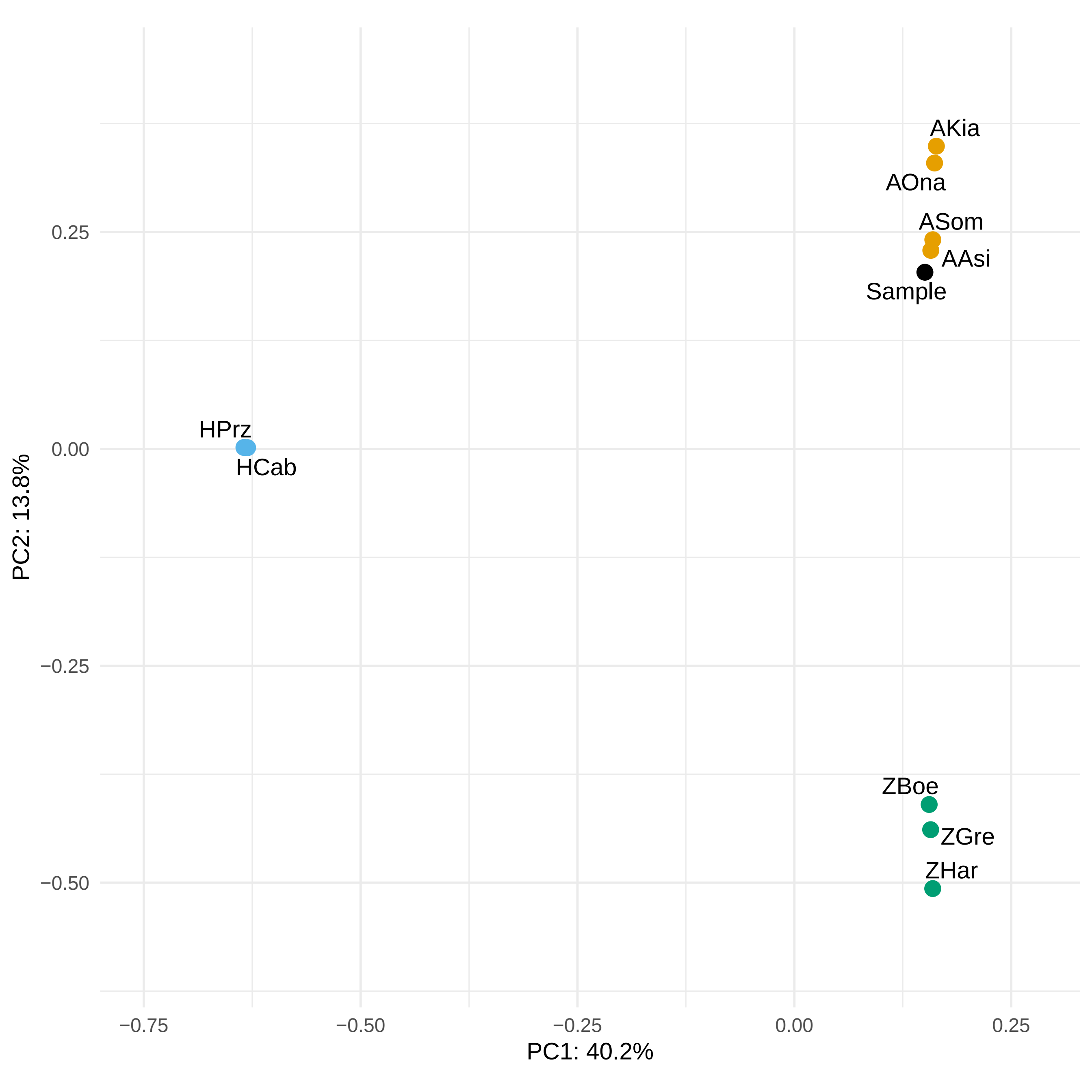

### excl_ts.png

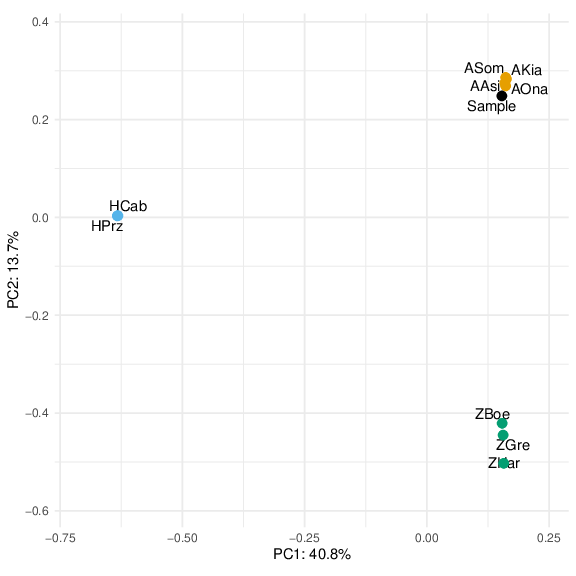

### excl_ts.png

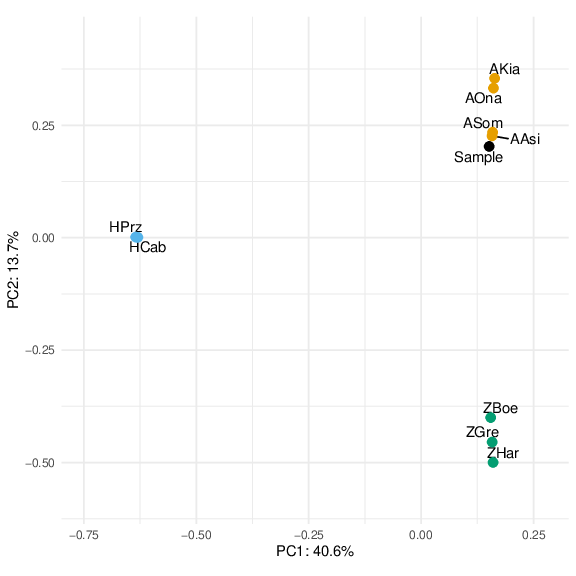

### excl_ts.png

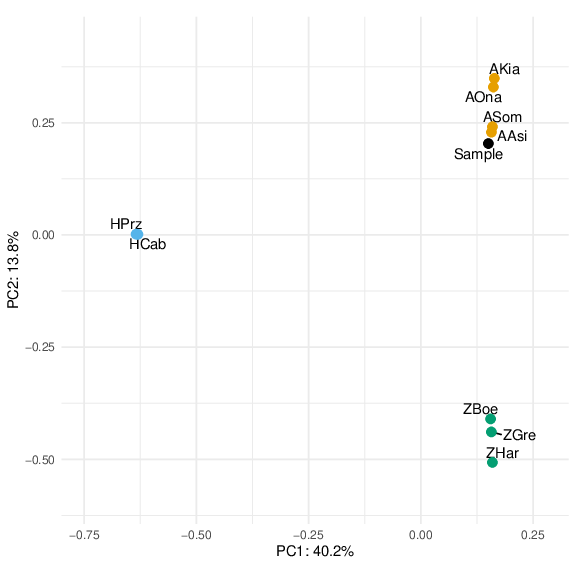

### excl_ts_0_residuals.pdf

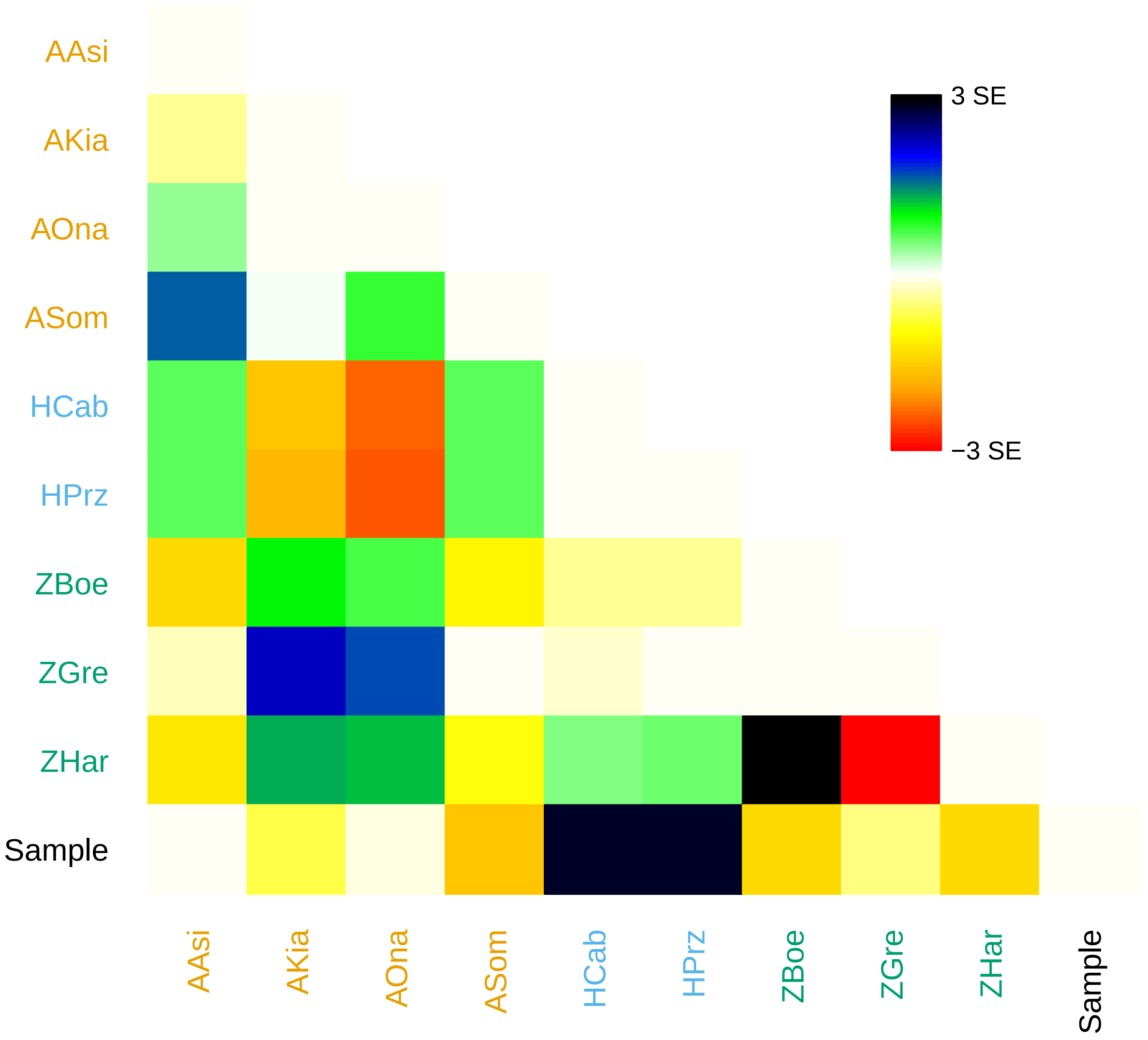

### excl_ts_0_residuals.pdf

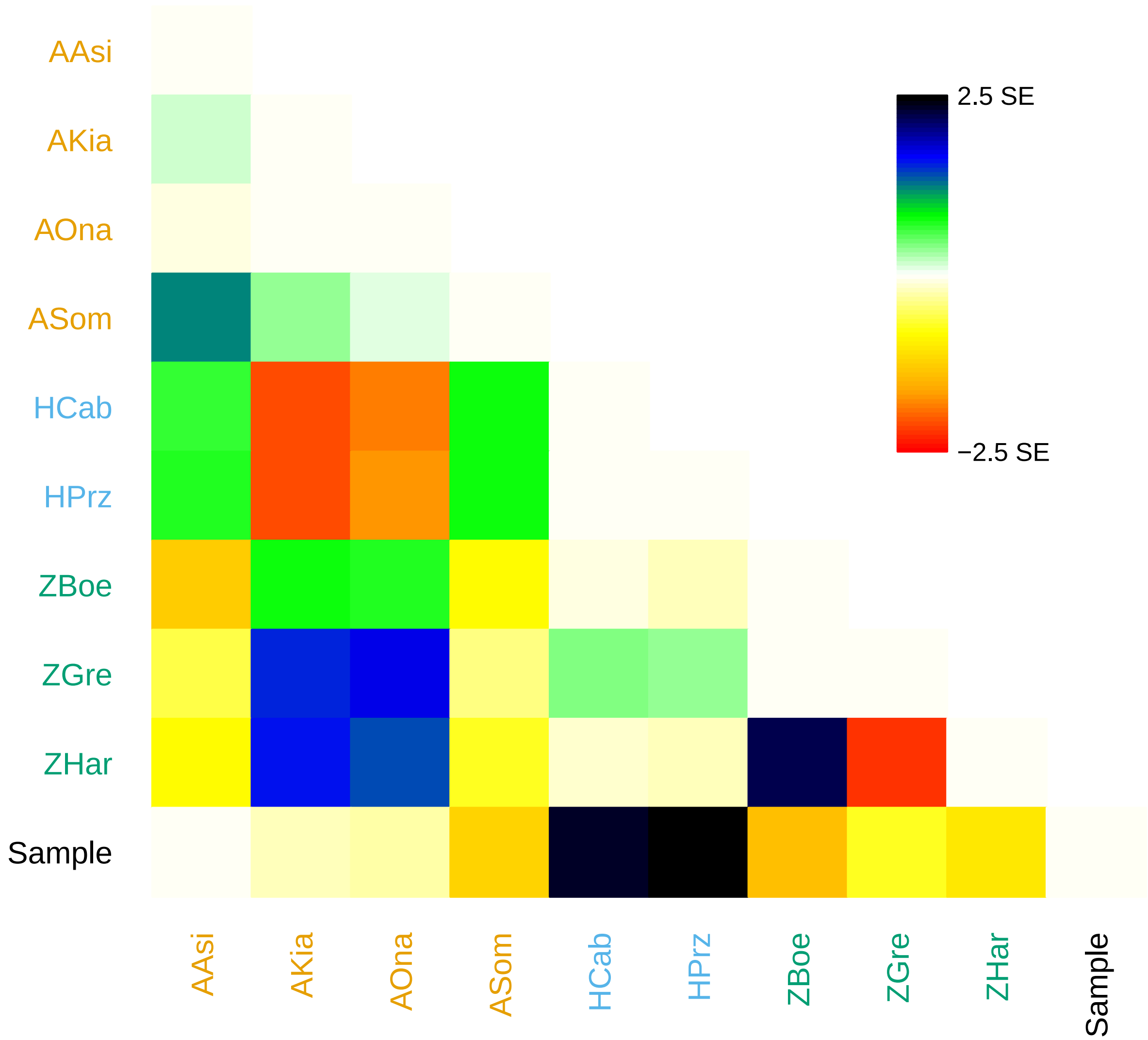

### excl_ts_0_residuals.pdf

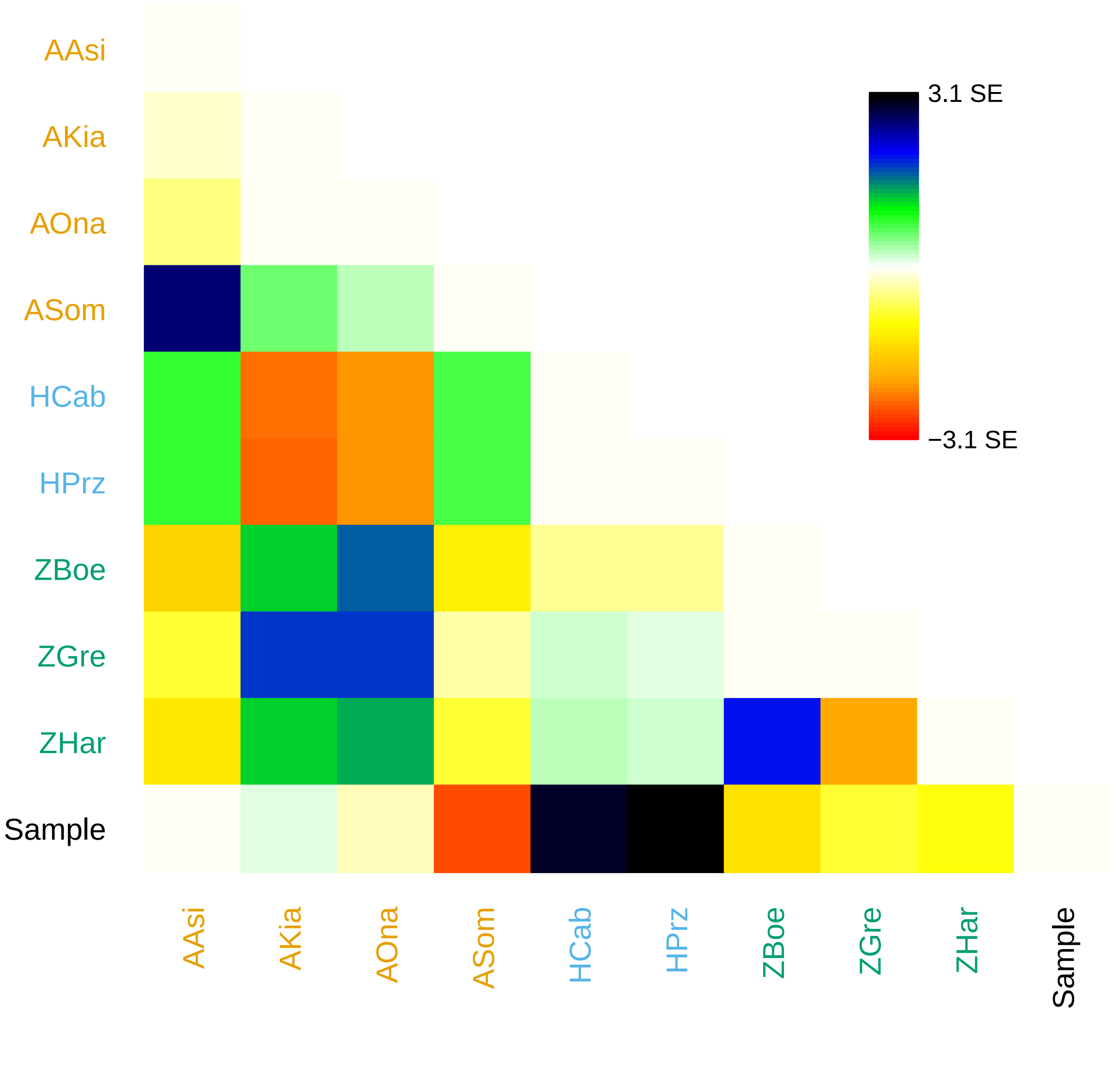

### excl_ts_0_residuals.png

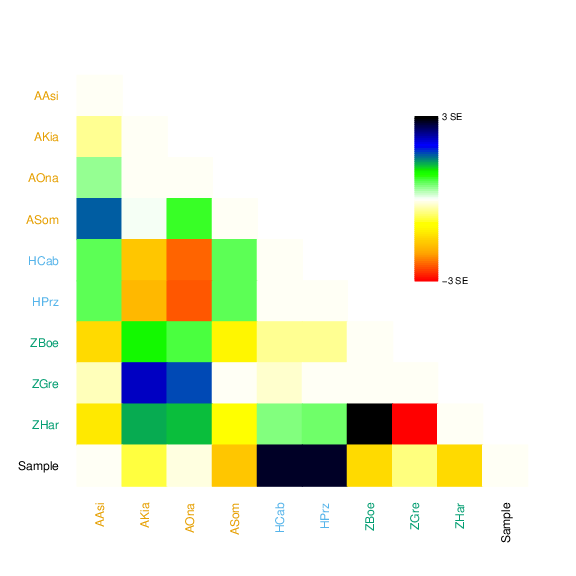

### excl_ts_0_residuals.png

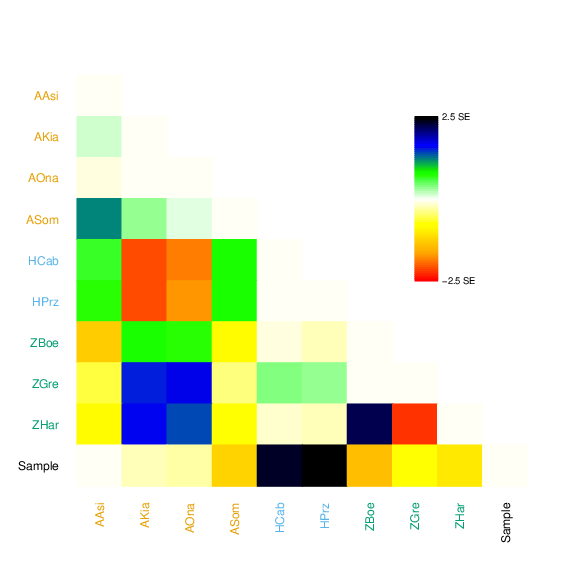

### excl_ts_0_residuals.png

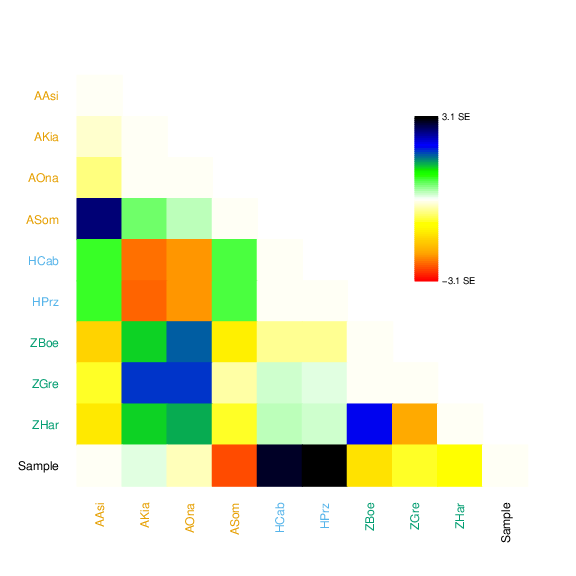

### excl_ts_0_tree.pdf

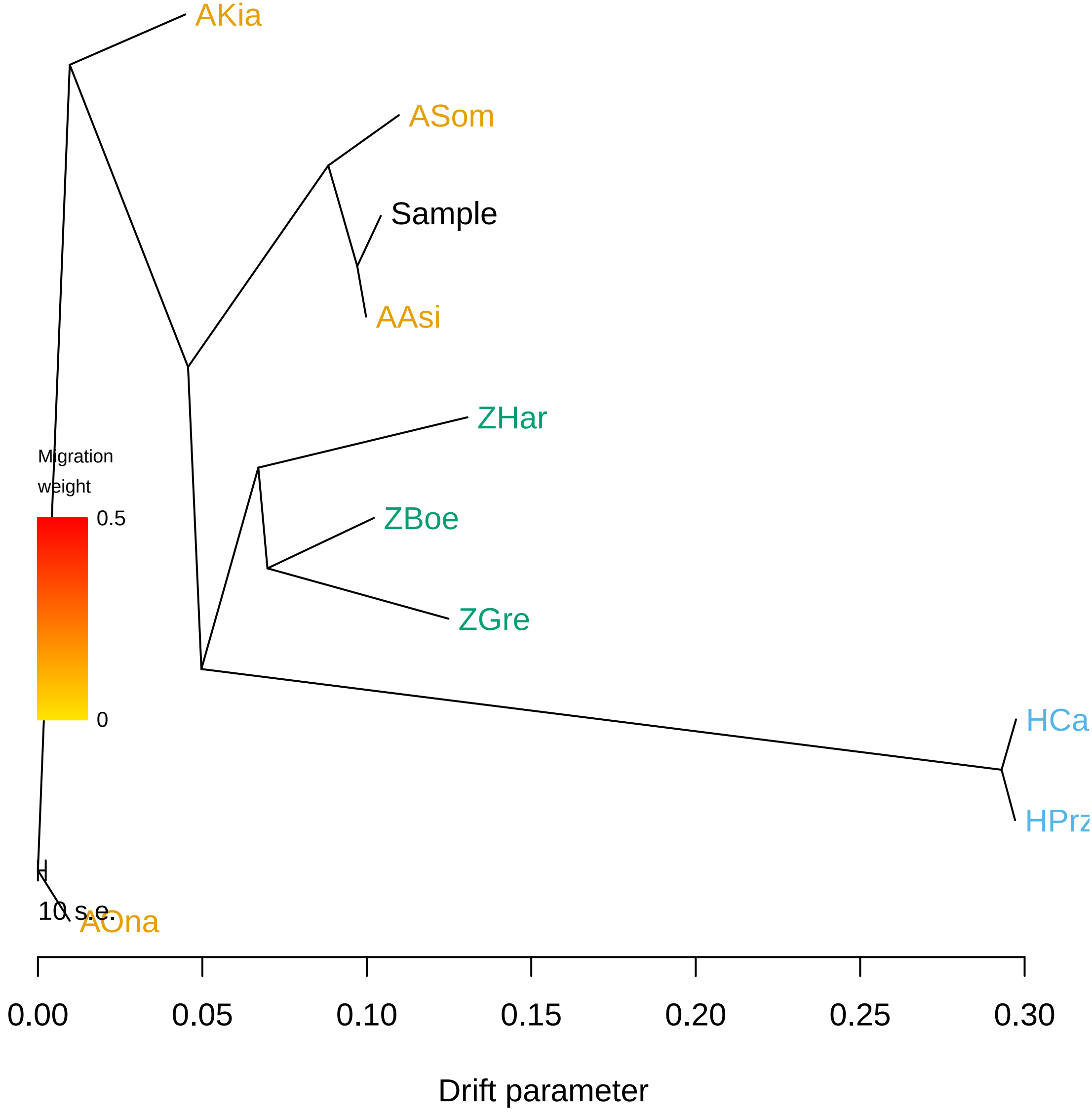

### excl_ts_0_tree.pdf

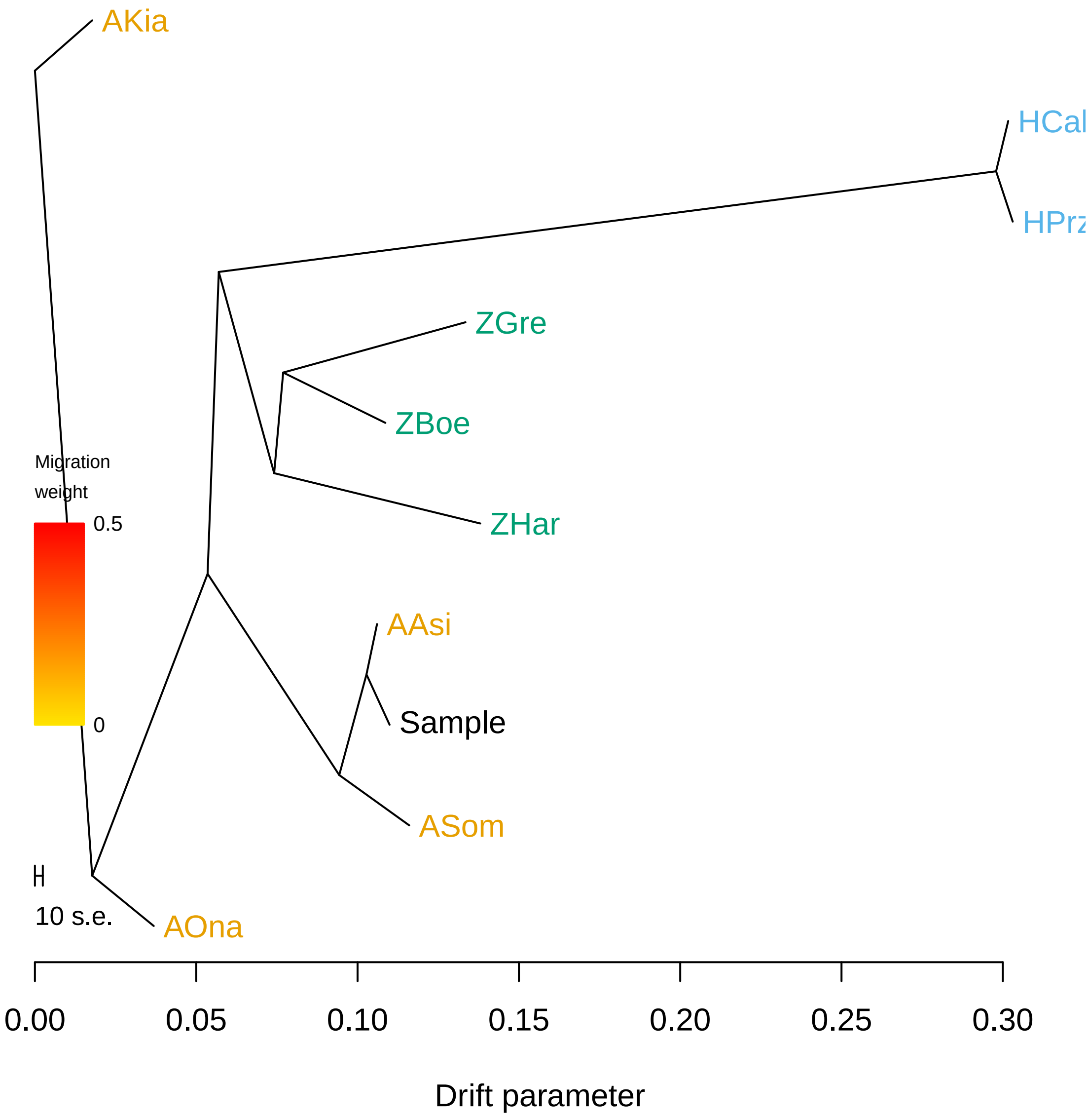

### excl_ts_0_tree.pdf

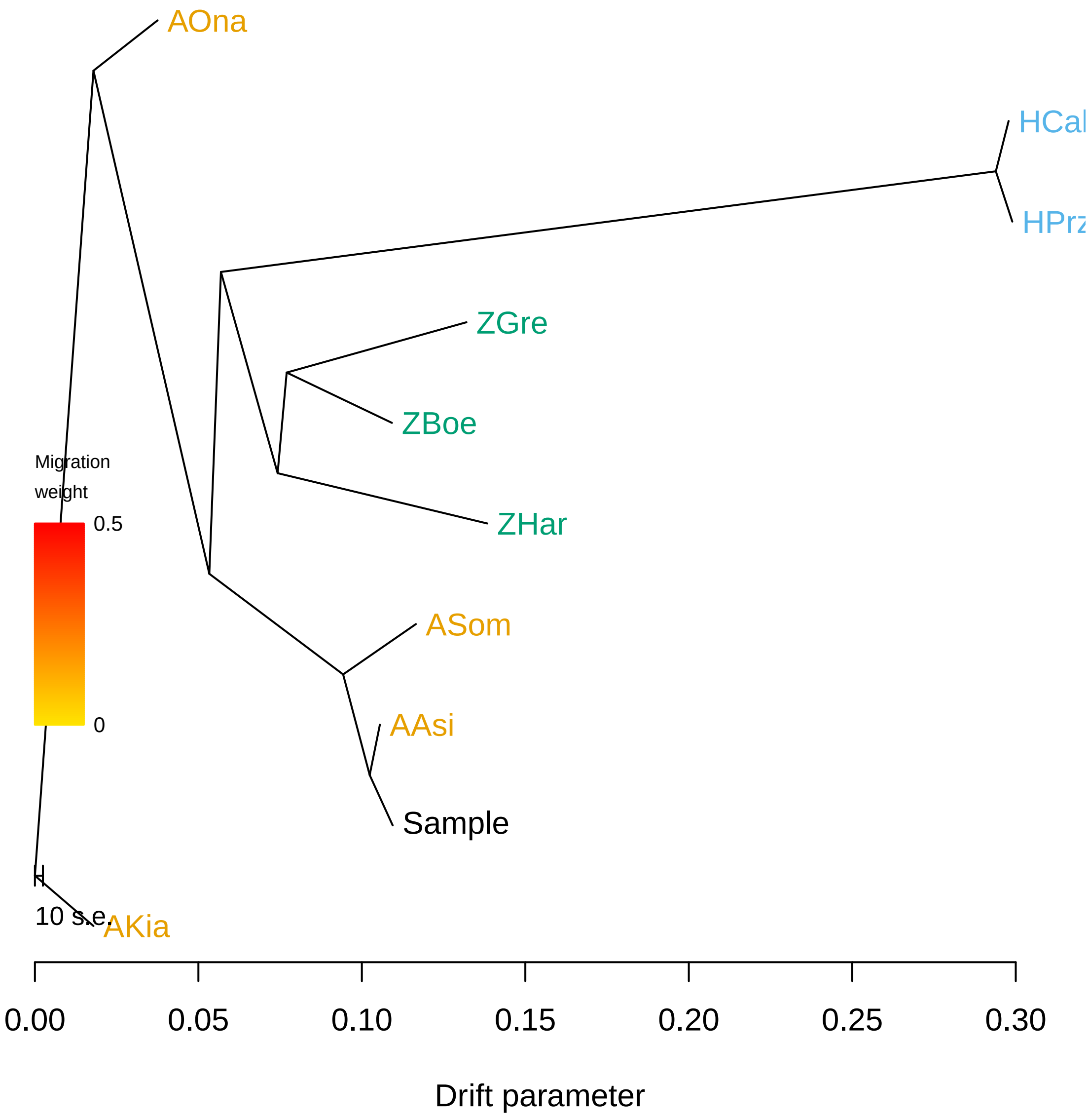

### excl_ts_0_tree.png

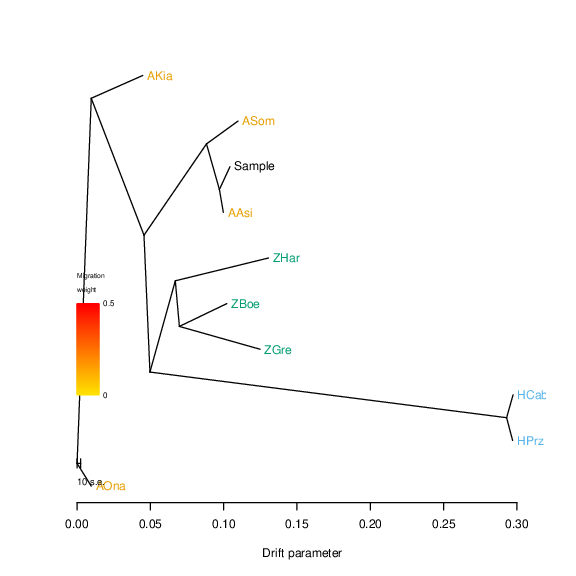

### excl_ts_0_tree.png

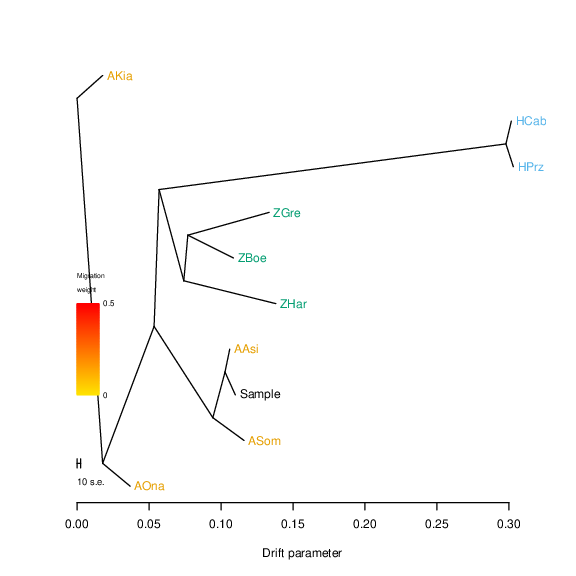

### excl_ts_0_tree.png

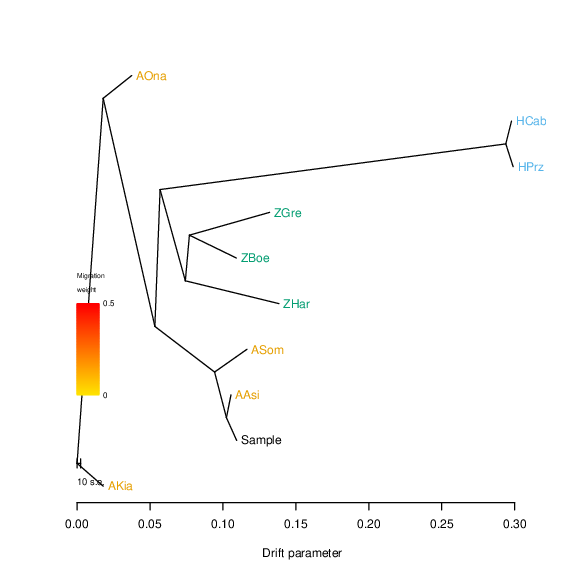

### excl_ts_1_residuals.pdf

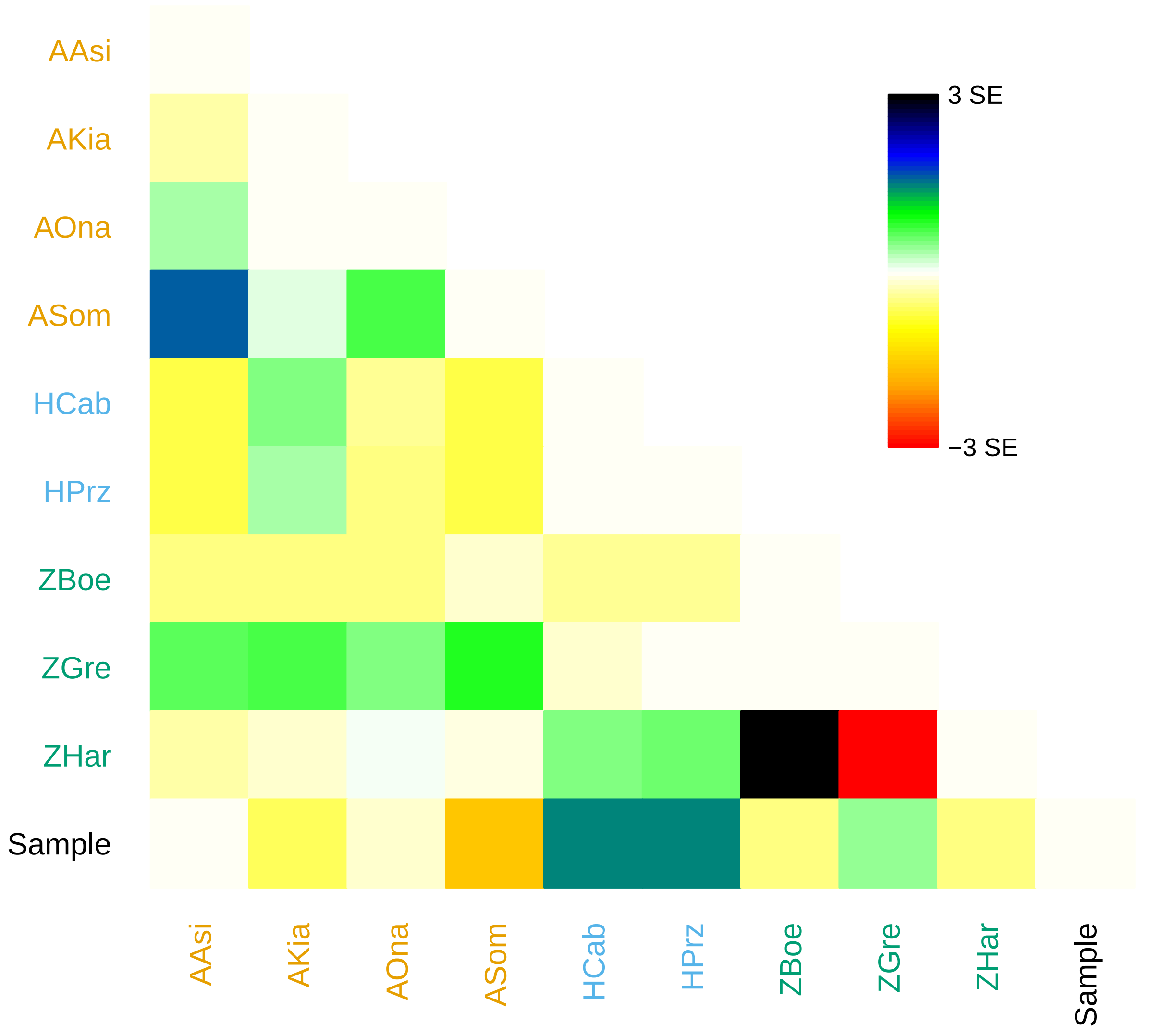

### excl_ts_1_residuals.pdf

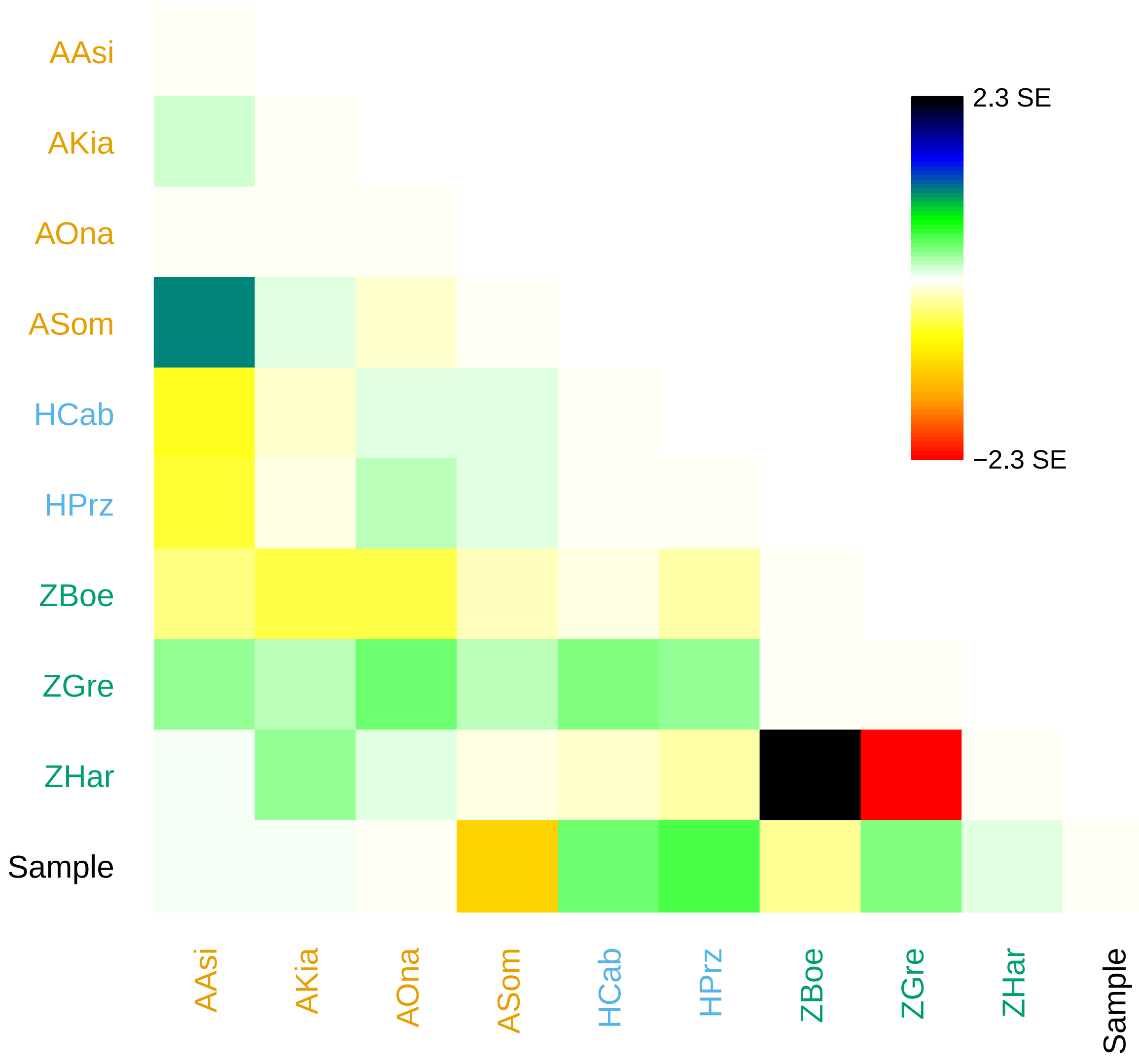

### excl_ts_1_residuals.pdf

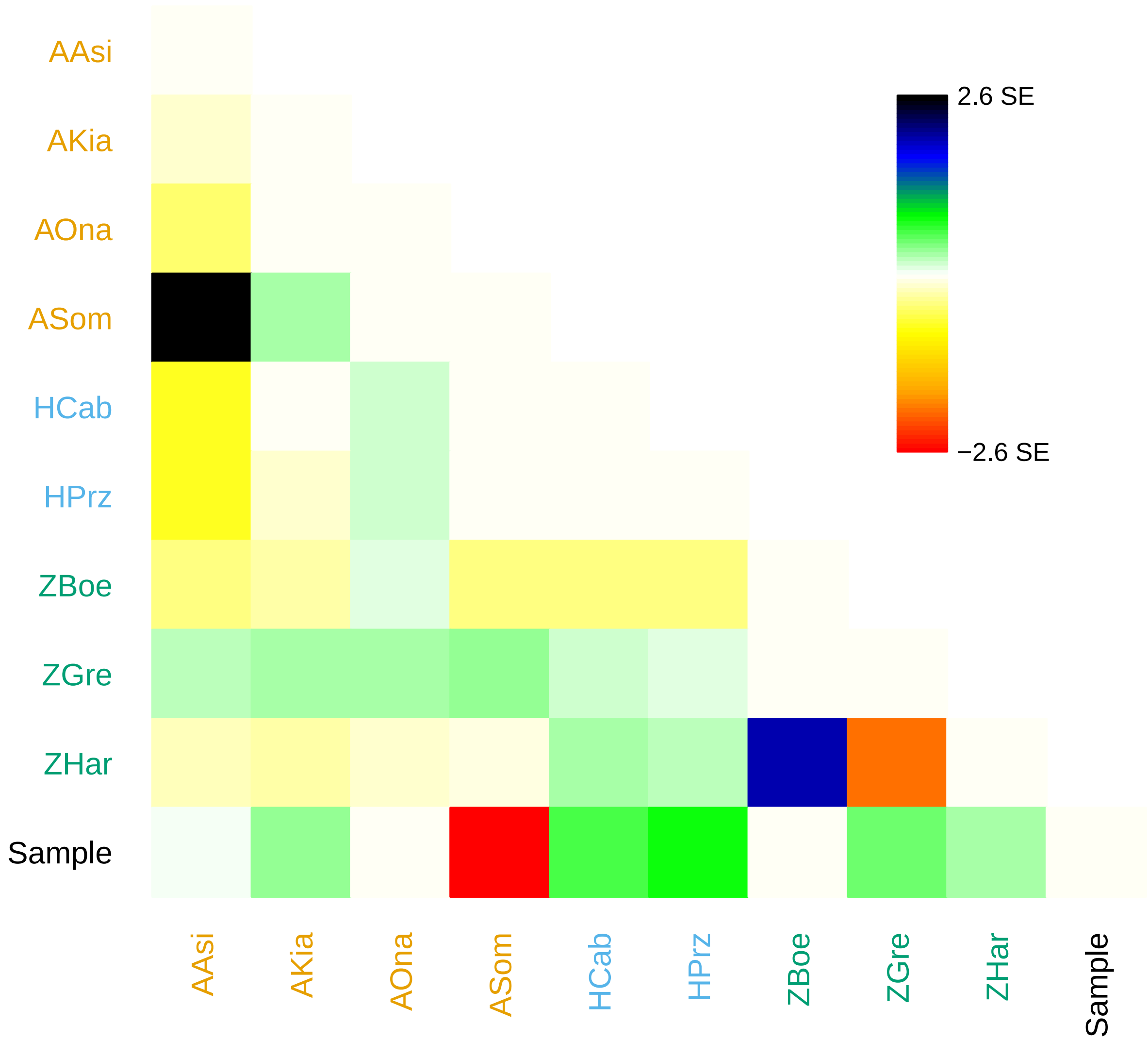

### excl_ts_1_residuals.png

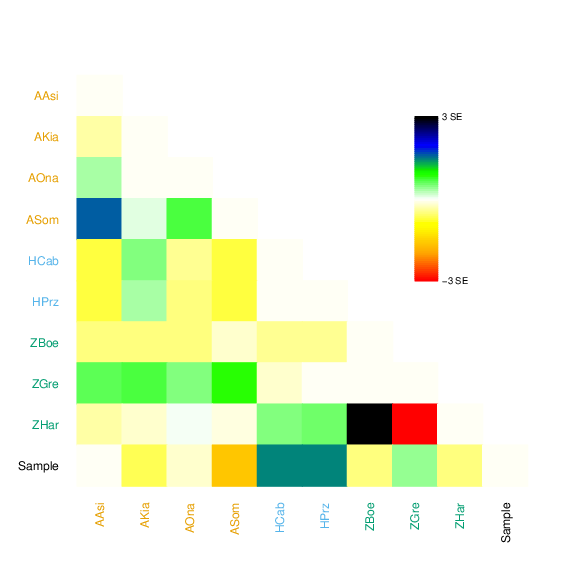

### excl_ts_1_residuals.png

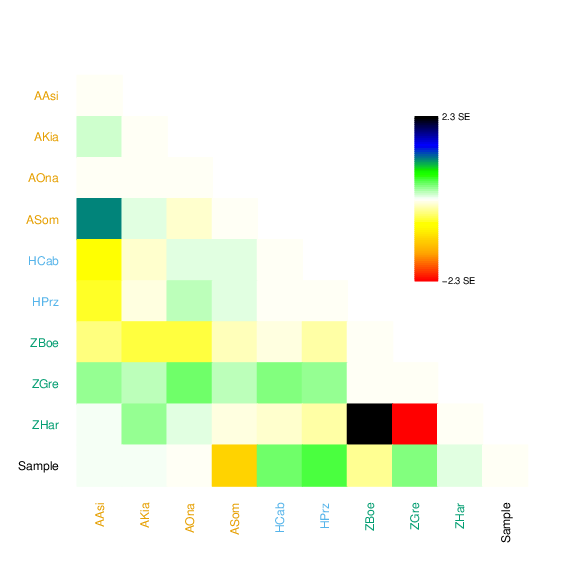

### excl_ts_1_residuals.png

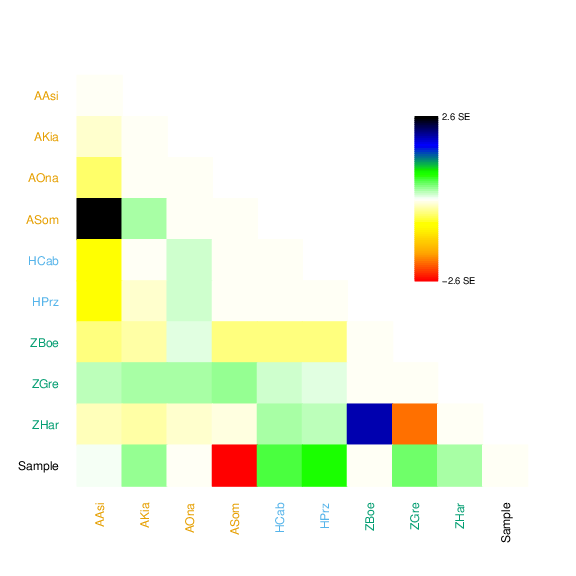

### excl_ts_1_tree.pdf

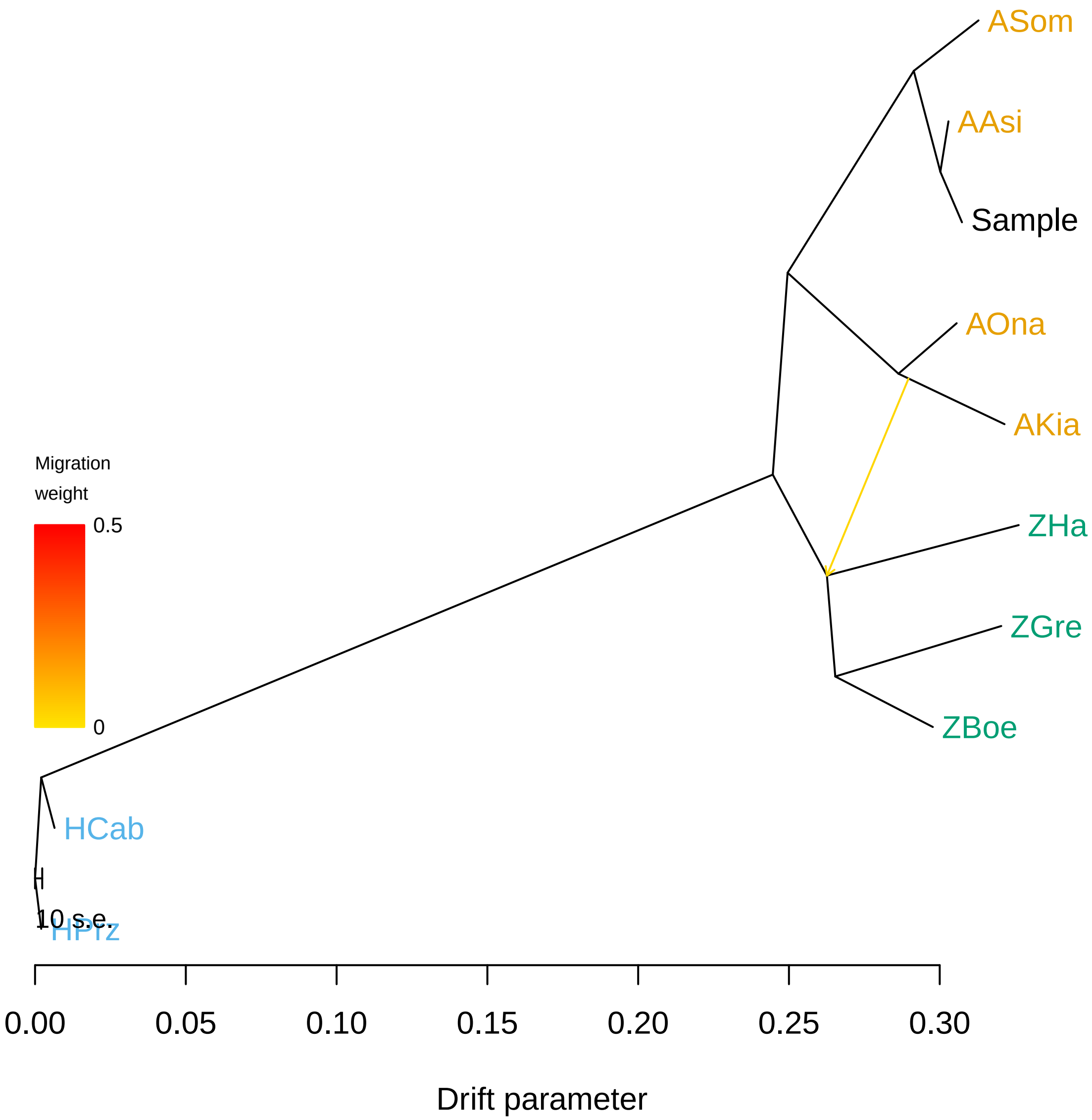

### excl_ts_1_tree.pdf

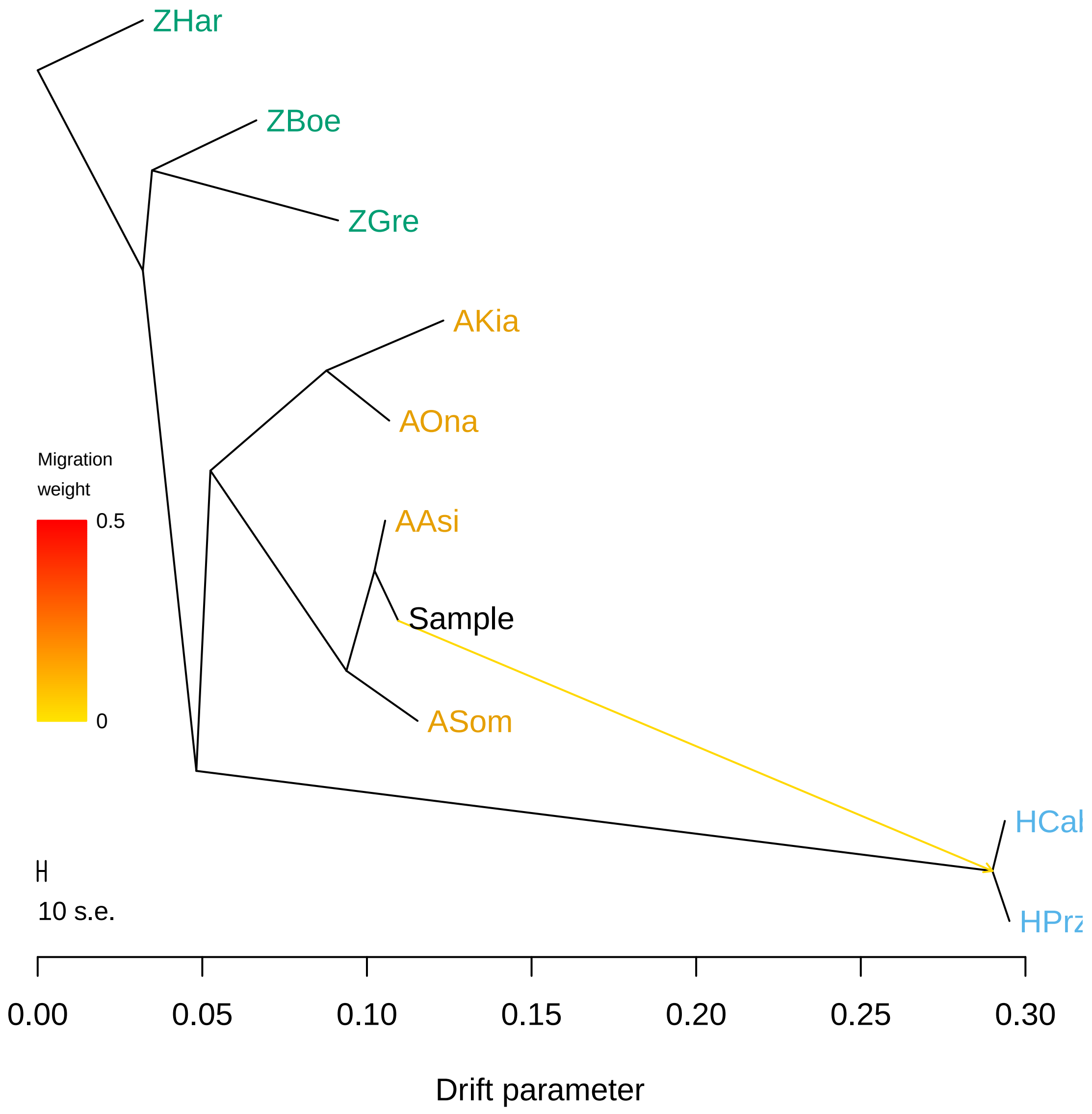

### excl_ts_1_tree.pdf

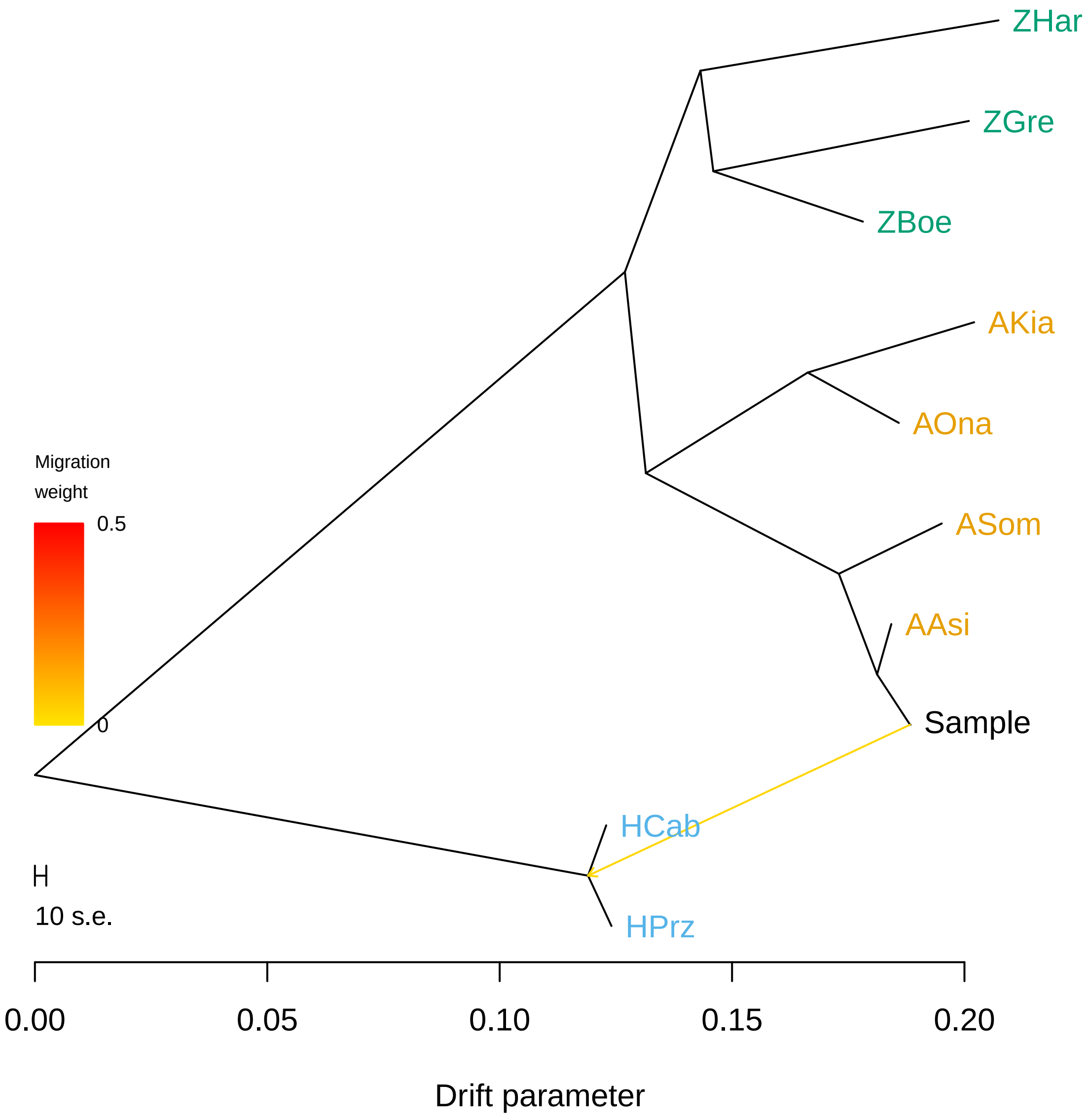

### excl_ts_1_tree.png

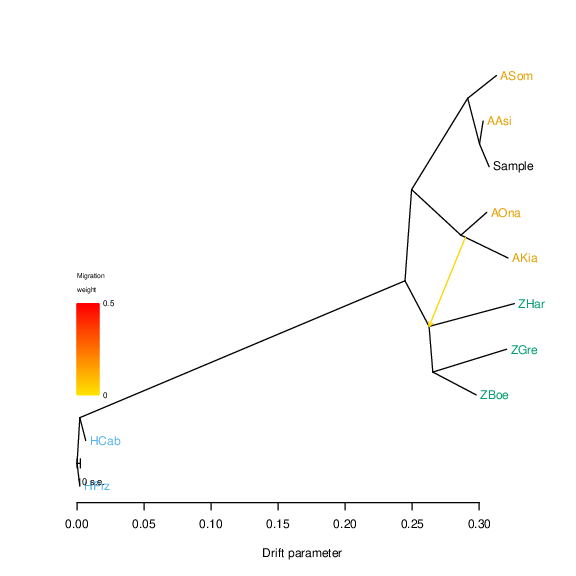

### excl_ts_1_tree.png

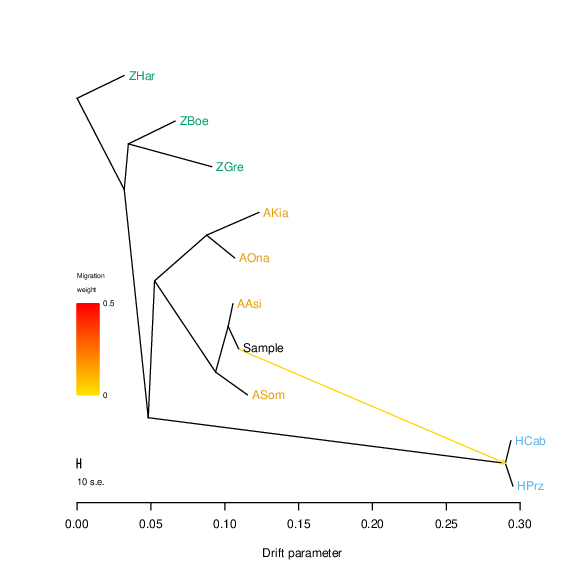

### excl_ts_1_tree.png

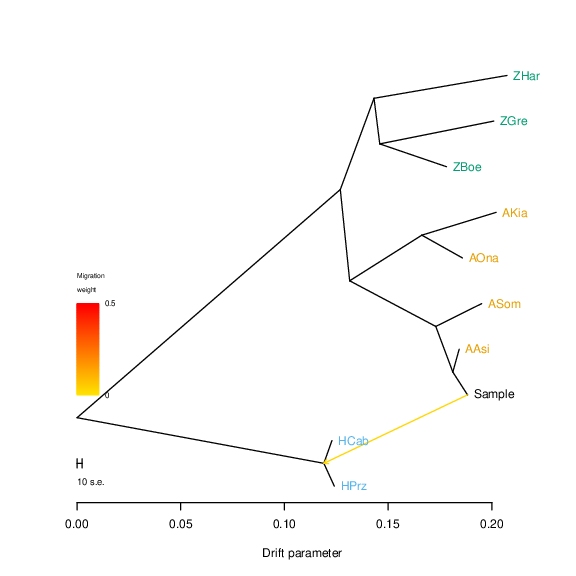

### excl_ts_k2.pdf

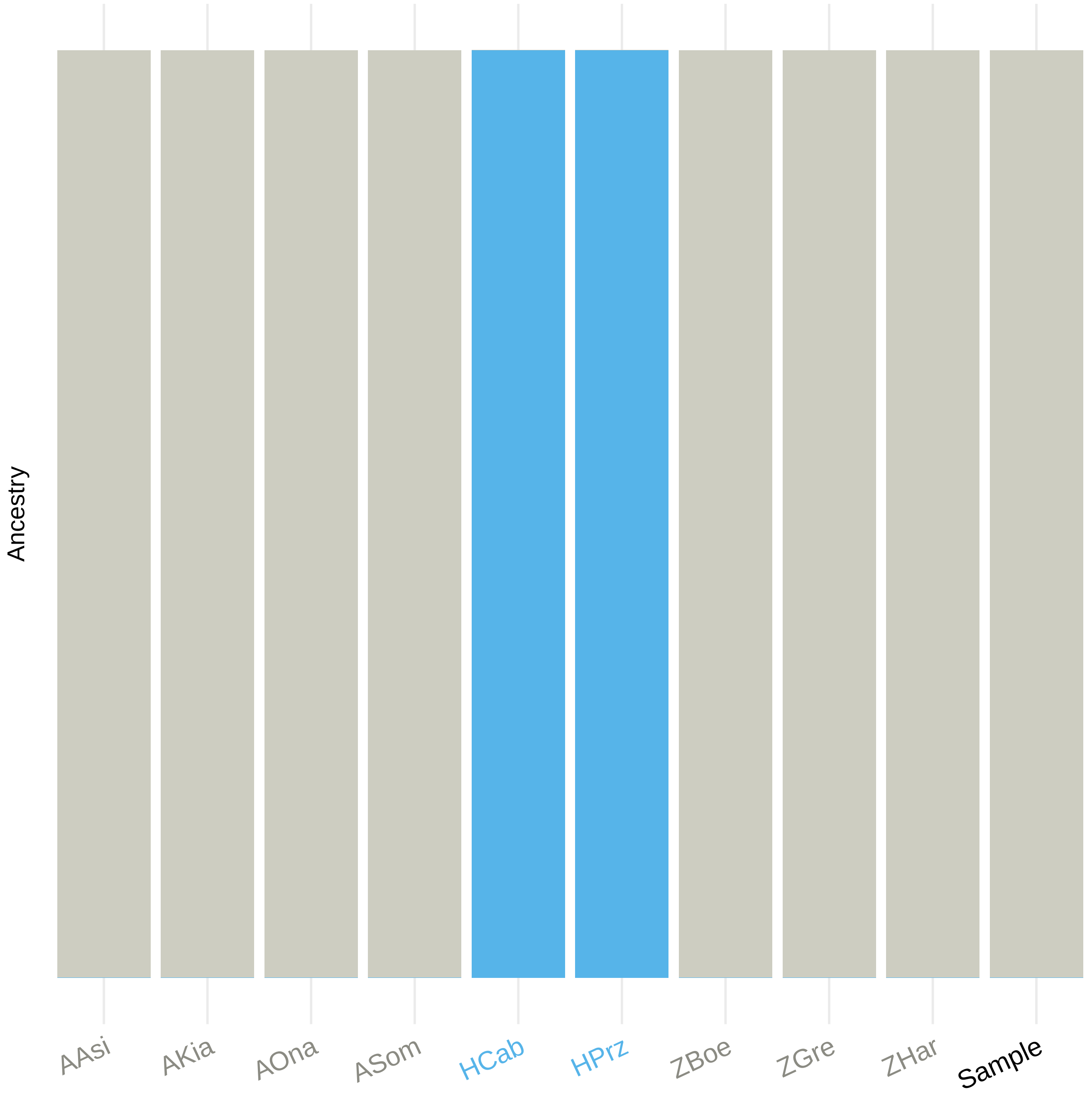

### excl_ts_k2.pdf

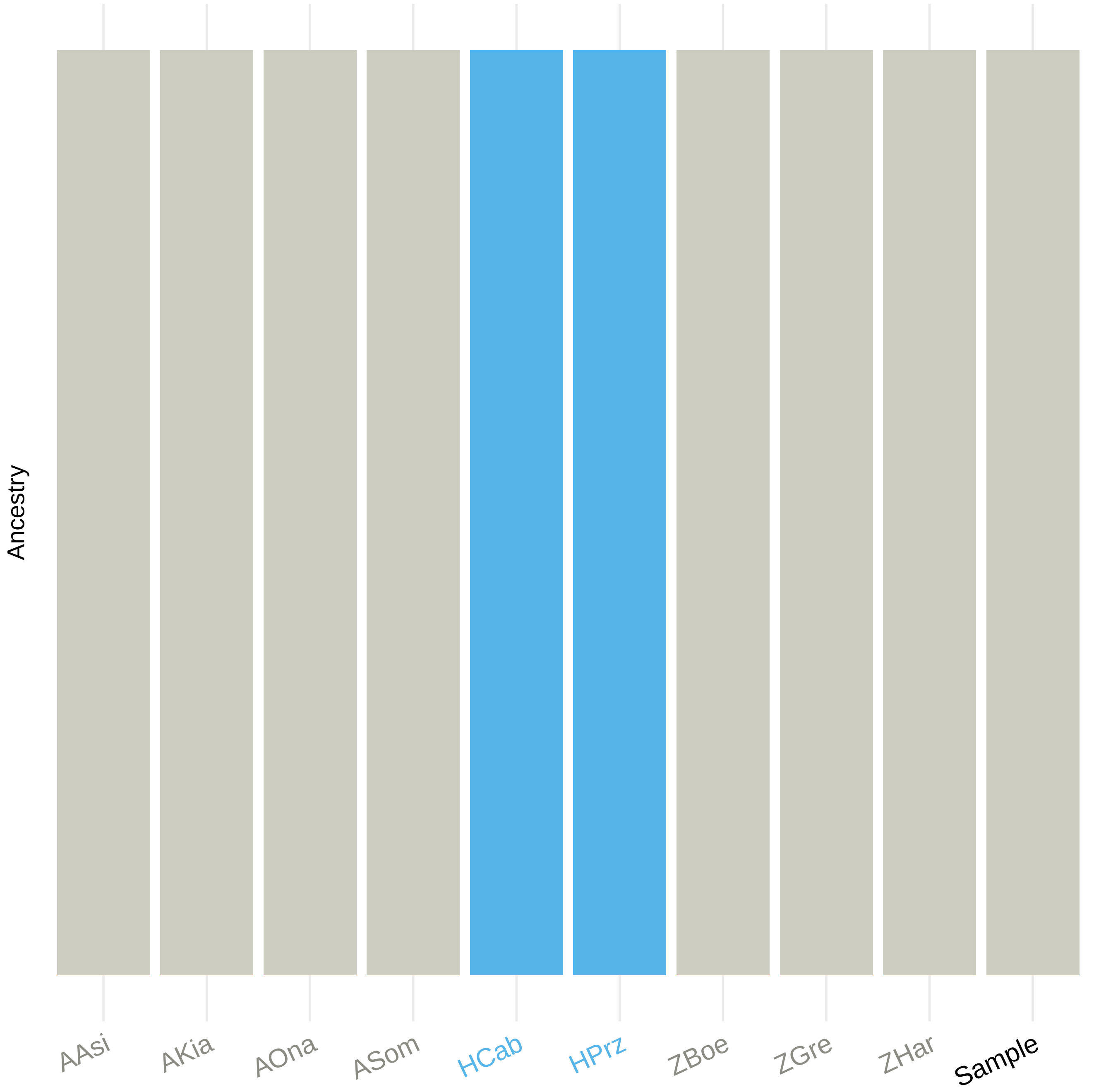

### excl_ts_k2.pdf

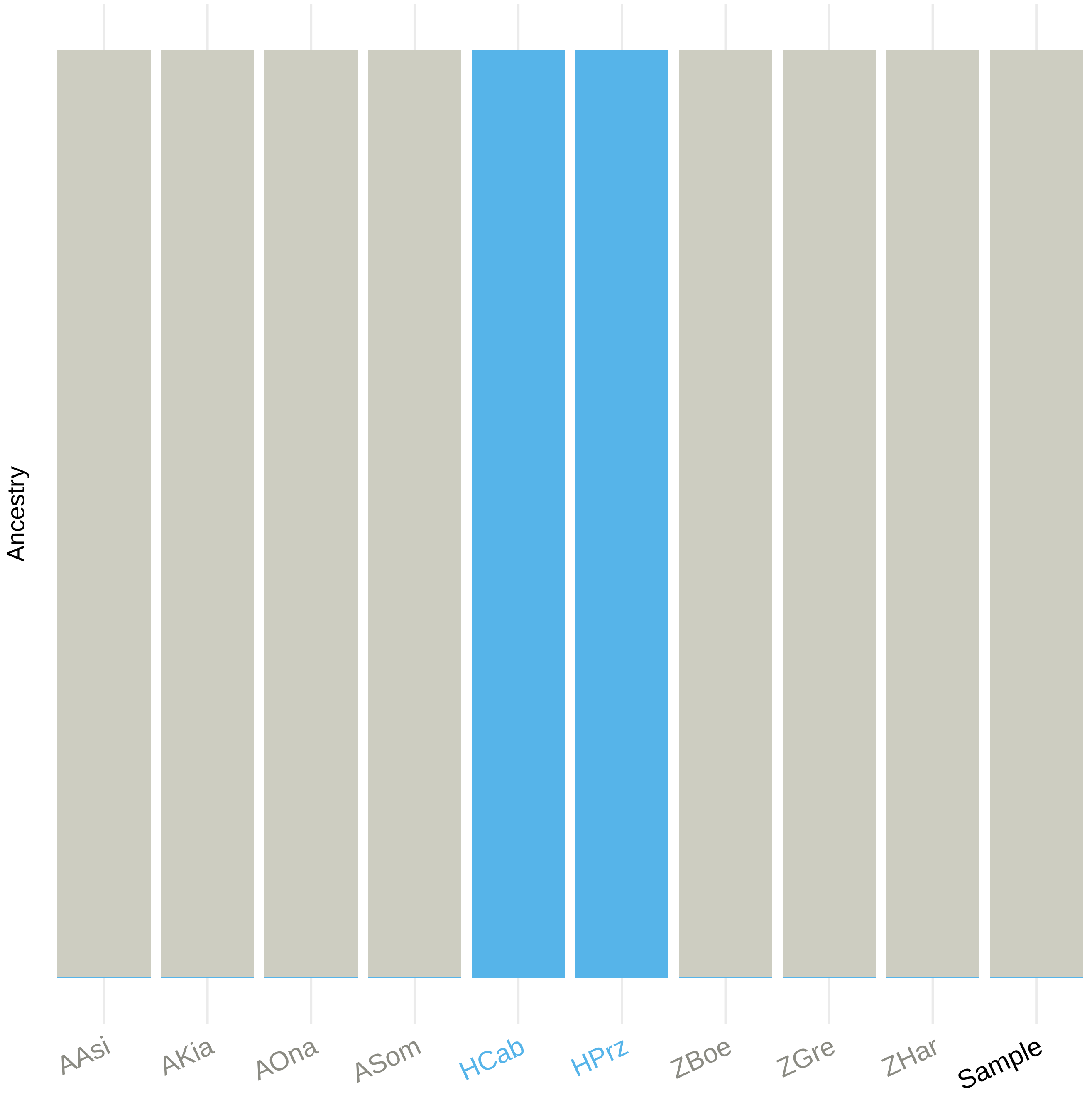

### excl_ts_k3.pdf

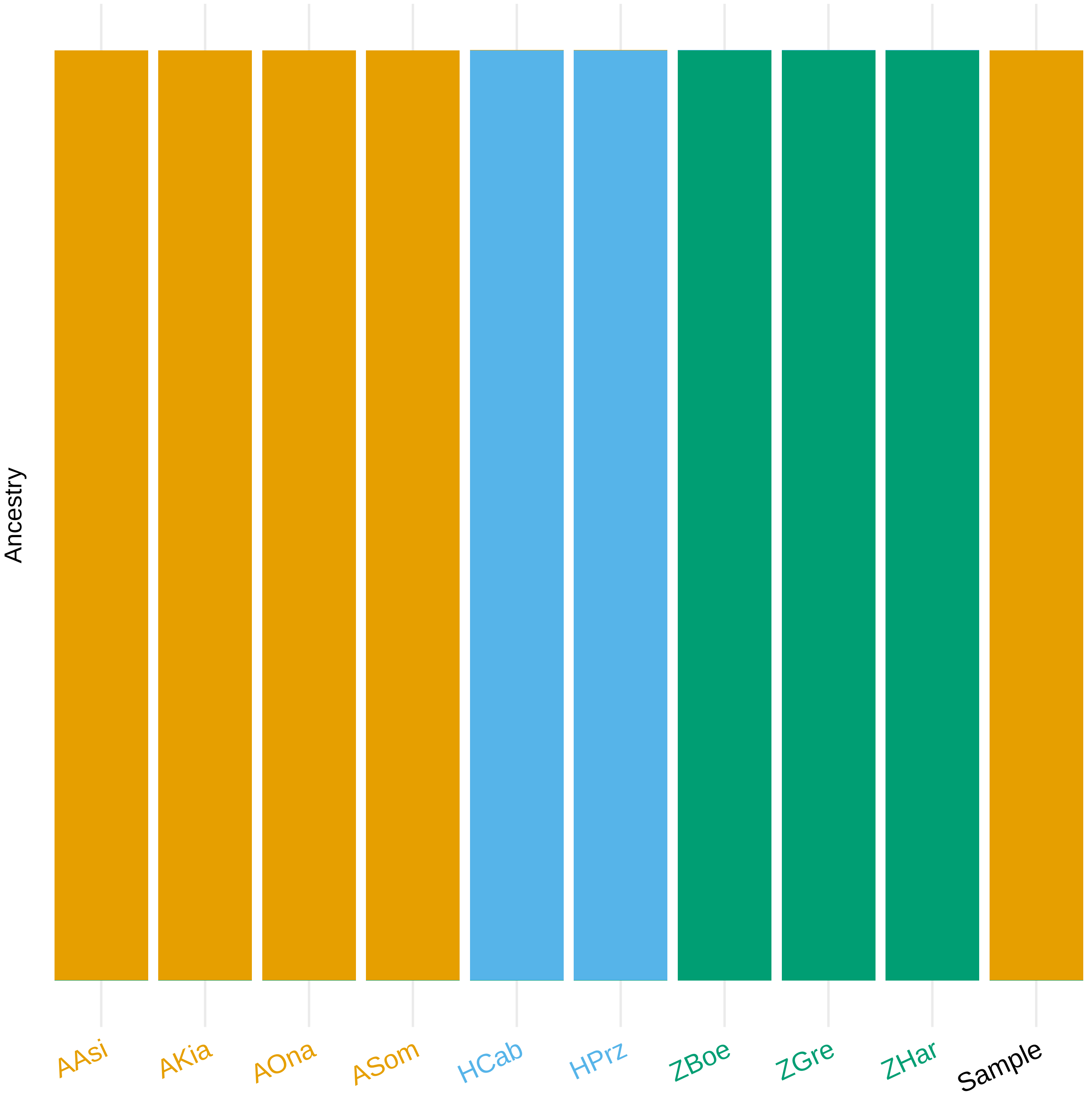

### excl_ts_k3.pdf

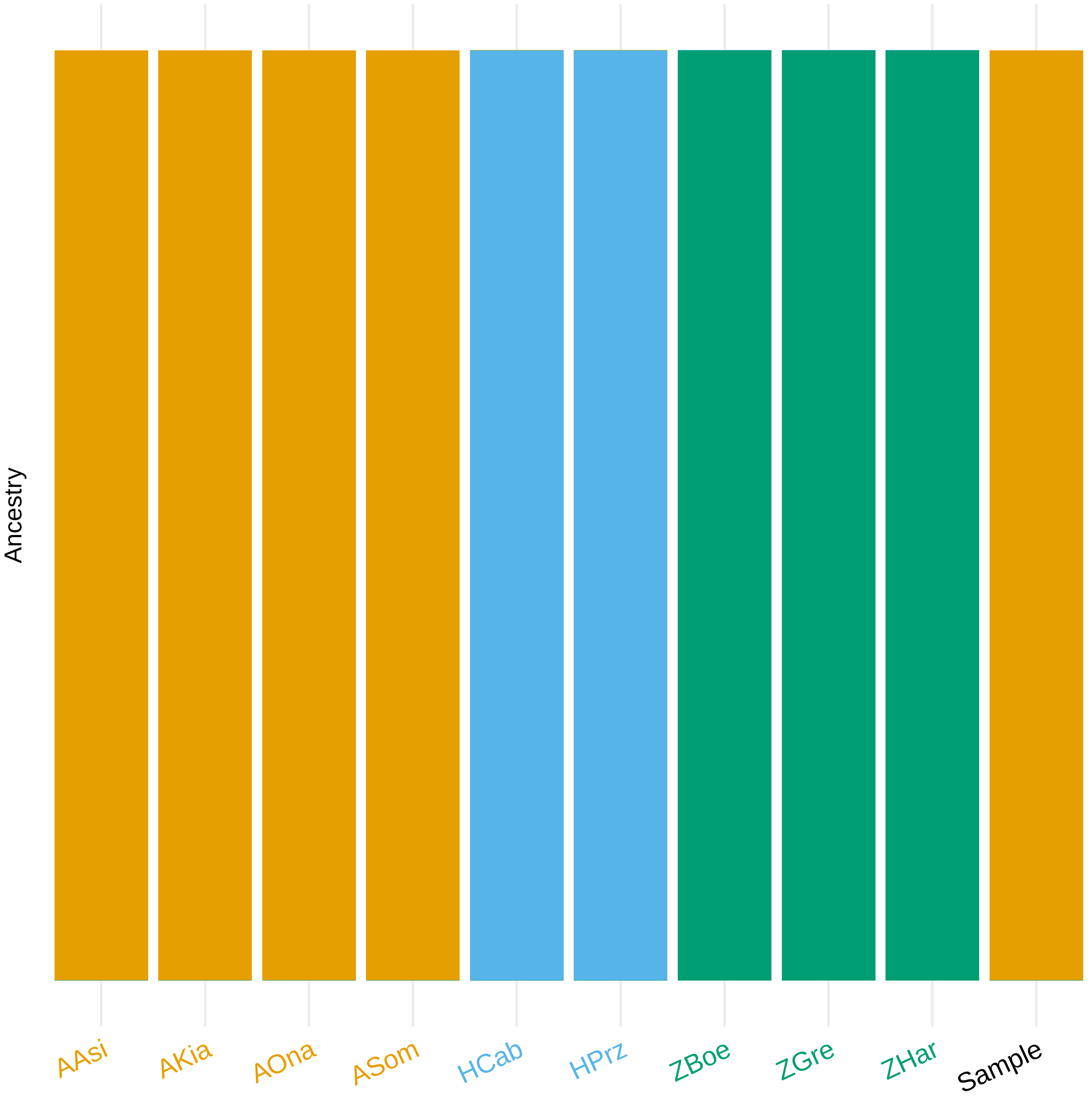

### incl_ts.pdf

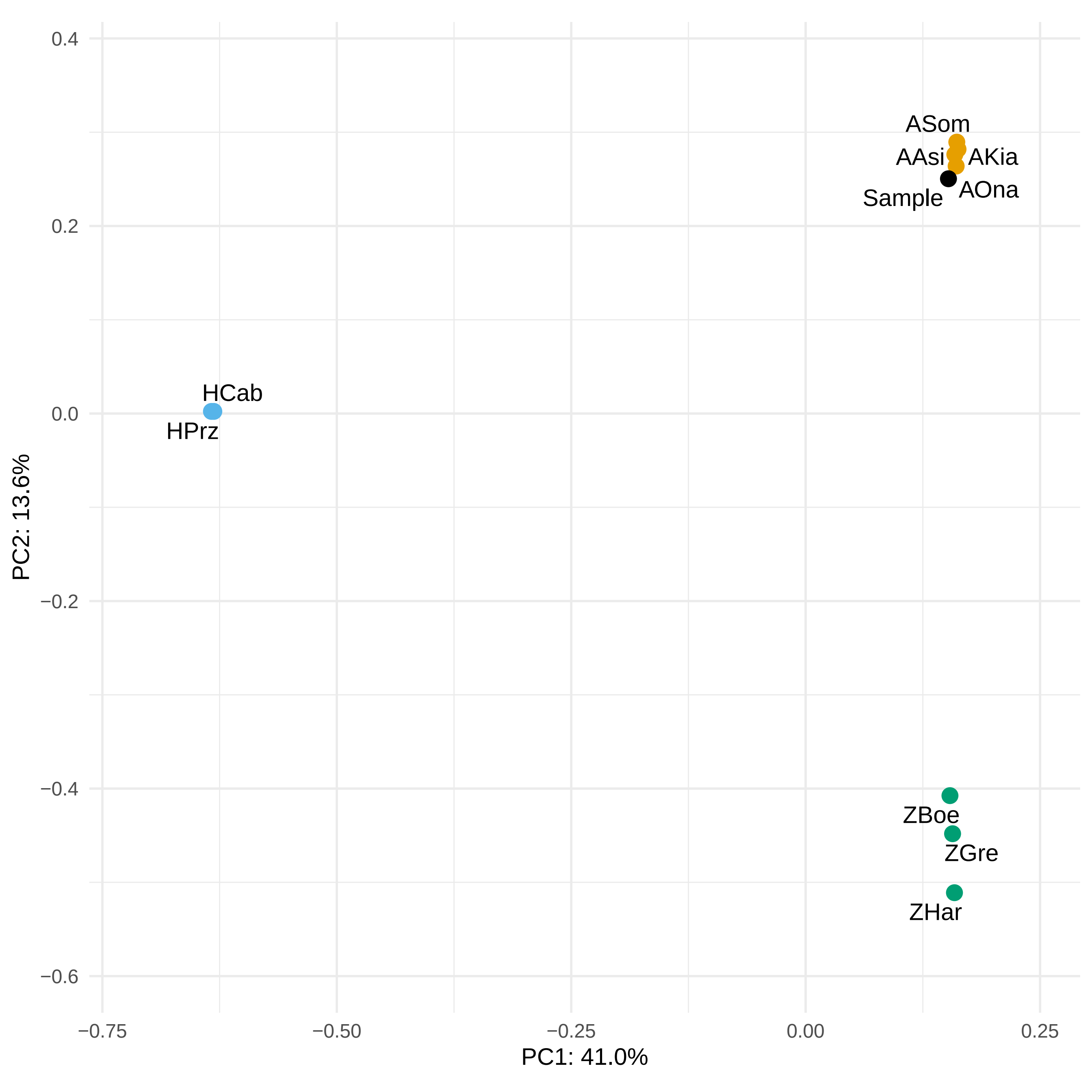

### incl_ts.pdf

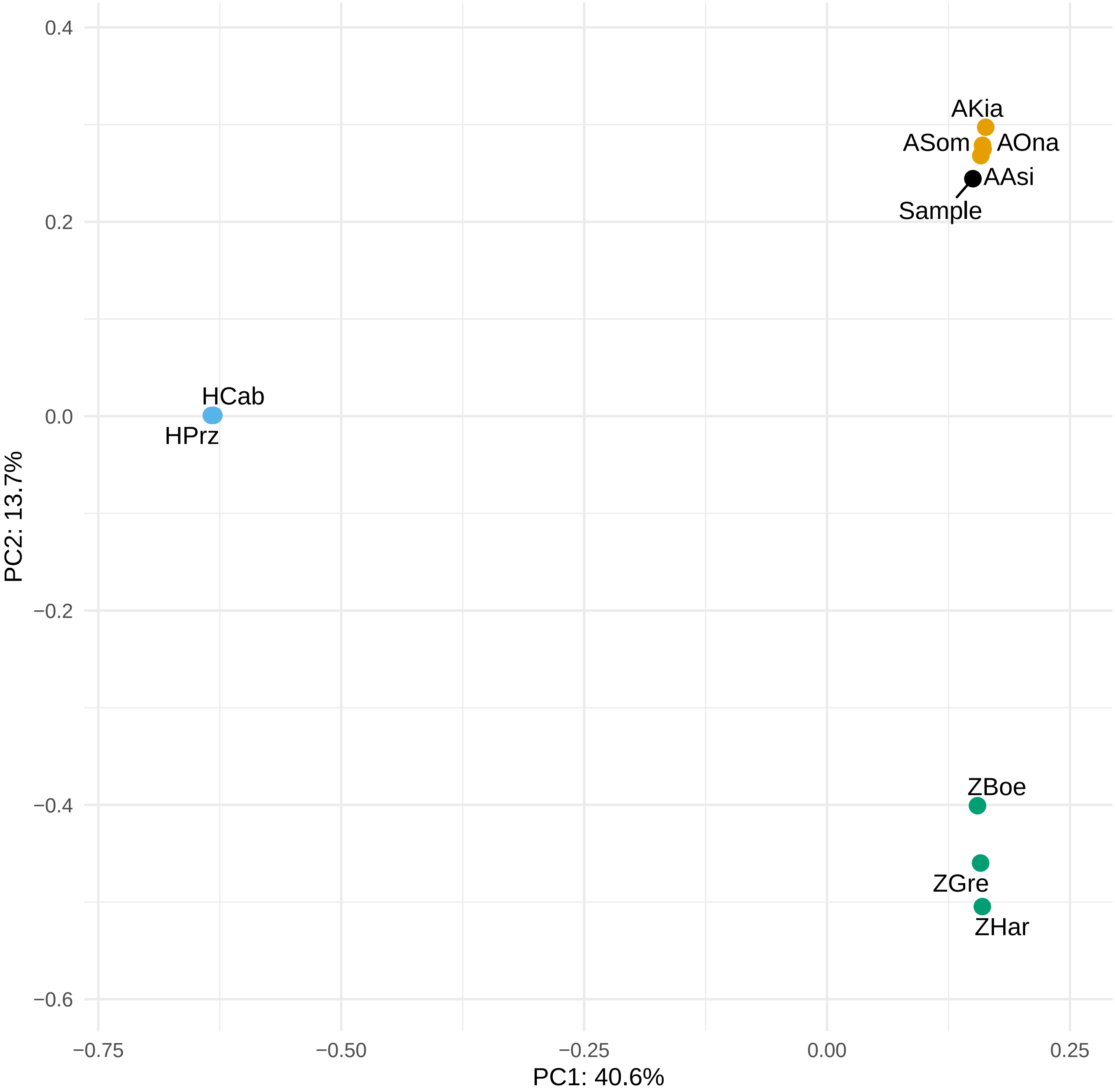

### incl_ts.pdf

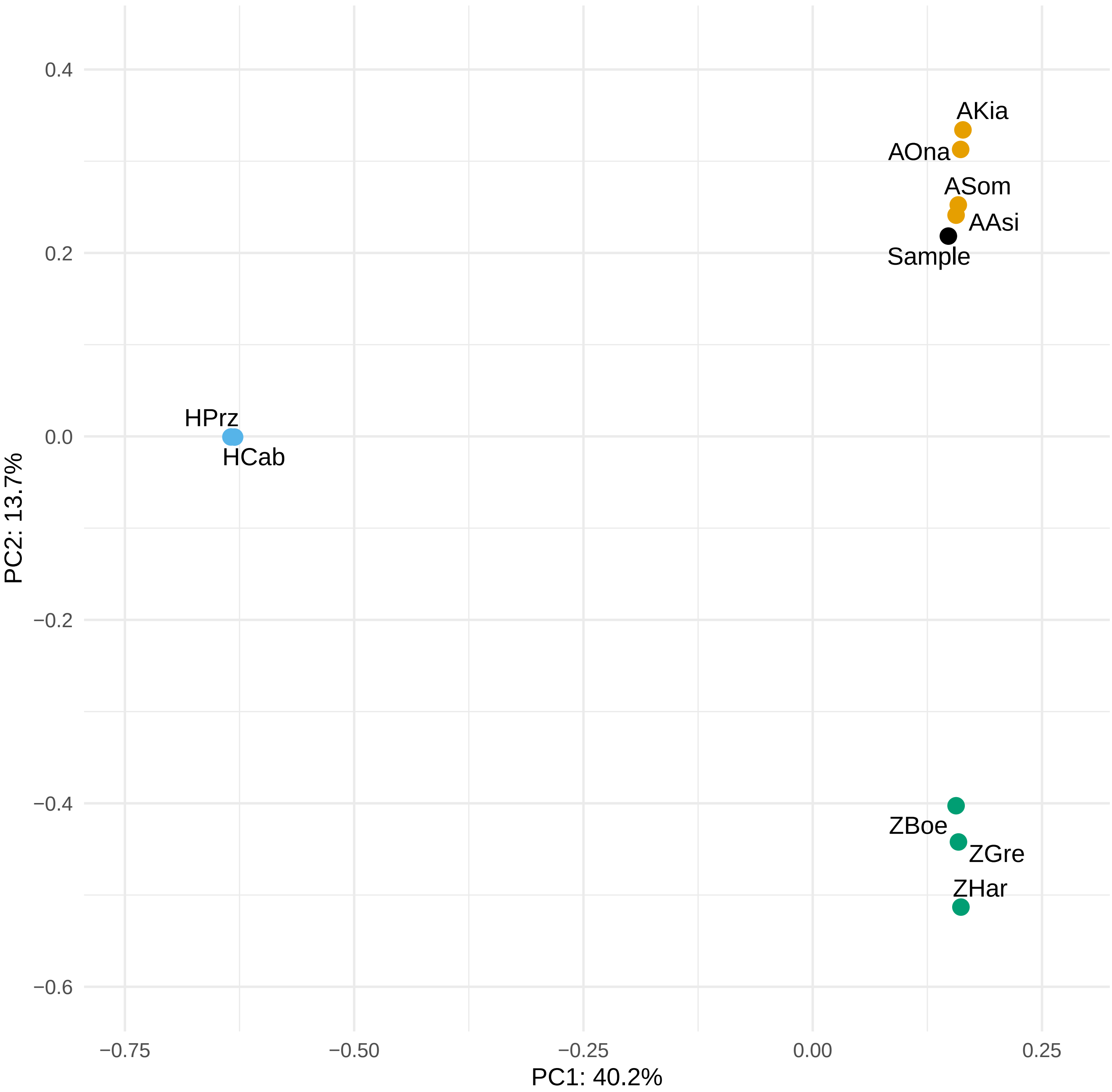

### incl_ts.png

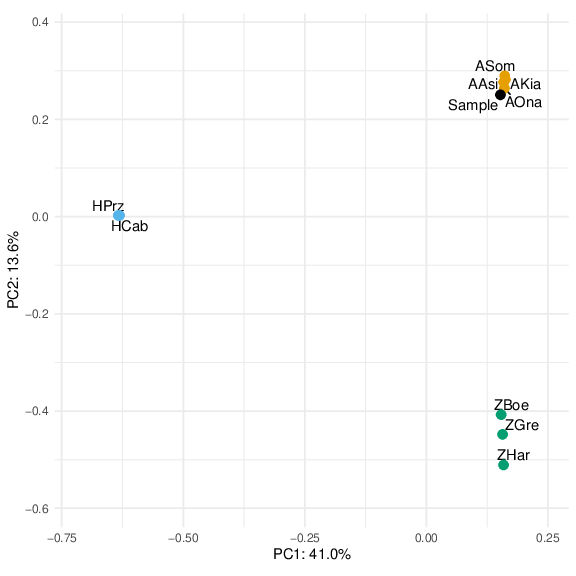

### incl_ts.png

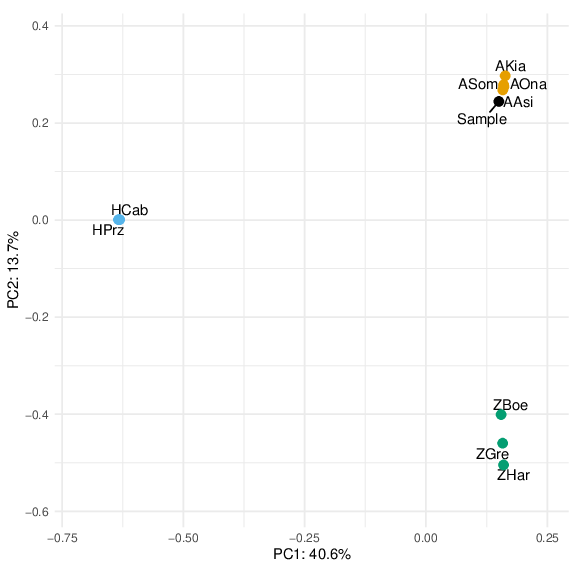

### incl_ts.png

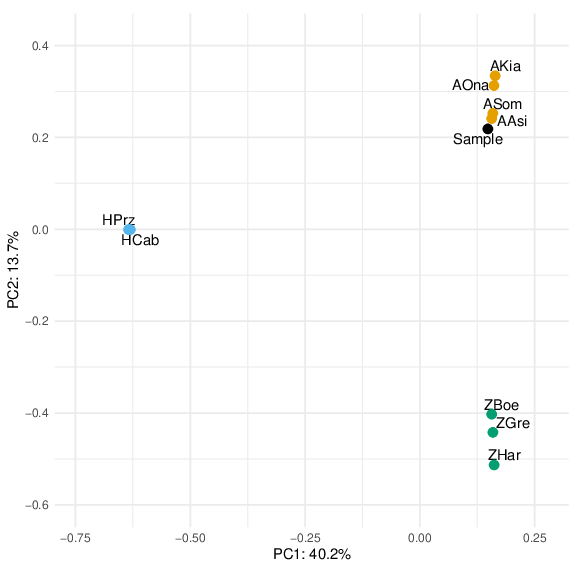

### incl_ts_0_residuals.pdf

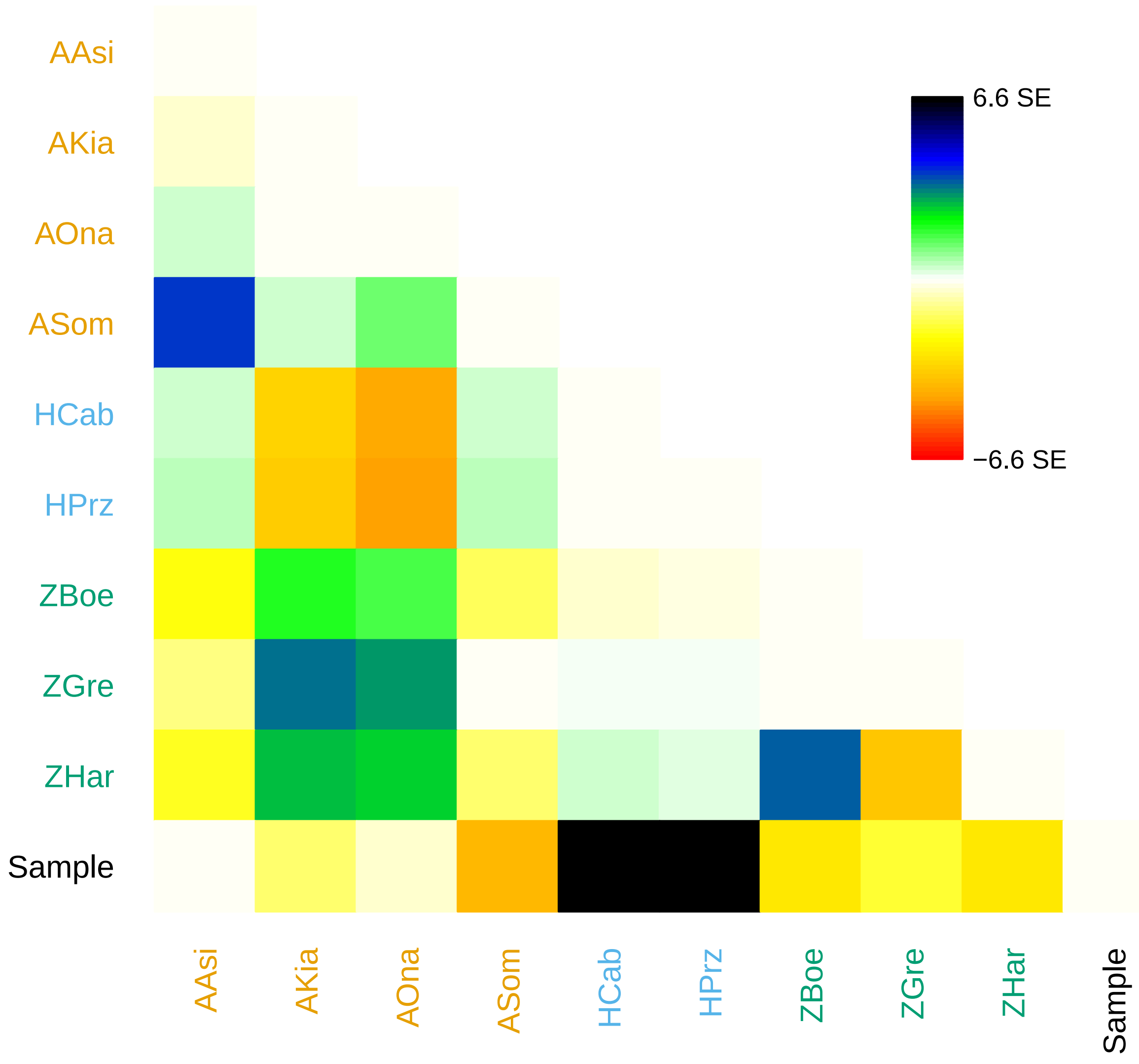

### incl_ts_0_residuals.pdf

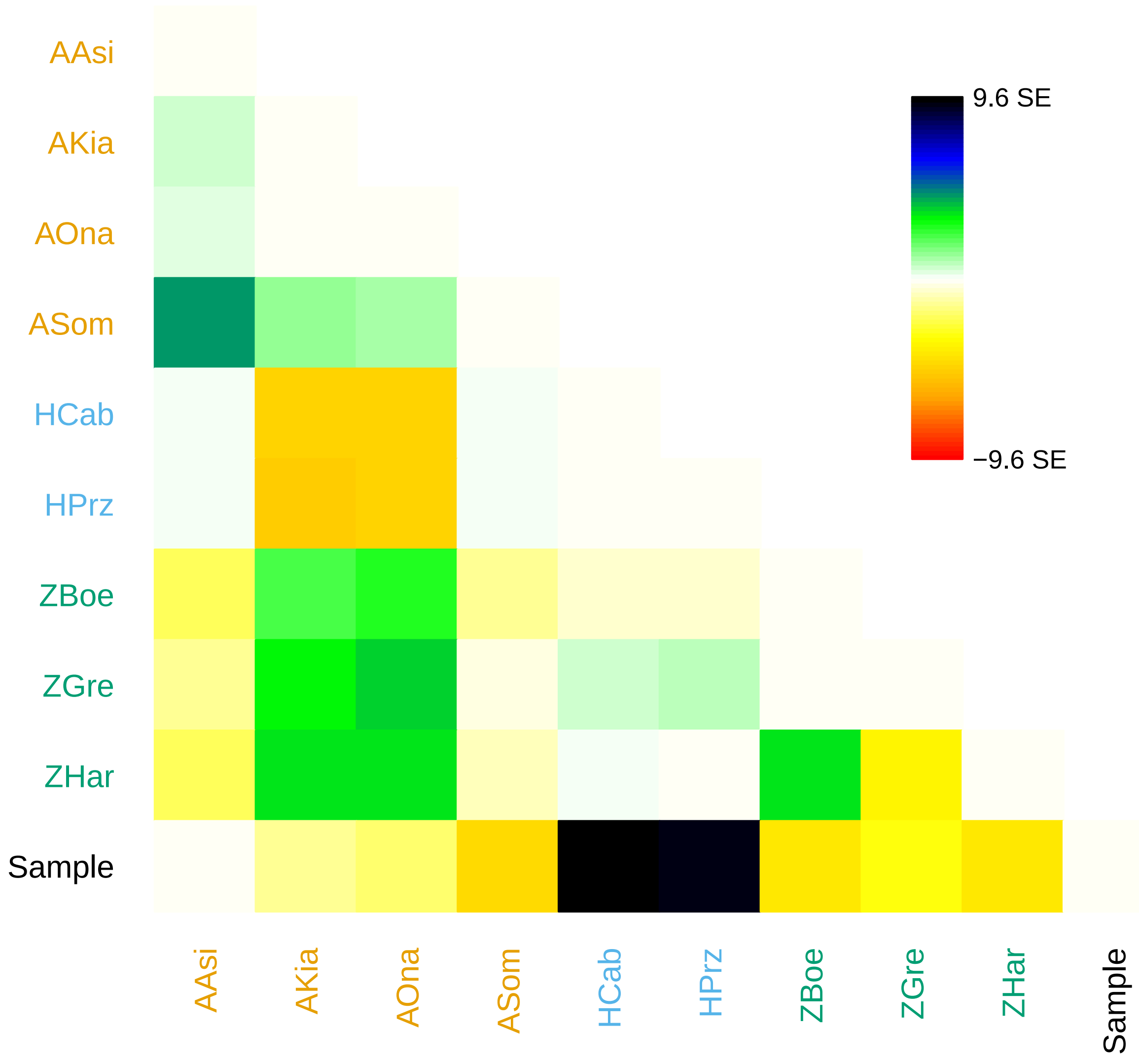

### incl_ts_0_residuals.pdf

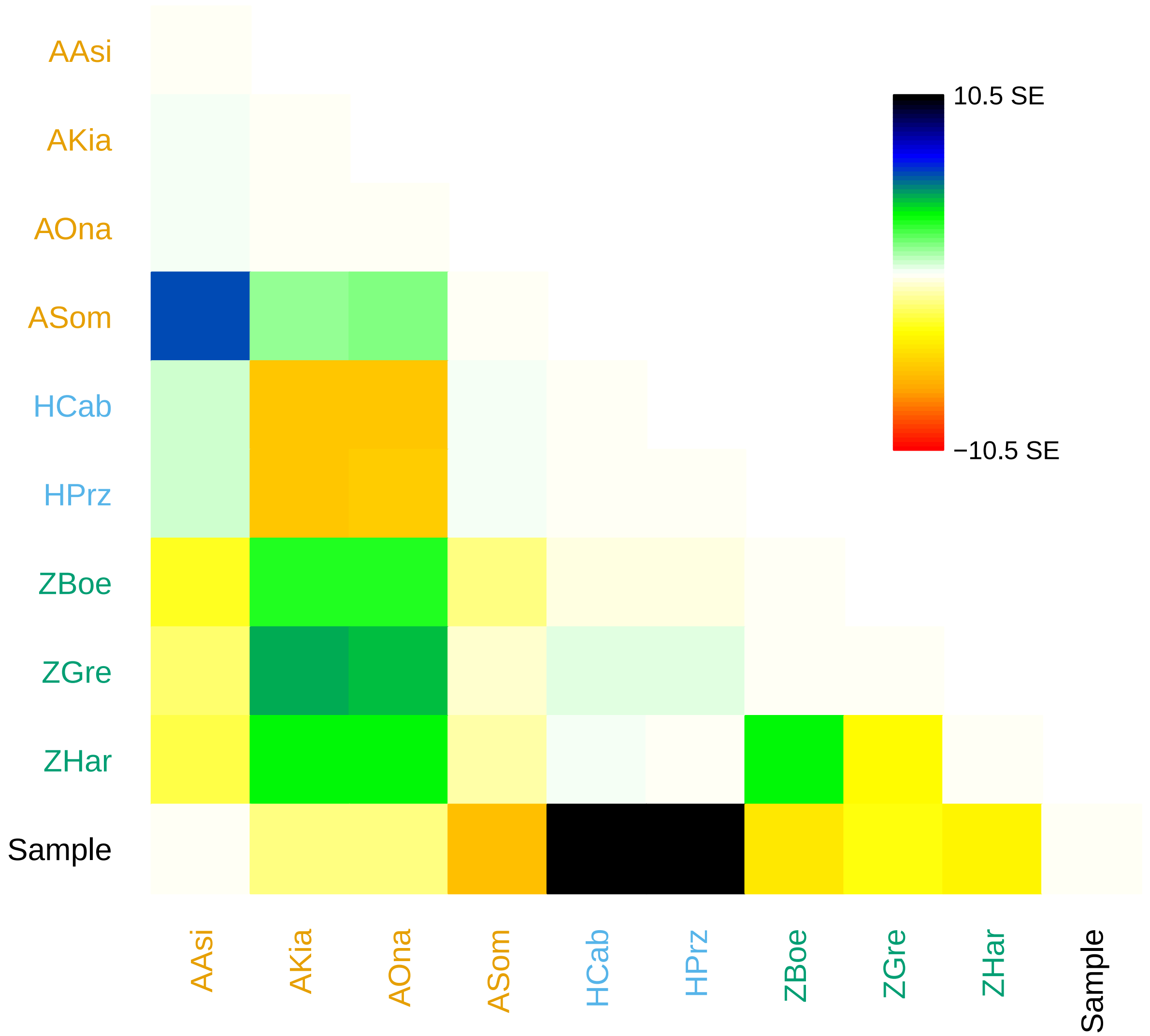

### incl_ts_0_residuals.png

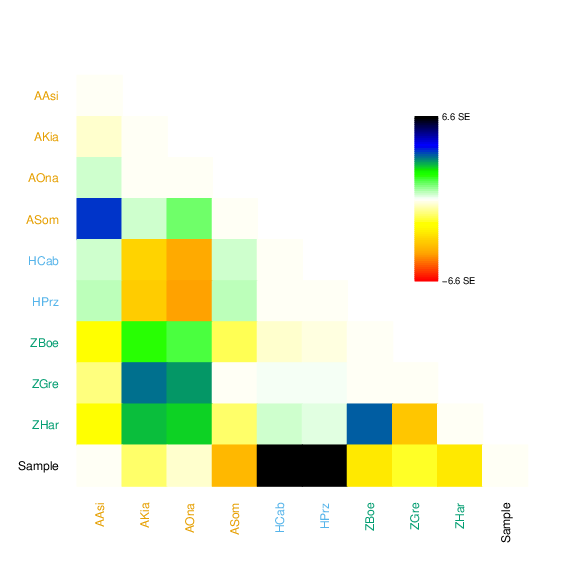

### incl_ts_0_residuals.png

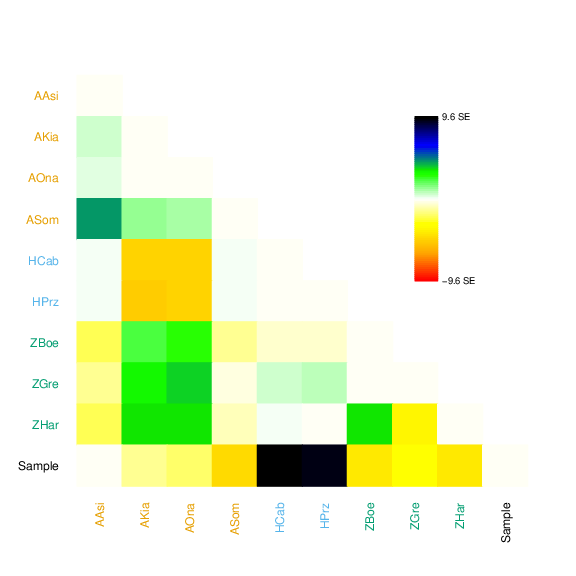

### incl_ts_0_residuals.png

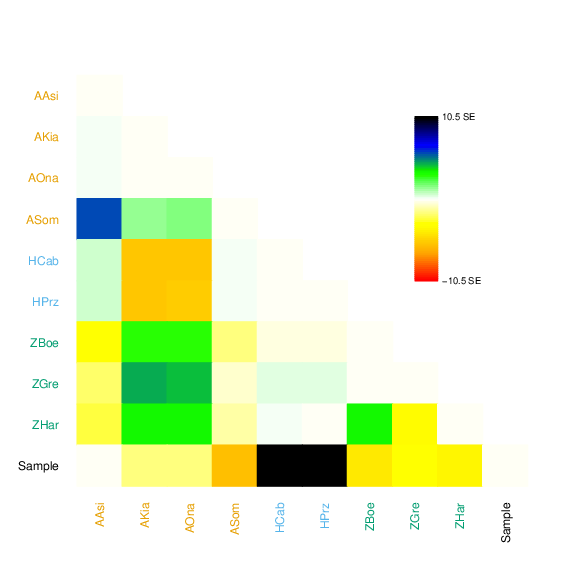

### incl_ts_0_tree.pdf

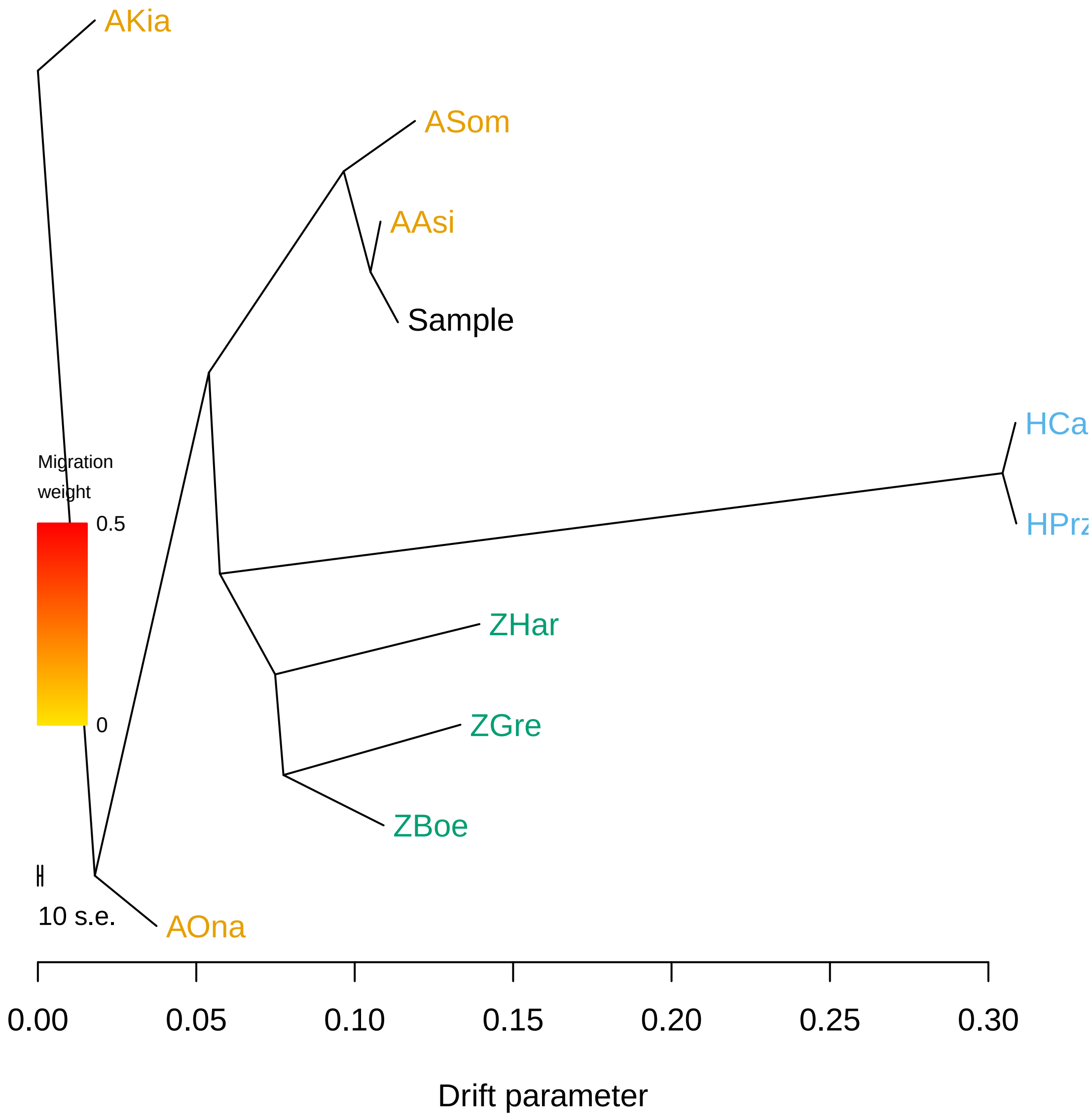

### incl_ts_0_tree.pdf

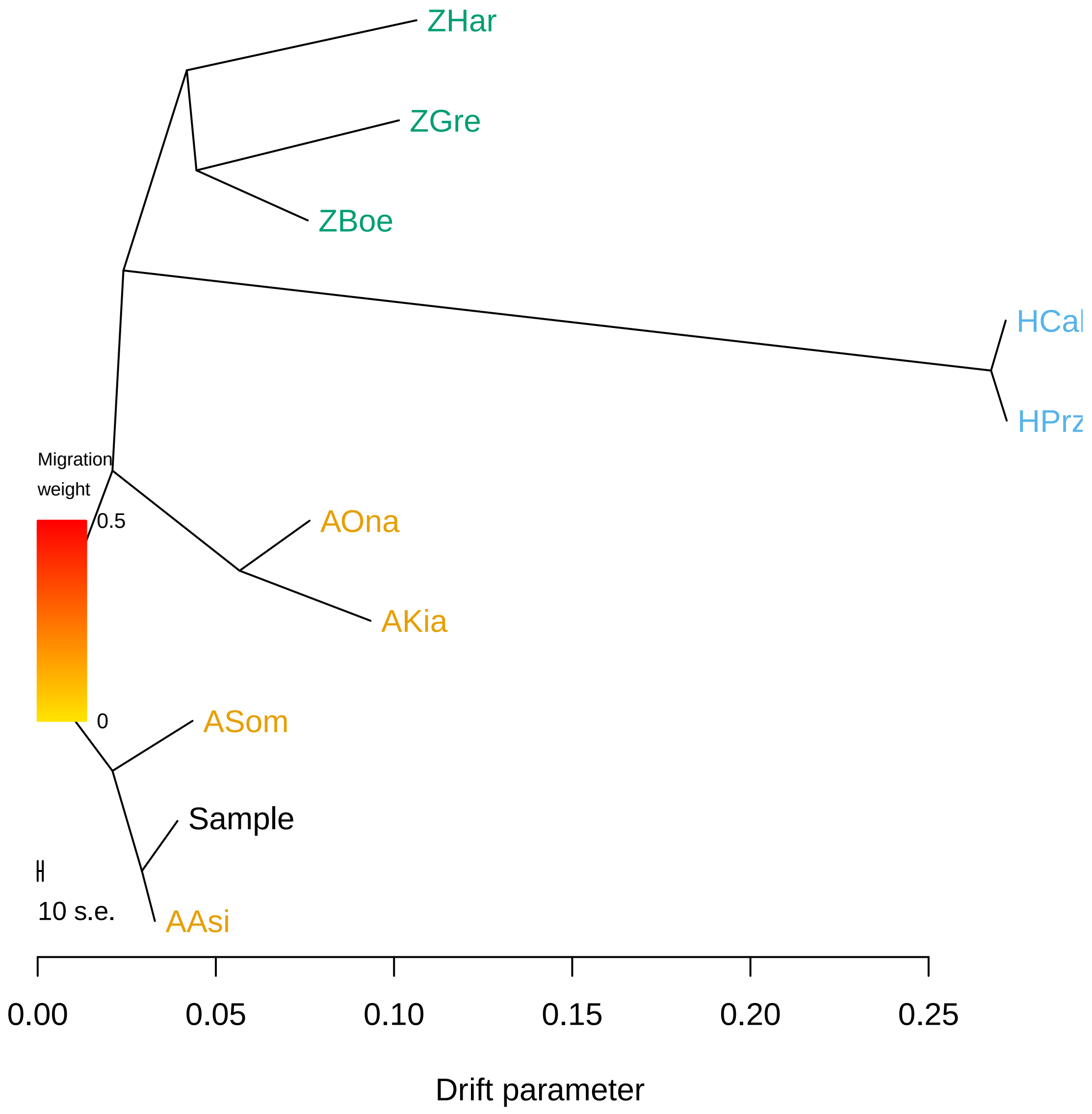

### incl_ts_0_tree.pdf

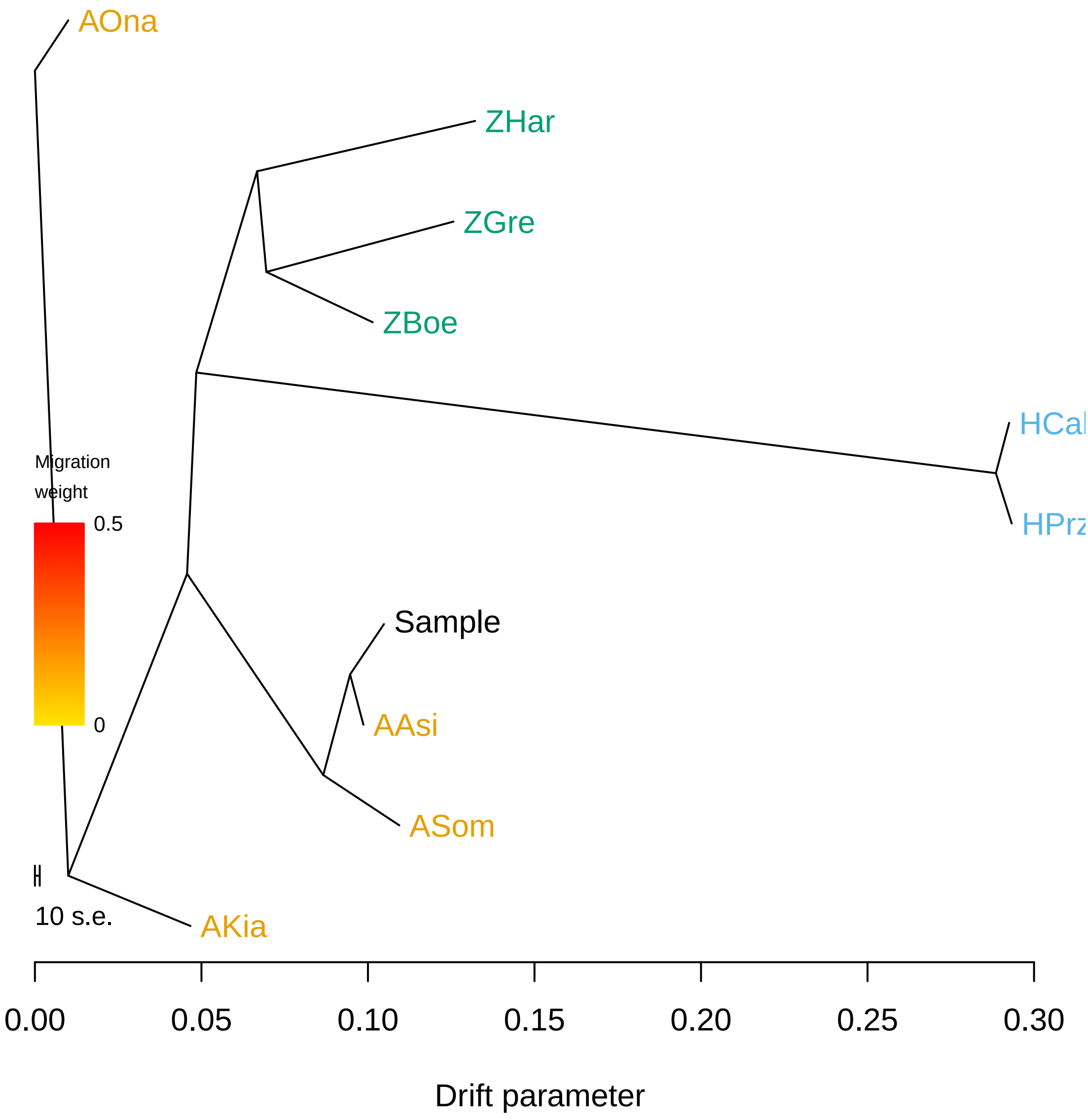

### incl_ts_0_tree.png

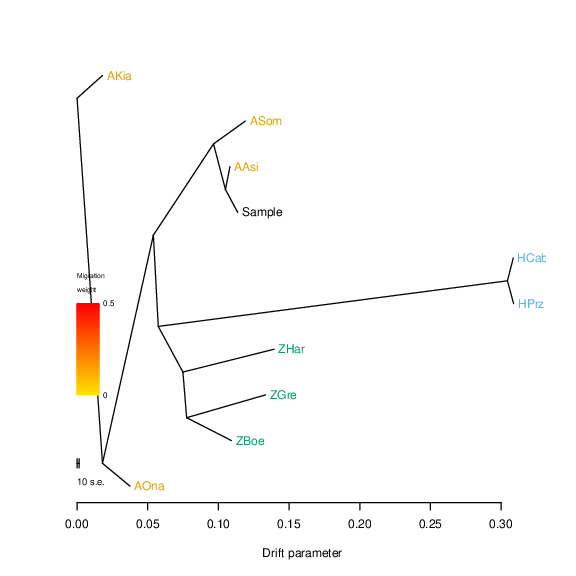

### incl_ts_0_tree.png

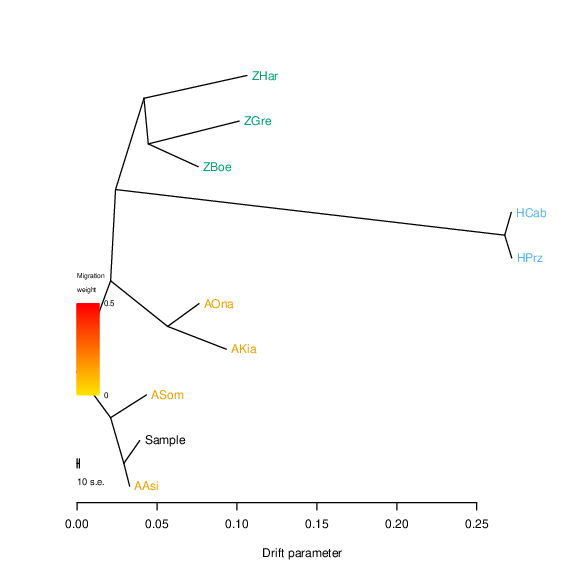

### incl_ts_0_tree.png

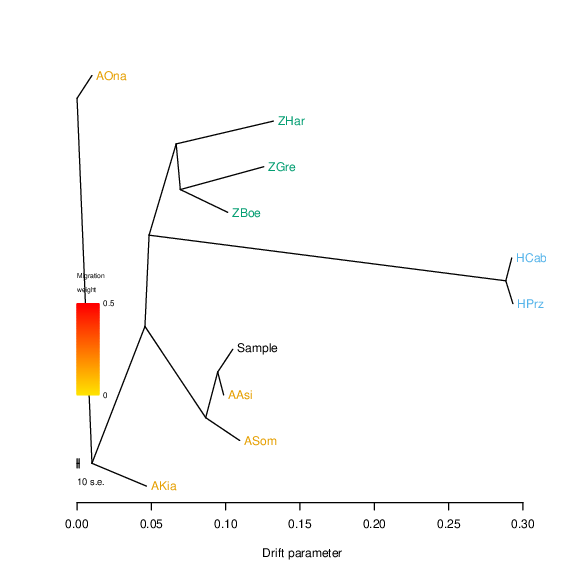

### incl_ts_1_residuals.pdf

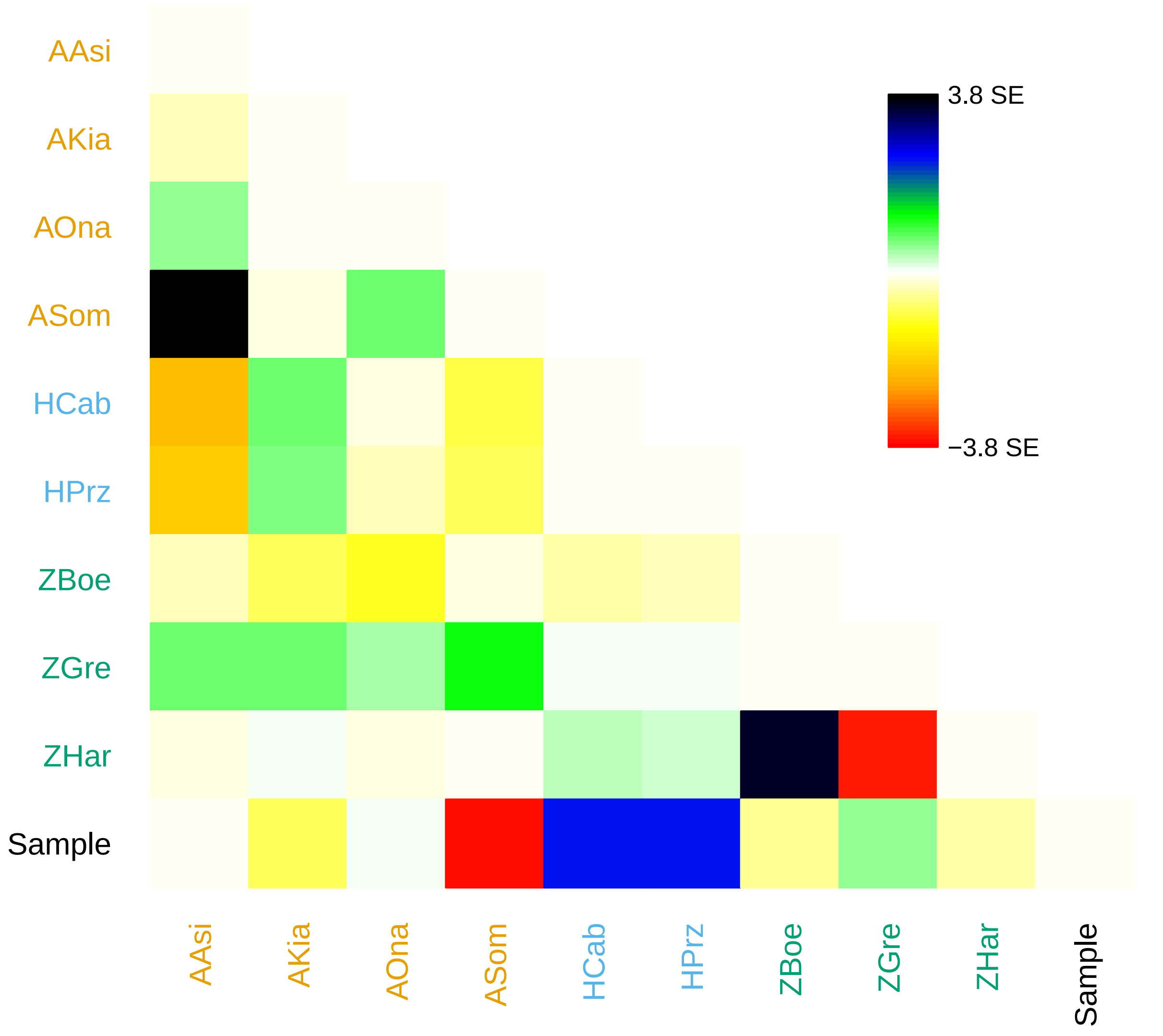

### incl_ts_1_residuals.pdf

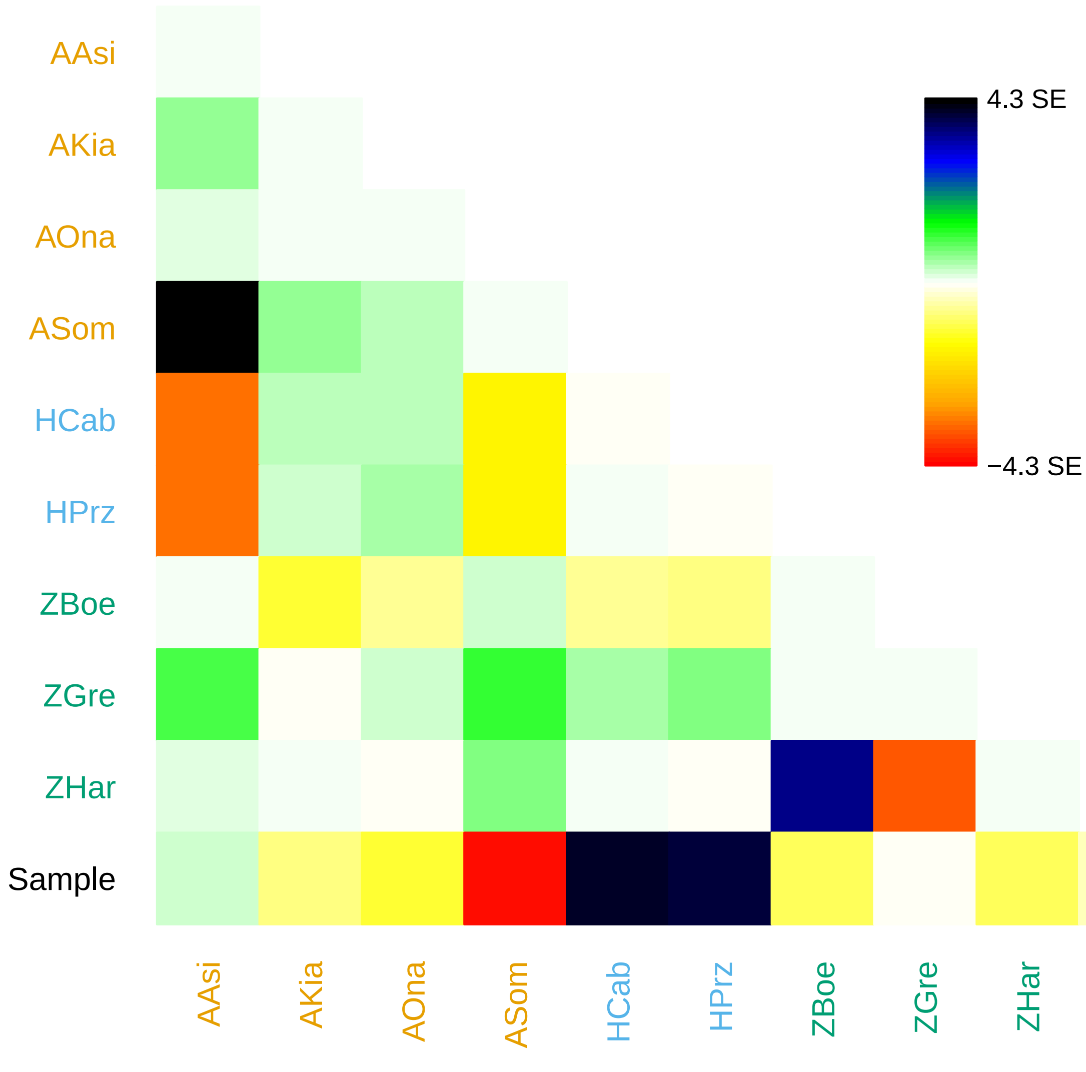

### incl_ts_1_residuals.pdf

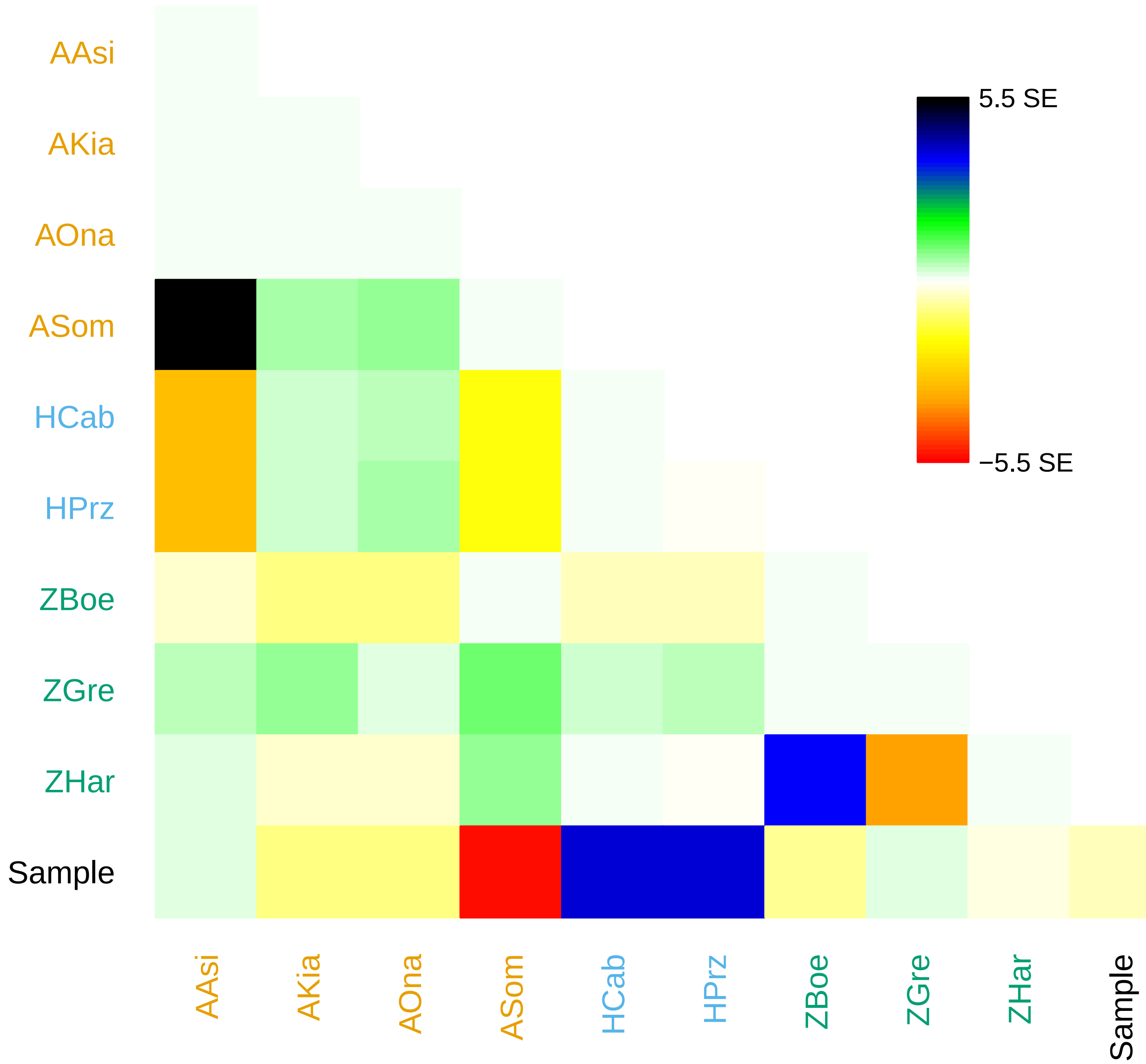

### incl_ts_1_residuals.png

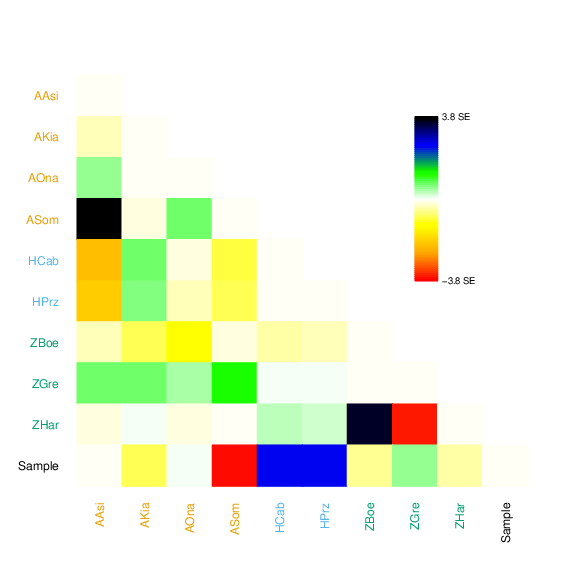

### incl_ts_1_residuals.png

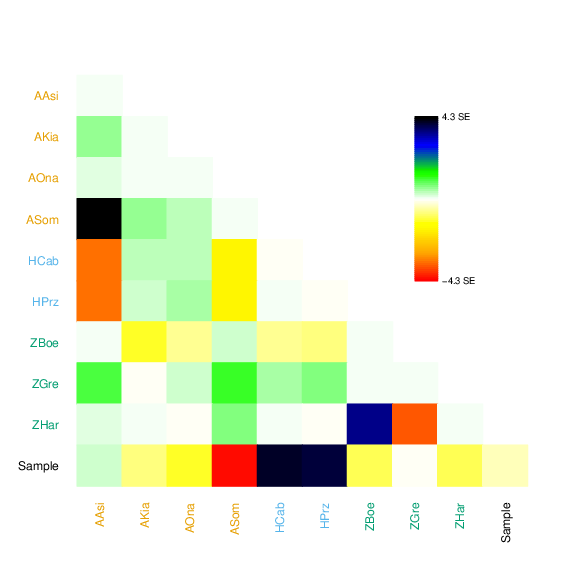

### incl_ts_1_residuals.png

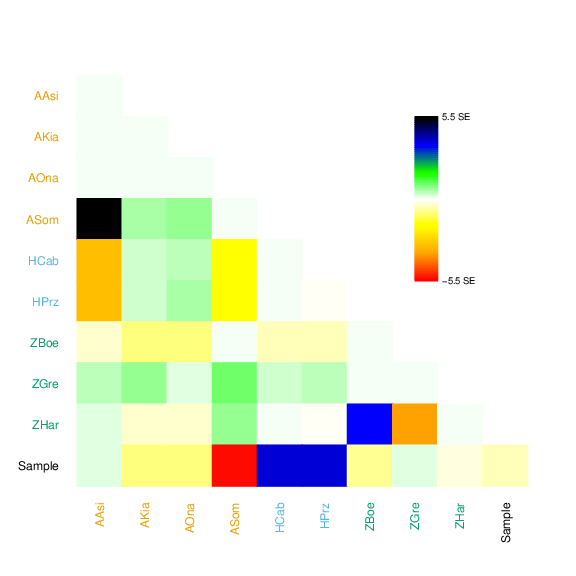

### incl_ts_1_tree.pdf

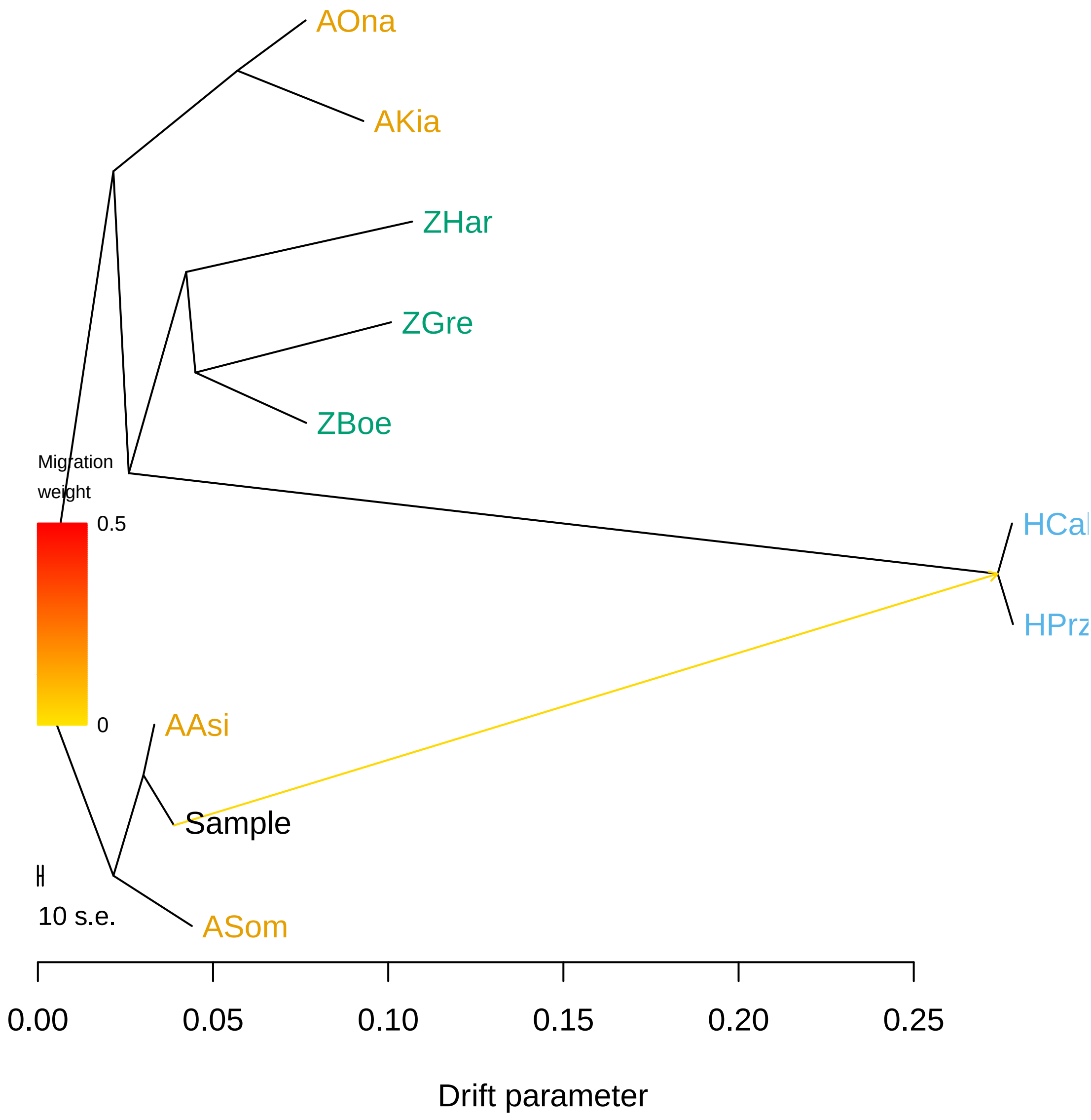

### incl_ts_1_tree.pdf

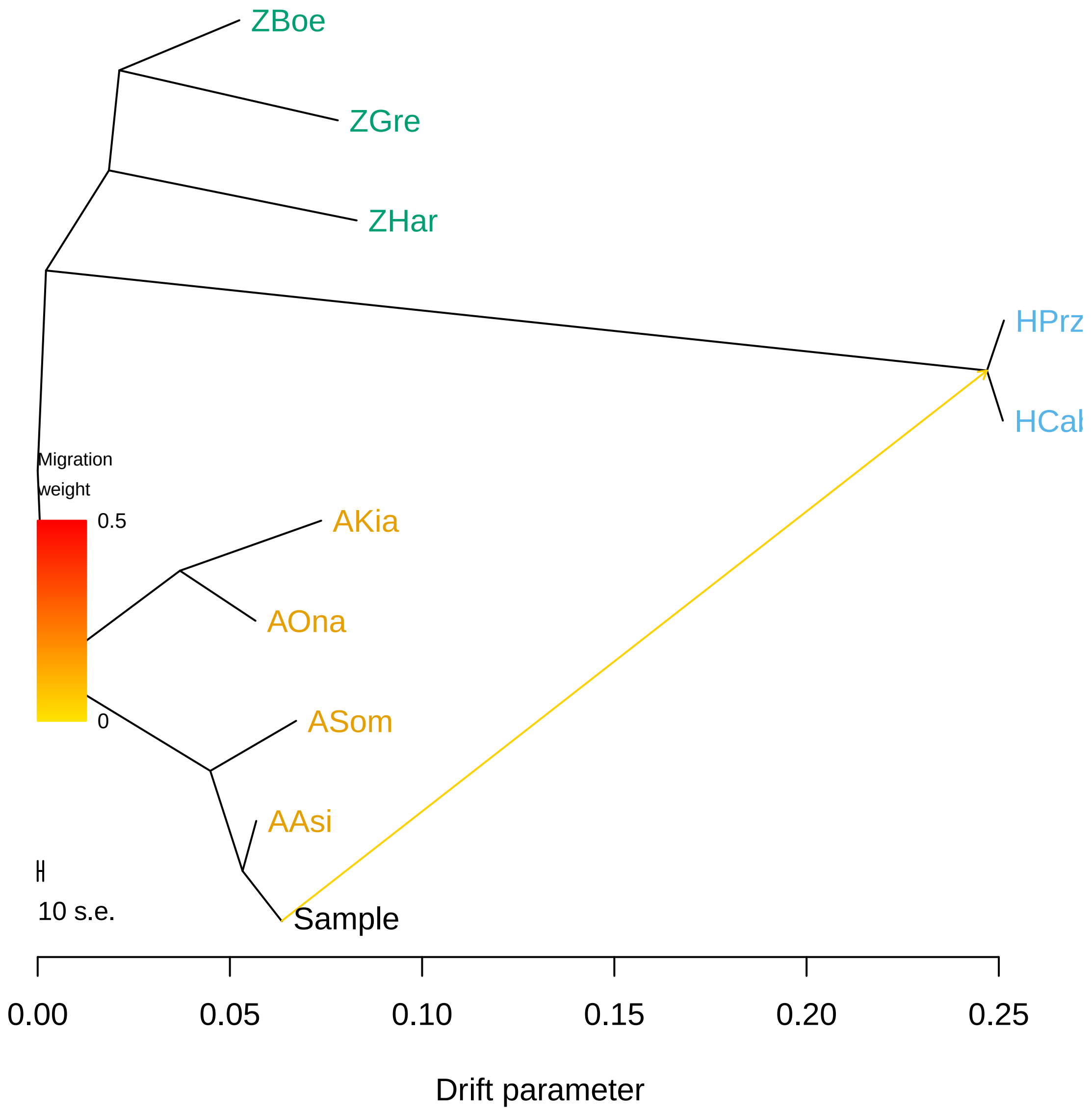

### incl_ts_1_tree.pdf

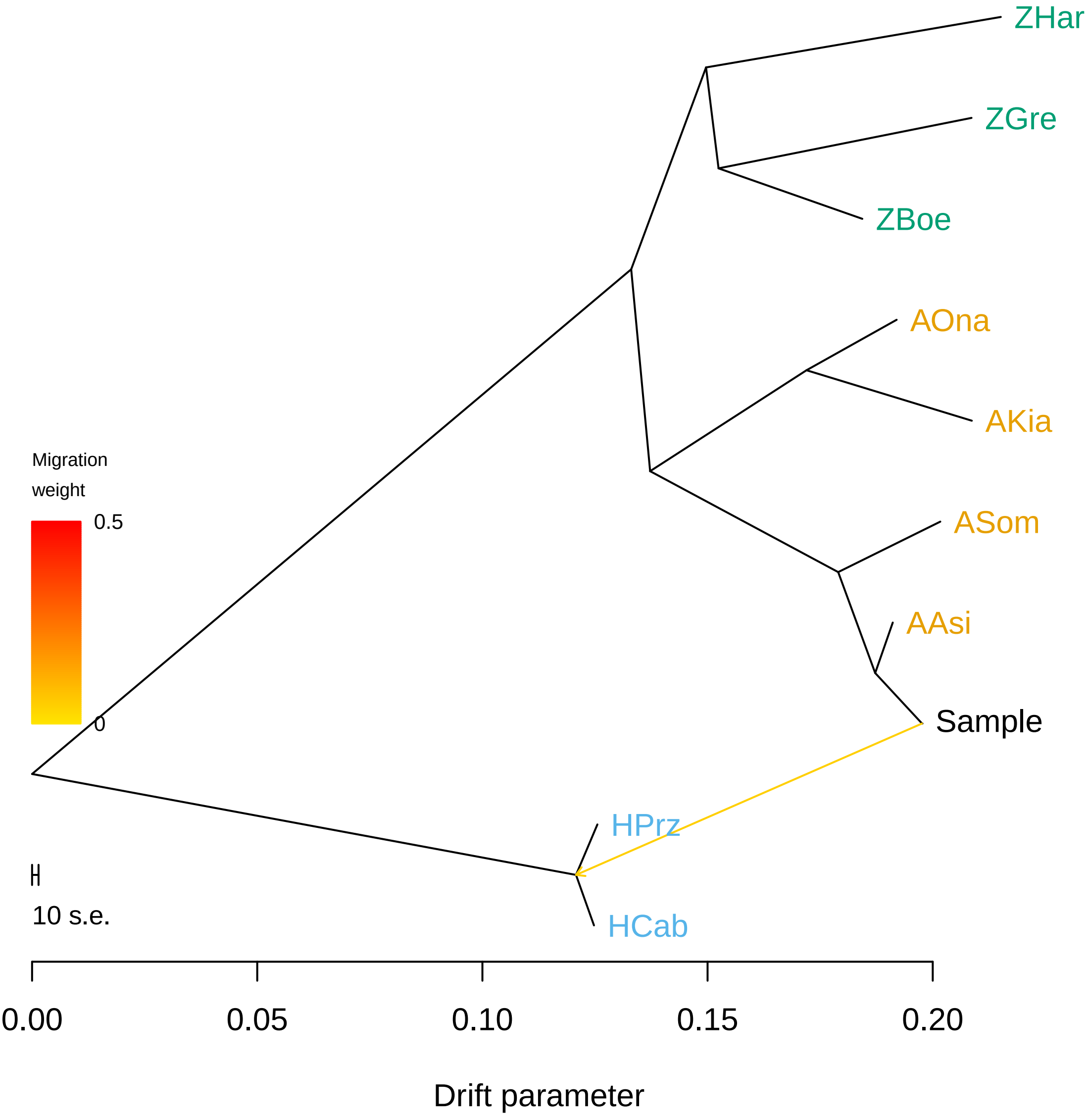

### incl_ts_1_tree.png

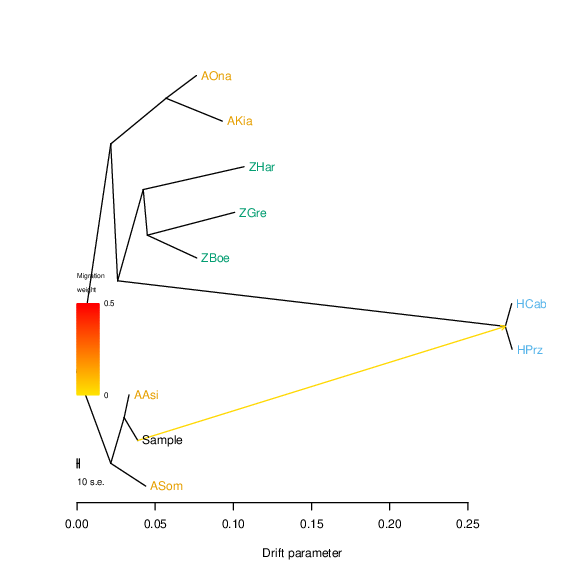

### incl_ts_1_tree.png

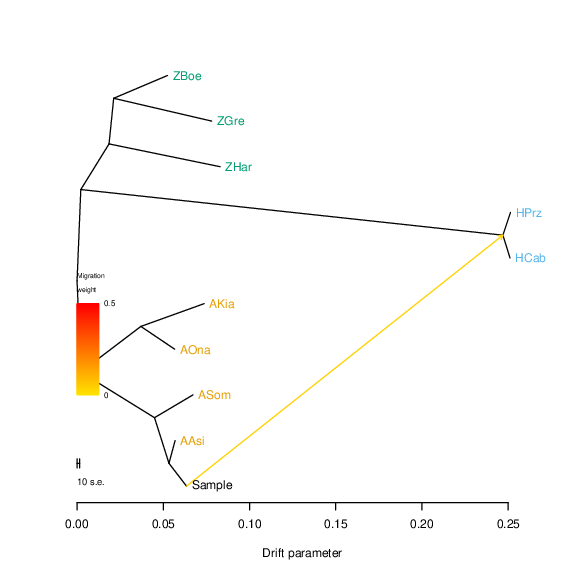

### incl_ts_1_tree.png

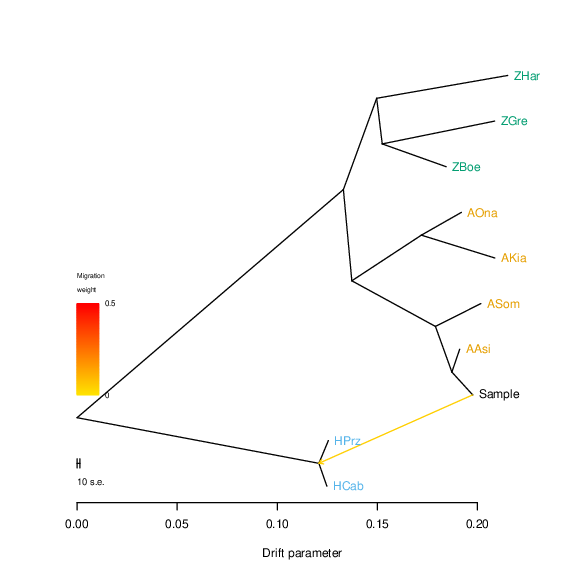

### incl_ts_k2.pdf

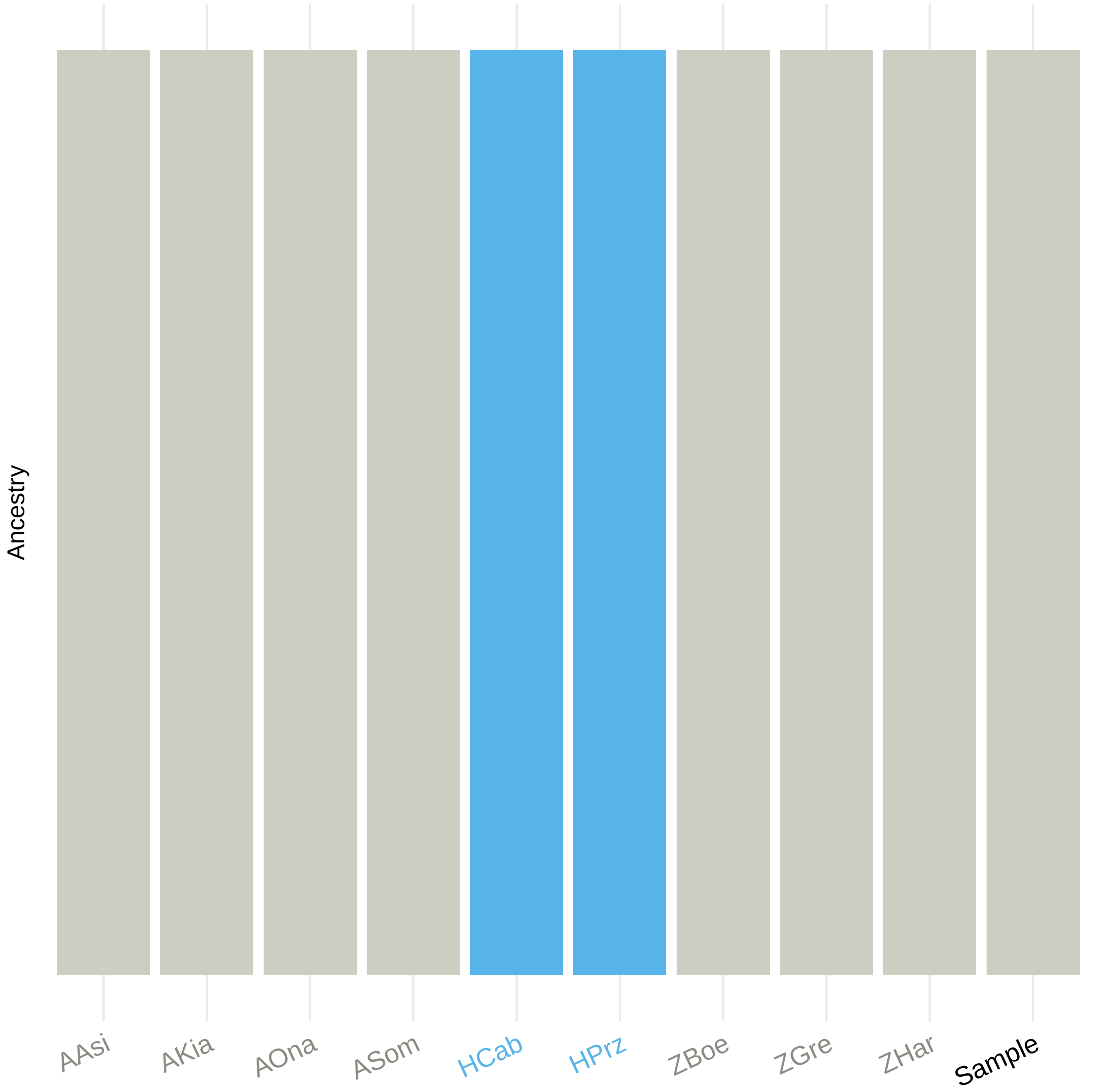

### incl_ts_k2.pdf

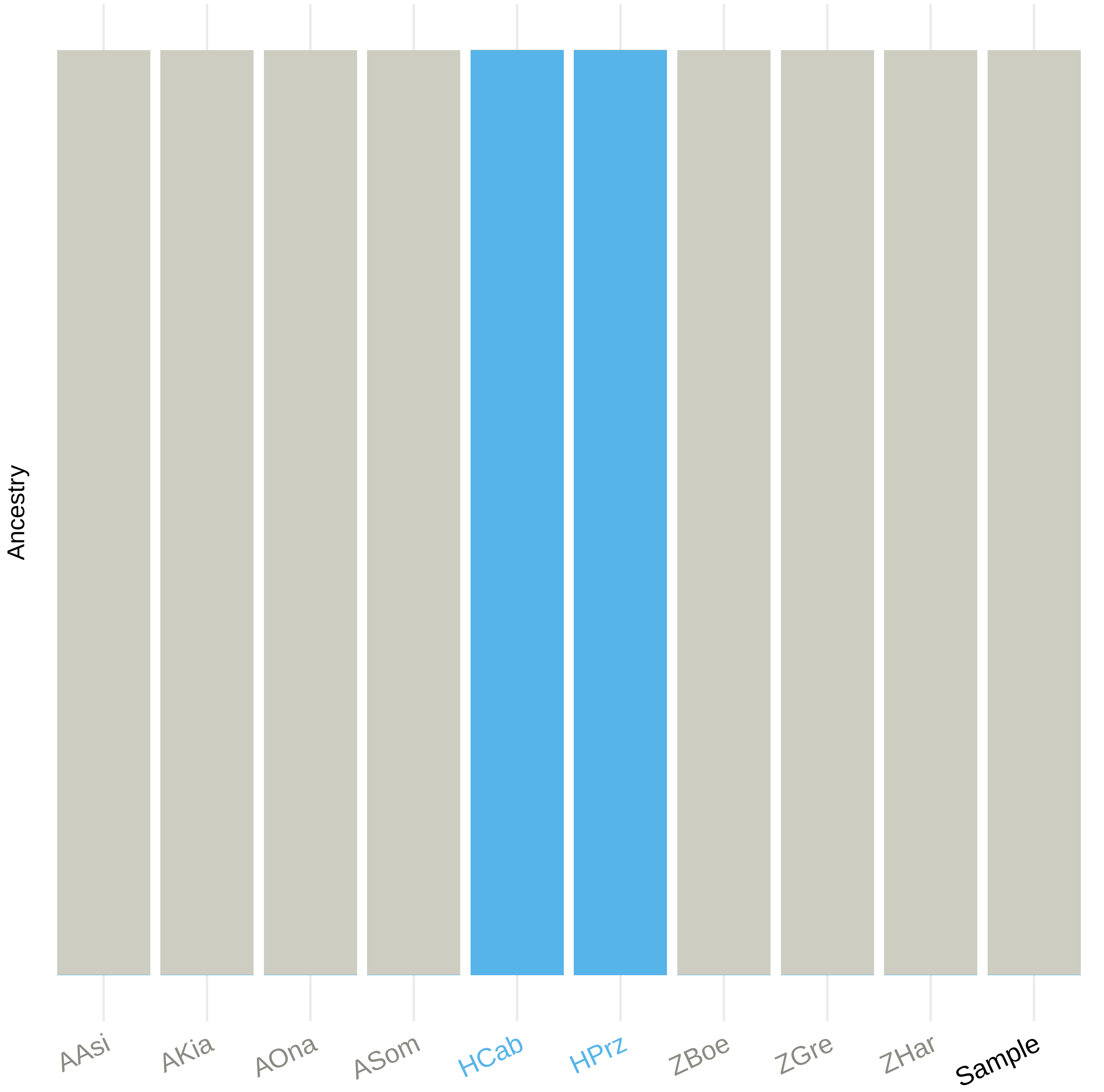

### incl_ts_k2.pdf

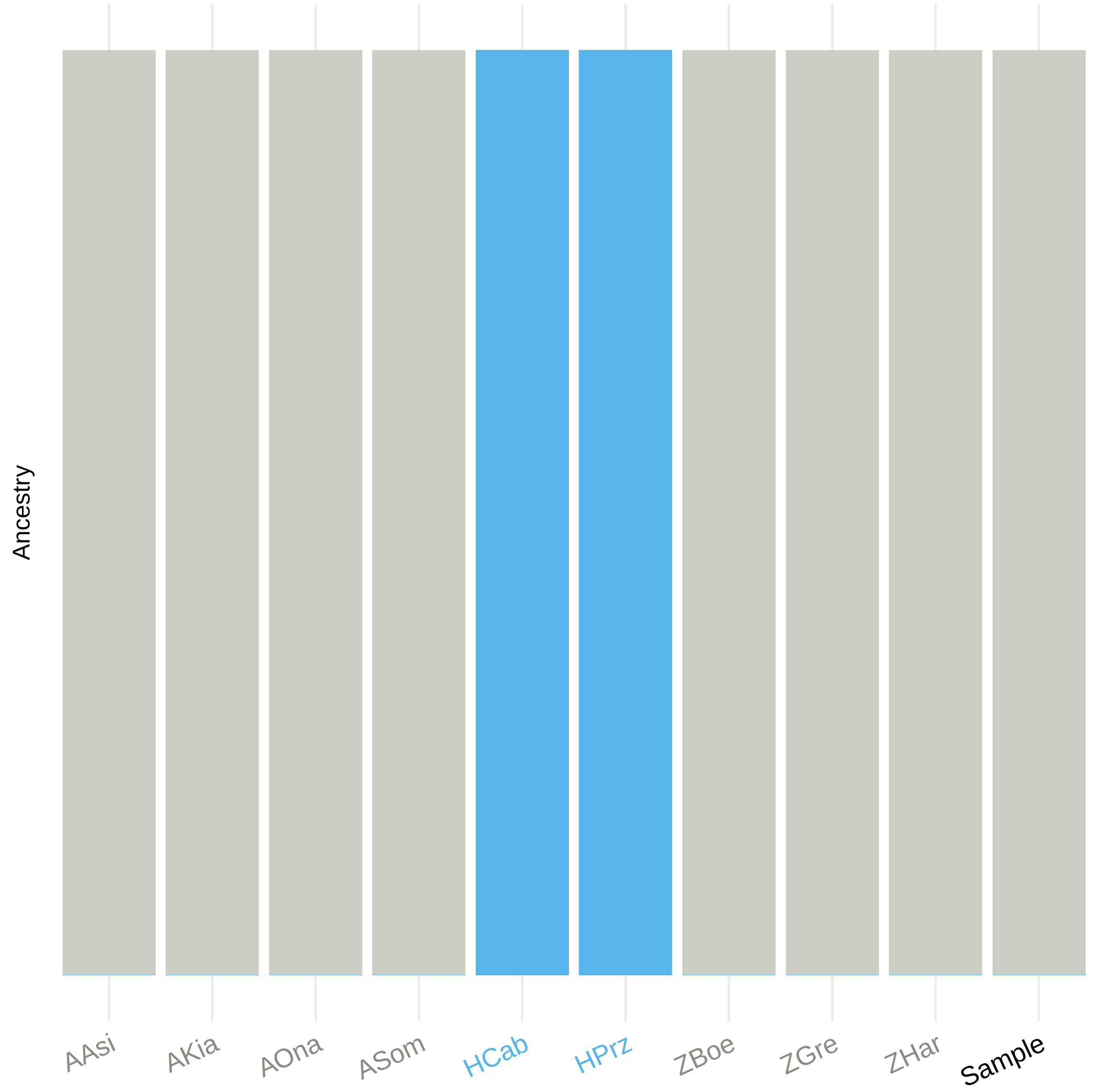

### incl_ts_k3.pdf

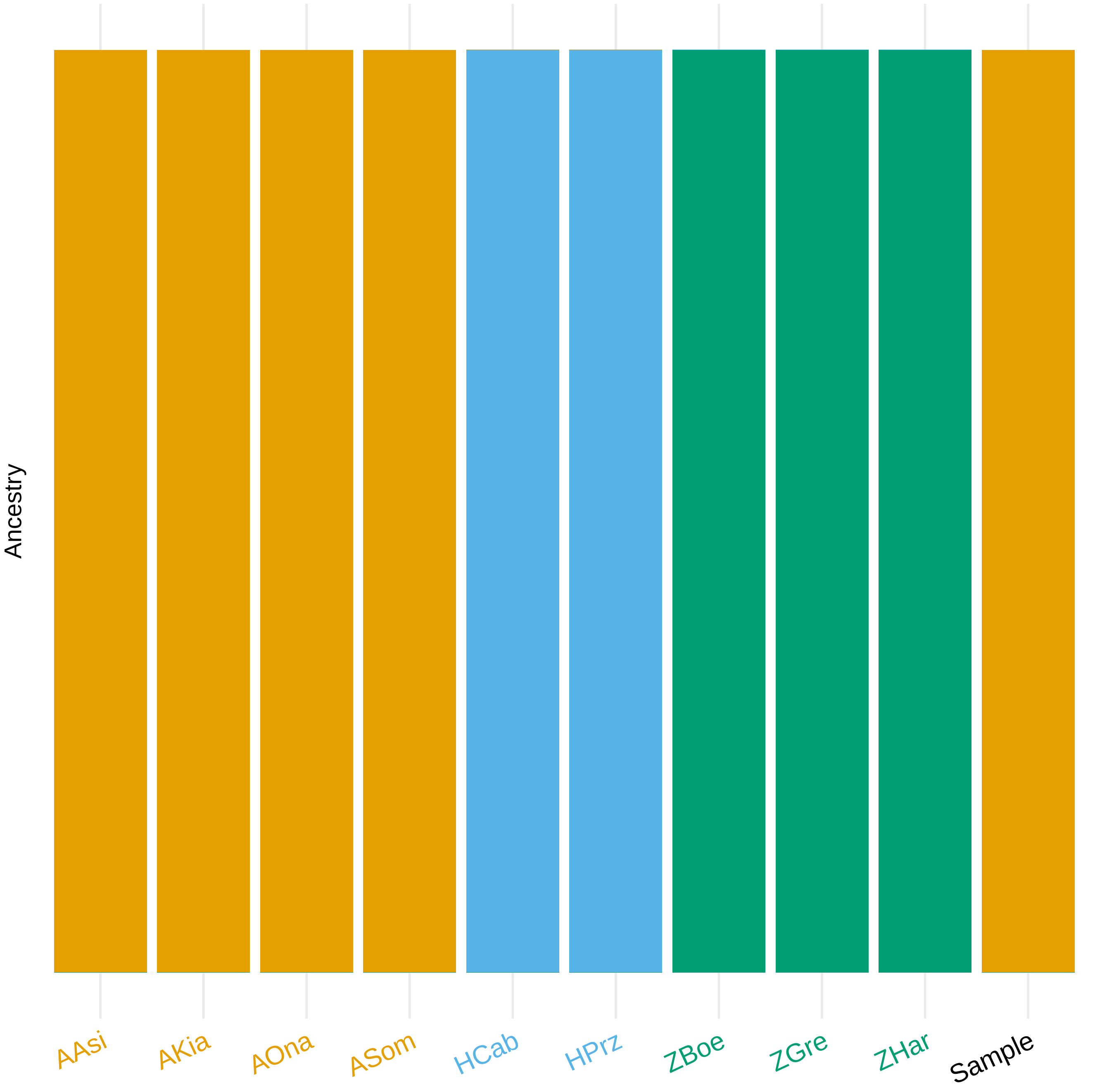

### incl_ts_k3.pdf

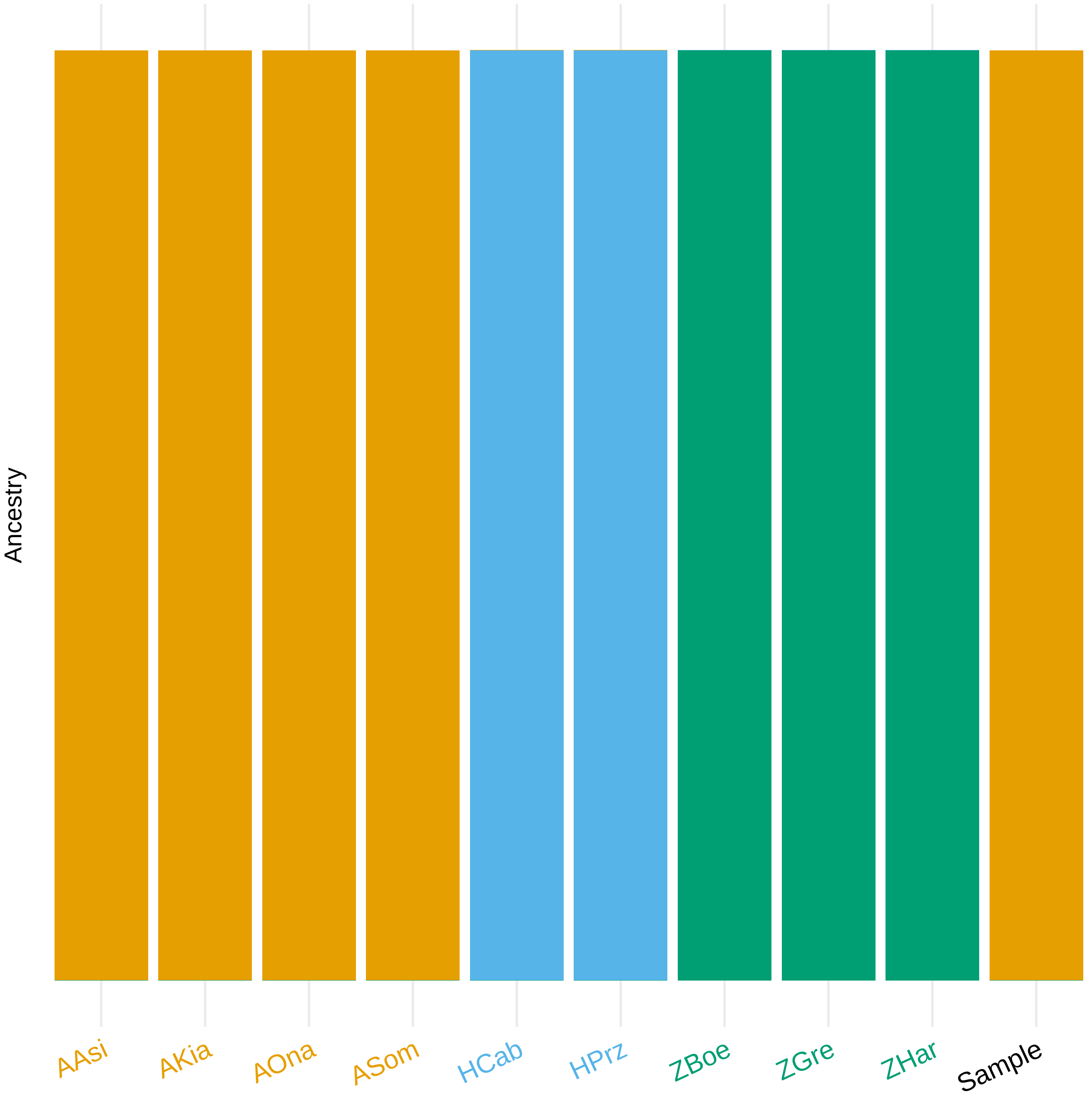

### incl_ts_k3.pdf

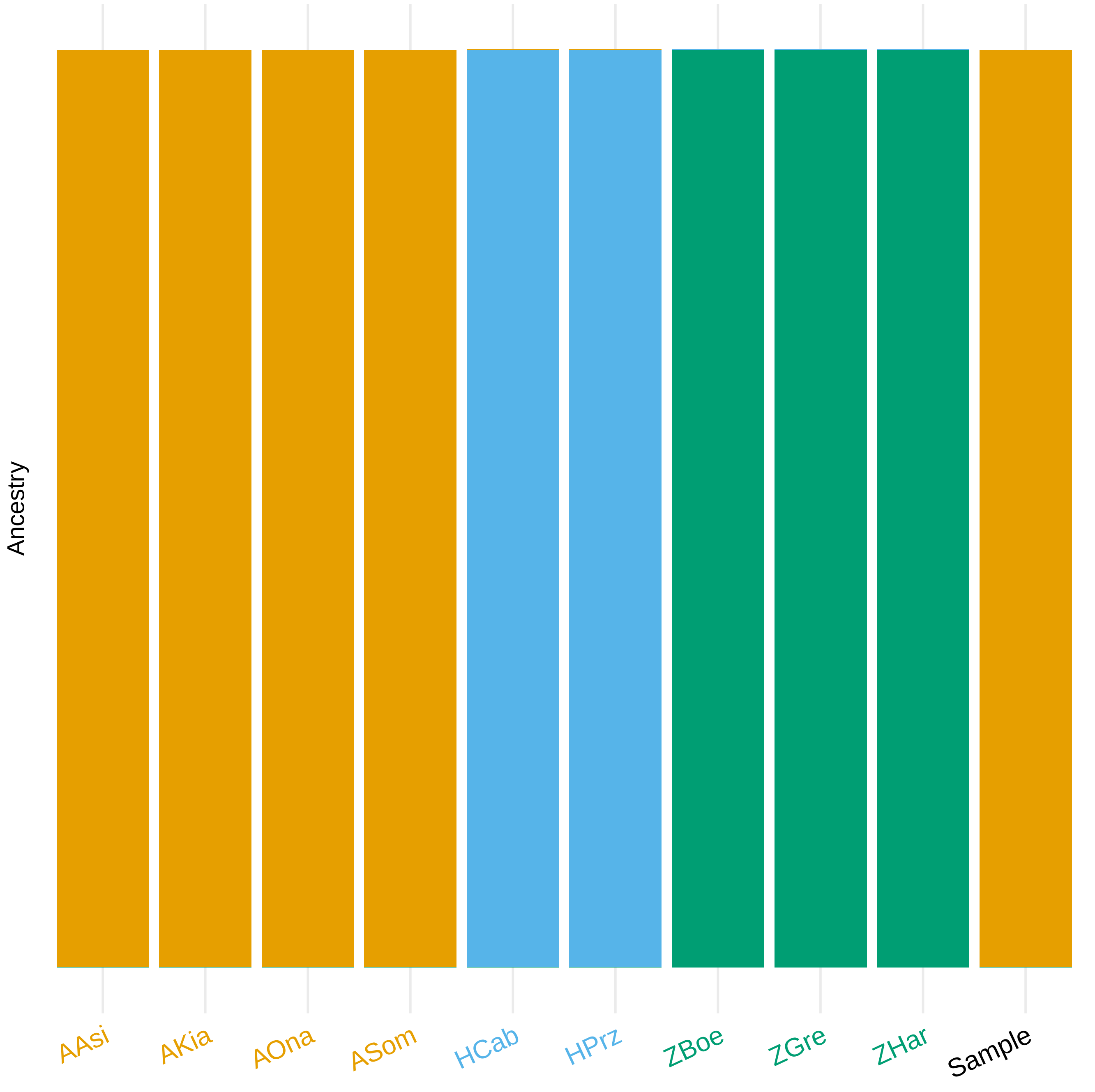

### mito_phylo.pdf

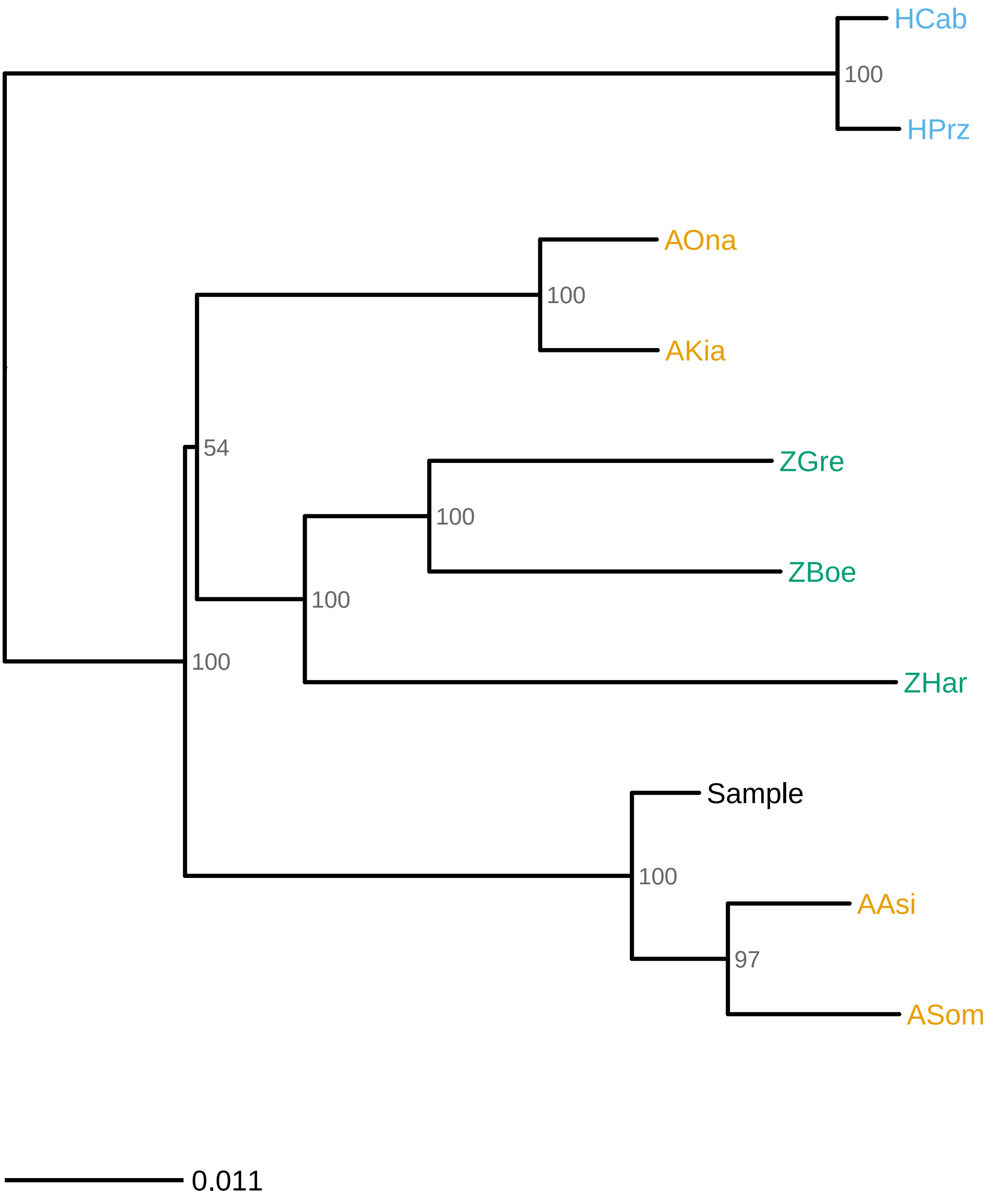

### mito_phylo.pdf

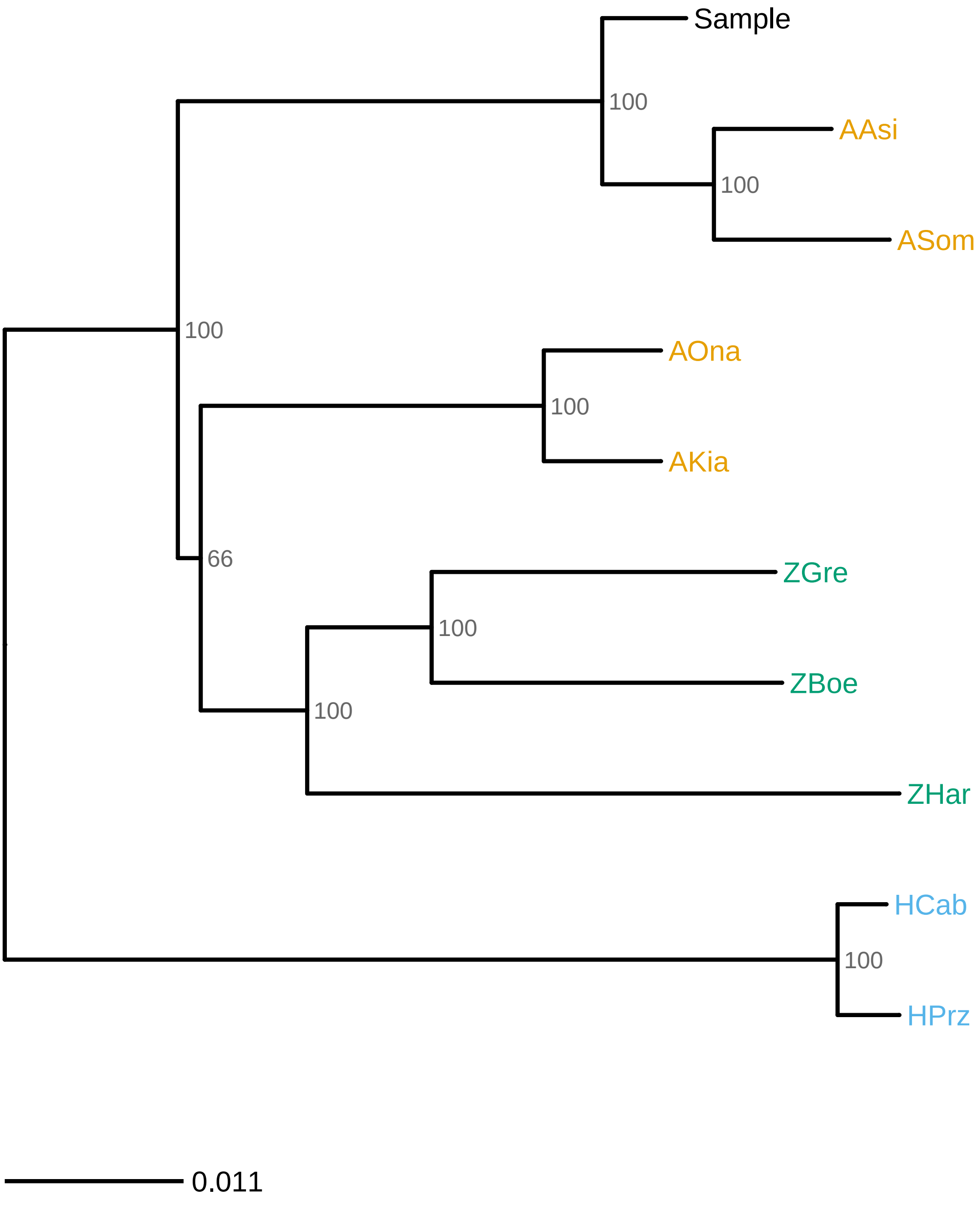

### mito_phylo.pdf

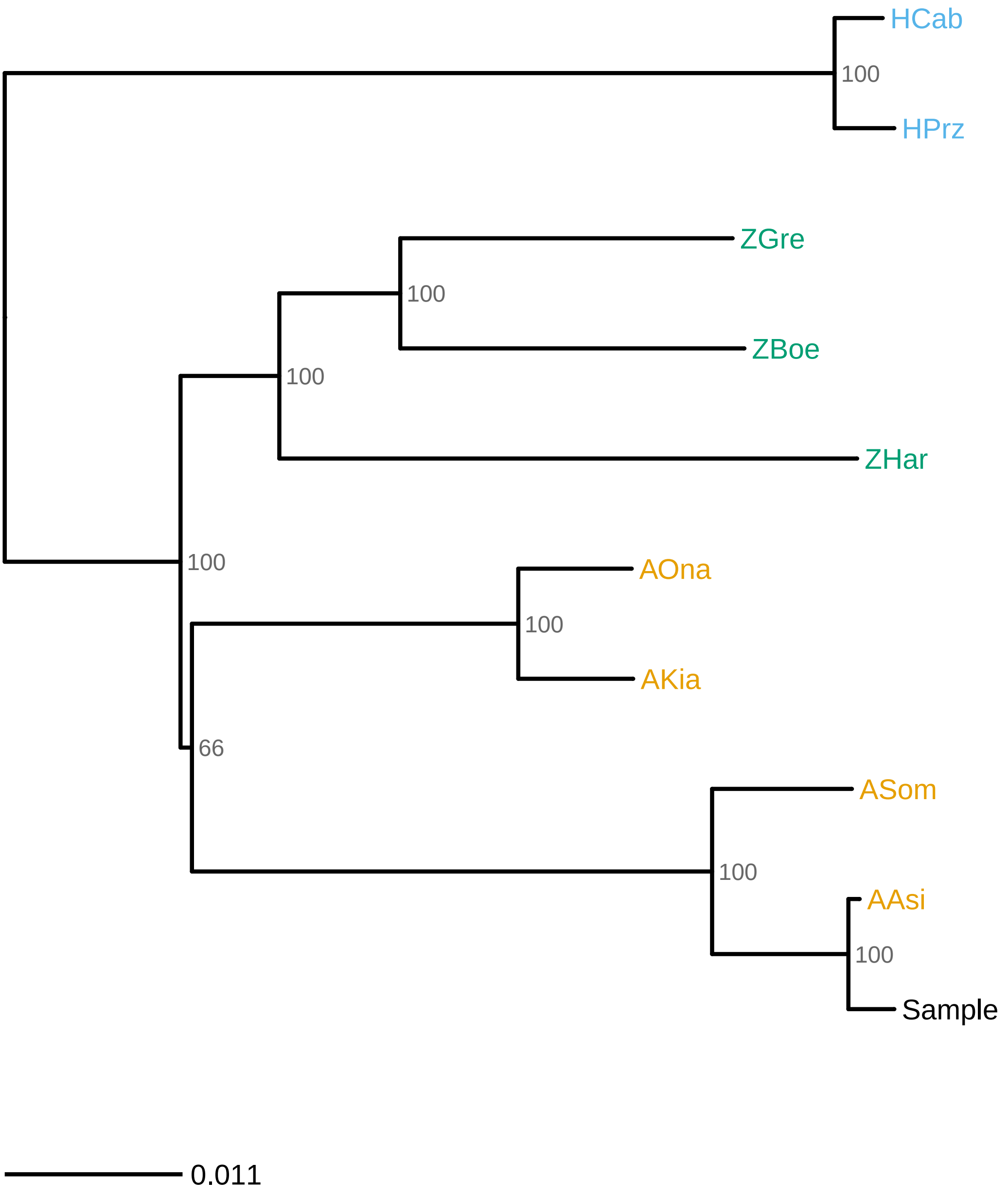

### mito_phylo.png

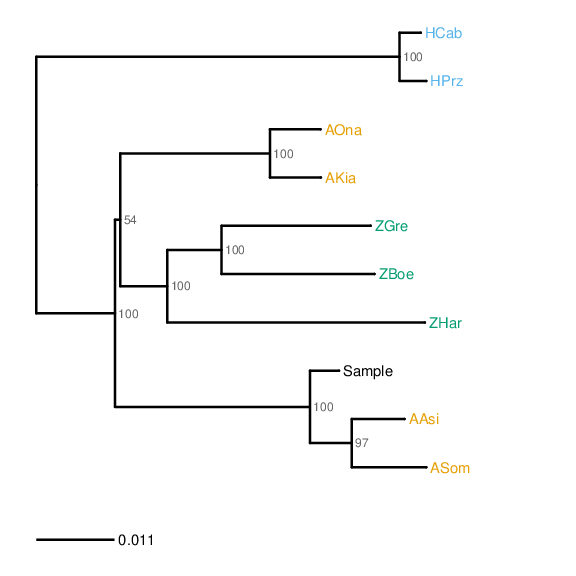

### mito_phylo.png

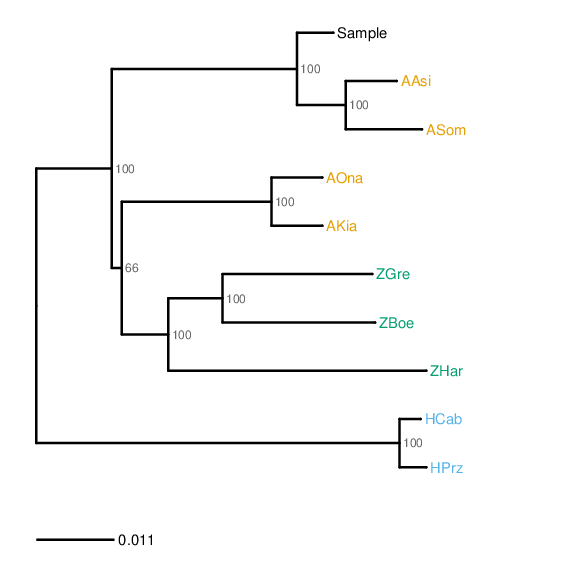

### mito_phylo.png

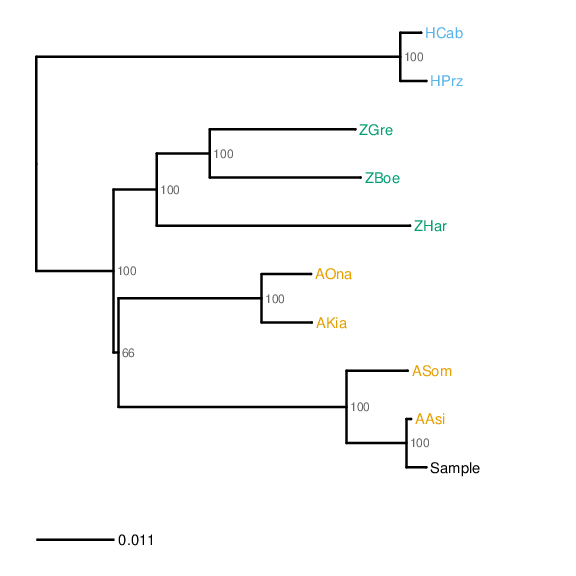
